# Characterization of adaptation mechanisms in sorghum using a multi-reference back-cross nested association mapping design and envirotyping

**DOI:** 10.1101/2023.03.11.532173

**Authors:** Vincent Garin, Chiaka Diallo, Mohamed Lamine Tekete, Korotimi Thera, Baptiste Guitton, Karim Dagno, Abdoulaye G. Diallo, Mamoutou Kouressy, Willmar Leiser, Fred Rattunde, Ibrahima Sissoko, Aboubacar Toure, Baloua Nebie, Moussa Samake, Jana Kholova, Julien Frouin, David Pot, Michel Vaksmann, Eva Weltzien, Niaba Teme, Jean-Francois Rami

## Abstract

The identification of haplotypes influencing traits of agronomic interest, with well-defined effects across environments, is of key importance to develop varieties adapted to their context of use. It requires advanced crossing schemes, multi-environment characterization and relevant statistical tools. Here we present a sorghum multi-reference back-cross nested association mapping (BCNAM) population composed of 3901 lines produced by crossing 24 diverse parents to three elite parents from West and Central Africa (WCA-BCNAM). The population was characterized in environments contrasting for photoperiod, rainfall, temperature, and soil fertility. To analyse this multi-parental and multi-environment design, we developed a new methodology for QTL detection and parental effect estimation. In addition, envirotyping data were mobilized to determine the influence of specific environmental covariables on the genetic effects, which allowed spatial projections of the QTL effects. We mobilized this strategy to analyse the genetic architecture of flowering time and plant height, which represent key adaptation mechanisms in environments like West Africa. Our results allowed a better characterisation of well-known genomic regions influencing flowering time concerning their response to photoperiod with Ma6 and Ma1 being photoperiod sensitive and candidate gene Elf3 being insensitive. We also accessed a better understanding of plant height genetic determinism with the combined effects of phenology dependent (Ma6) and independent (qHT7.1 and Dw3) genomic regions. Therefore, we argue that the WCA-BCNAM constitutes a key genetic resource to feed breeding programs in relevant elite parental lines and develop climate-smart varieties.

## Introduction

The quantitative nature of complex traits and their context specific expression are major hindrances for marker assisted selection (MAS) (Bernardo 2016; Cobb et al., 2019). The genotype by environment (GxE) effect is particularly problematic for MAS because it can strongly reduce or even reverse the QTL effect (Malosetti et al., 2013). However, the combination of advanced genetic resources, improved statistical methodology, and envirotyping data give us a chance to improve our understanding of the QTL by environment (QTLxE) effects. This understanding should increase our capacity to mobilize those effects for MAS and design varieties able to take advantage of specific environmental conditions.

### A. The BCNAM design and its properties

Multiparental populations (MPPs) combining the genomes of several founders have progressively emerged as central genetic resources for research (Scott et al., 2020; Bernardo 2021). The nested association mapping (NAM) design composed of crosses between a recurrent parent and donor parents is a well-spread MPP design (McMullen et al., 2009, Gage et al., 2020), with examples in maize (Bauer et al., 2013), rice (Fragoso et al., 2017), wheat (Kidane et al., 2019, Christopher et al., 2021) and sorghum (Bouchet et al., 2017). Sorghum is also the species that was used to develop the back-cross NAM (BCNAM) design, which consists of introgressing diverse alleles from donors in a recurrent (elite) line using one generation of back-cross followed by several generation of selfing (Jordan et al., 2011, Mace et al., 2021). BCNAM designs allow the introgression of diverse alleles in elite background while limiting the risk of introgressing deleterious alleles by keeping around 75% of the elite genome. It can serve research purposes for genetic analysis and breeding purposes (Scott et al., 2020).

BCNAM design has several interesting properties for genetic analyses. Compared to bi-parental crosses it addresses a larger genetic diversity and captures more recombination events. Compared to association panels, it offers better control over the population structure, which can reduce the detection of false positive signals (Myles et al., 2009). BCNAM designs also allow to trace back the origin of favourable alleles to a specific parent, a highly desirable feature to design future crosses. MPPs like BCNAM increase the rare allele frequencies, which is essential to precisely estimate their additive effects (Myles et al., 2009). Moreover, the possibility to extend the reference NAM design by using several recurrent parents allows the characterization of the genetic effect in multiple genetic backgrounds (Christopher et al., 2021). Finally, BCNAM designs are also interesting for GxE studies because it can be used to measure the expression of an interconnected set of diverse alleles in contrasting environments (Cobb et al., 2019).

### B. Improved statistical methods for QTL detection in MPP designs

Several approaches have been developed to detect QTL in MPPs characterized in a single environment. For example, Garin et al. (2017, 2018) proposed a framework assuming different allelic configurations at the QTL position. Li et al. (2011) developed a method based on maximum likelihood parental allelic effect significance. Xavier et al. (2015) used mixed models employed for genome-wide association analysis accounting for crossing structure. More recently Paccapelo et al. (2022), adapted the whole genome interval mapping method for the NAM design. A more general strategy consists of using models with identical by descent probability that can be estimated for any type of MPP designs (Zheng et al., 2015; Li et al., 2021).

Compared to separate within-environment analyses, the QTL detection using MPP data characterized in multiple environments (MPP-ME) in a joint model is more challenging, but it allows a more direct comparison of the simultaneously estimated effects. Until now, phenotypic values were averaged across environments (e.g. Giraud et al., 2014), which does not use the full potential of those data. Therefore, Garin et al. (2020) extended the MPP-ME QTL detection methodology using joint analyses. Diouf et al. (2020) proposed a forward-backward algorithm for MPP-ME analysis. De Walsh et al. (2022) proposed a meta-analysis of single environments analyses. Those MPP-ME models are particularly useful to characterize the trait variability in terms of genetic (parents, genetic background) and non-genetic (environment) effects.

### C. Extending genetic modelling with envirotyping

The recent progresses in sensor technologies have considerably increased the availability of large-scale environmental information (Xu 2016, Costa-Neto et al., 2021). Therefore, in this study, we extended MPP-ME QTL detection models by integrating environmental covariables (ECs) to refine our understanding of the GxE interaction by testing the sensitivity of multiple parental alleles with respect to various ECs. Among the available ECs, photoperiod is a key variable for sorghum development, especially in West Africa. Photoperiodism is the developmental responses of plants to the relative length of daylight or photoperiod (Hopkins 2009). Sorghum is a short-day plant generally sensitive to photoperiod that flowers when days become shorter than a certain length (Wolabu and Tadege 2016). When day length is longer than the critical photoperiod, photoperiod sensitive sorghum delays its panicle initiation. The flowering time can be represented as a broken linear function of the photoperiod with a baseline duration remaining constant until a certain photoperiod then an increasing slope where flowering time increases with the photoperiod (Van Oosterom et al., 2001; Figure 4G). The photoperiod sensitivity is the steepness of the slope. Since adaptation of sorghum to its cultivation site relies largely on its photoperiod sensitivity it is of paramount importance to integrate this environmental dimension in our analysis.

In this article, we present a multi-reference sorghum back-cross nested association mapping populations composed of 24 diverse parents anchored on three West African elite lines that represents one of the most relevant publicly available resources for West and Central Africa sorghum (WCA-BCNAM, Table 2, Figure S1). The sub-populations were phenotyped for flag leaf appearance (FLAG), plant height (PH), number of internodes (NODE_N), average length of the internodes (NODE_L), peduncle length (PED), panicle length (PAN), 1000 grains weight (GWGH), and grain yield (YIELD) in multiple environments contrasting for sowing date (photoperiod), rainfall, temperature, and soil fertility over two seasons.

To analyse those data, we developed a methodology for MPP-ME QTL analysis integrating environmental covariables (Figure 1). We illustrate our approach through a fine characterization of major QTL for flowering and plant height and discuss how the combination of advanced genetic resources and statistical methodology can support the design of climate-smart varieties.

**Figure 1.**
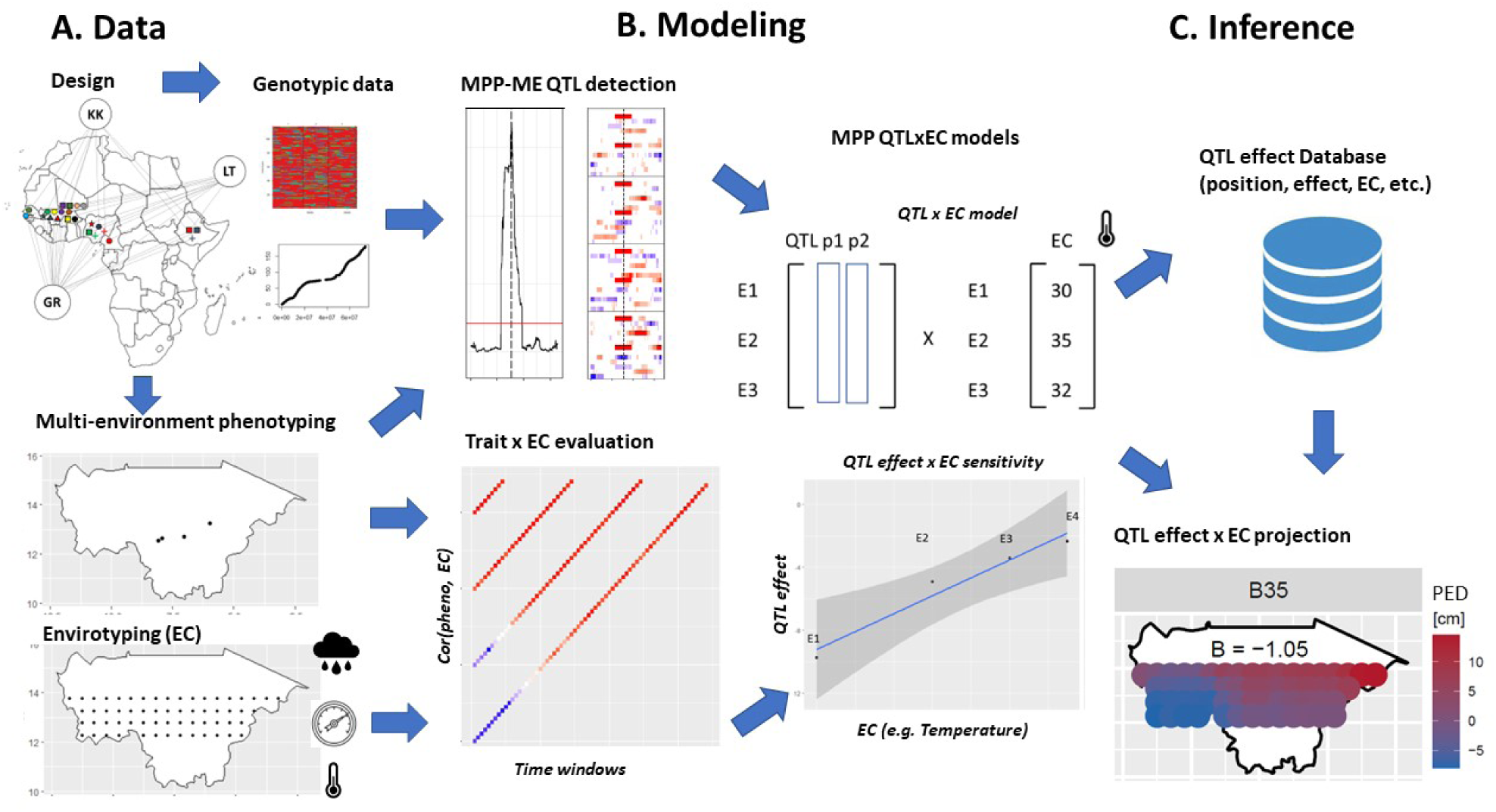
Overview of the analytical strategy. A) Raw genotypic phenotypic and environmental covariable; B) Statistical models for QTL detection in MPP characterized in multiple environment (ME), correlation between trait and environmental covariables (EC) analysis, and synthesis in QTLxEC models; C) Inference using the results gathered in a database and projection of the QTL effect beyond the tested environments.

## Results

### D. Genetic diversity

Figure 2A illustrates the genetic diversity covered by the parental lines of the WCA-BCNAM population compared to a panel representative of the global sorghum diversity (Methods S1). We compared the number of common polymorphic SNPs of our population, of the US-NAM (Bouchet et al. 2017) and the global diversity panel. Overall, the WCA-BCNAM parents covered 90.2% of the global sorghum genetic diversity considered for the analysis (192K SNP) which offer a slightly better coverage (11 %) than the parents from the sorghum US-NAM which already captured 79.2% of the considered diversity. Principal component analysis of the WCA-BCNAM genetic data (Figures 2 B, C and D) detected three distinct groups corresponding to the three recurrent parents (Figure S2). Clear subdivisions of the populations according to the donor parent race and some specific divergences from this general pattern were observed especially for the populations involving the Hafijeka Guinea Margaritiferum accession.

**Figure 2.**
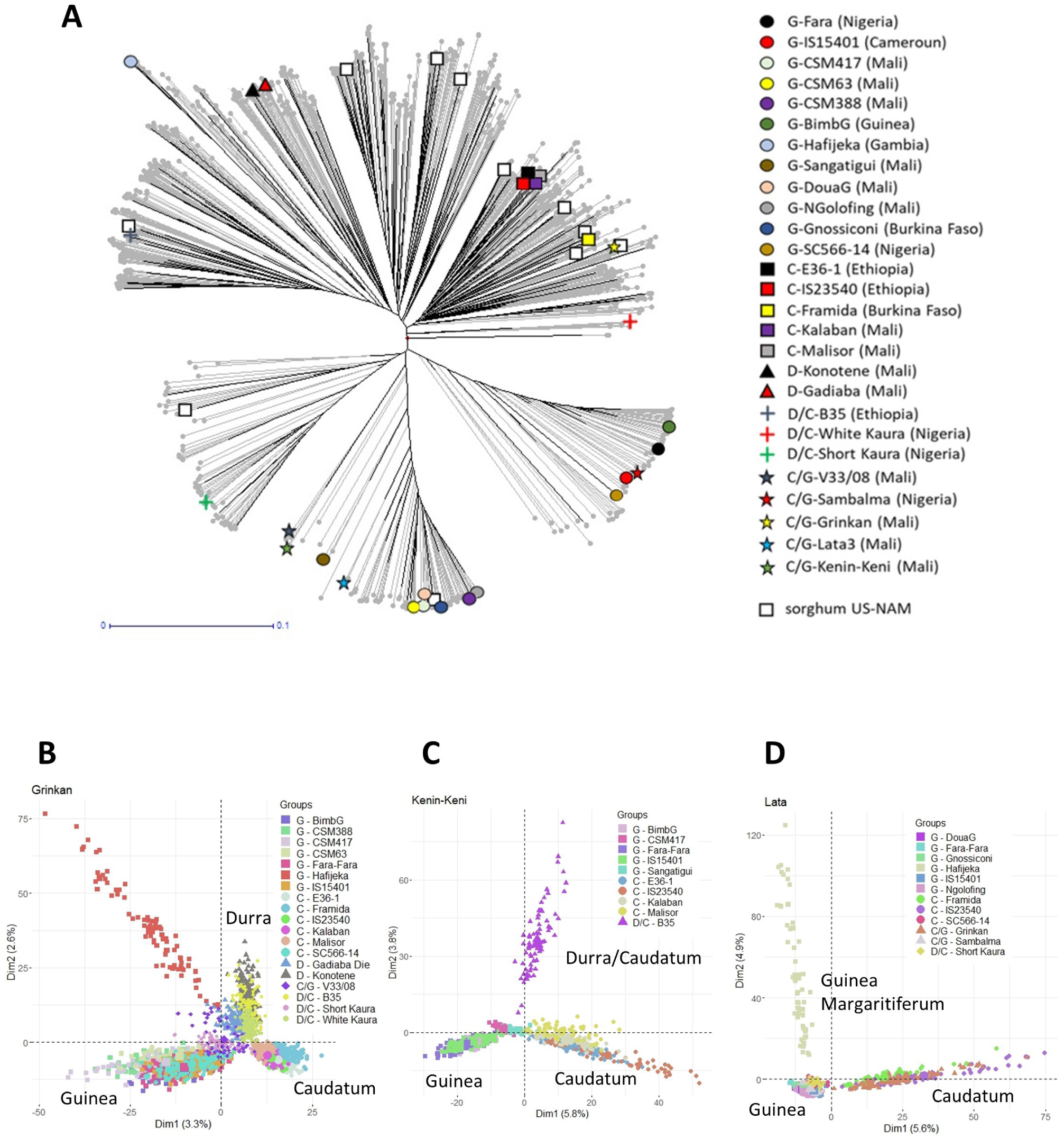
Genetic diversity and structure of the WCA-BCNAM design. A) Coverages of the global sorghum molecular diversity by the WCA-BCNAM and sorghum US-NAM (white square, Bouchet et al. 2017) parents. Principal component bi-plots performed on a subset of 5000 markers randomly selected of the (B) Grinkan (C), Kenin-Keni (D) Lata3 sub-populations.

**Figure 3.**
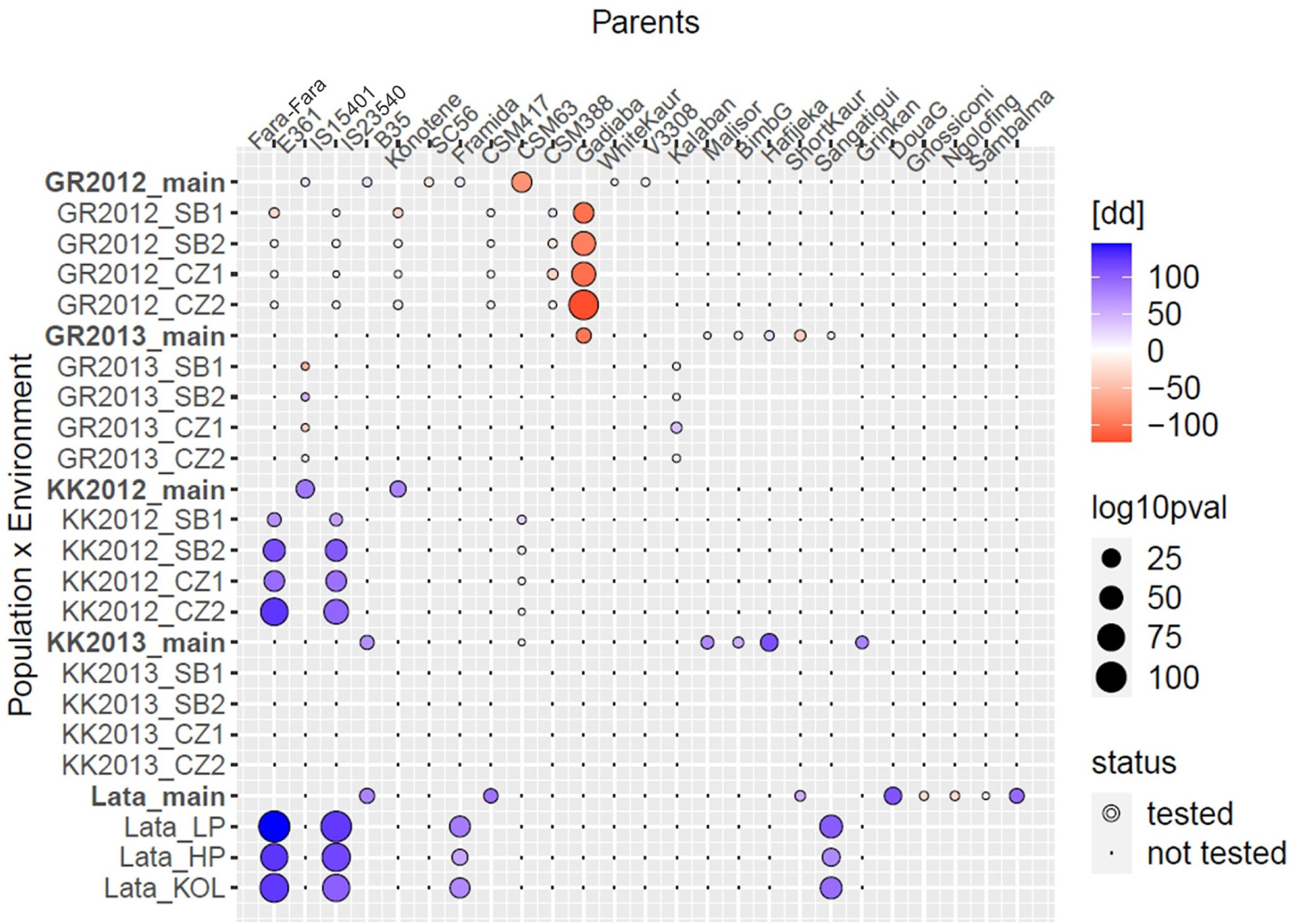
Estimated QTL allelic effect for flag leaf appearance at chromosome three position 77.3 cM for the donor parents (x axis) evaluated in different genetic background (Grinkan, Kenin-Keni, Lata3) and/or environments (SB1-2: Sotuba sowing 1 and 2, CZ1-2: Cinzana sowing 1 and 2, LP/HP: low/high phosphorus, KOL: Kolombada). The colour is proportional to the effect size and direction and the circle size to its significance. An empty space means that the main effect across environments is more significant than the QTLxE effects. Dots means that the parental allele effect was not evaluated in this specific background in this specific trial.

### E. Phenotypic data

The average heritability values over populations were larger for traits like FLAG (0.78-0.95), PH (0.76-0.88) or NODE_L (0.8-0.9) compared to YIELD (0.37-0.64) (Tables S3). Heritability values were larger in the Lata3 sub-population which is due to the within-environment replication as well as the larger similarity between the environmental conditions in which the Lata3 sub-population was phenotyped. In terms of correlation between traits (Figures S3 and S4), we observed an overall negative relationship between FLAG and YIELD. With an average Pearson correlation of -0.28 and a standard deviation of 0.17. This negative relationship was observed in all genetic backgrounds and environments but was stronger at the second sowing (-0.38 ±0.15). FLAG and NODE_N were positively correlated in all backgrounds (Grinkan, Kenin-Keni, and Lata3). This correlation was stronger at the second sowing time (S1: 0.34 ±0.15; S2: 0.45 ±0.15). Concerning the correlation of PH with its components, the strongest one was with NODE_L (0.74 ±0.12), the lowest with NODE_N (0.26 ±0.14). It took intermediary values for PED (0.53 ±0.15) and PAN (0.41 ±0.13). This pattern was observed in all configurations. PH was positively correlated with YIELD (0.23-0.56), except for the Grinkan sub-population measured in 2013 at Sotuba (-0.11 ±0.01). Looking at the correlation between PH, its components (NODE_L, NODE_N, PED, PAN), and YIELD, we noticed that it was undetermined with NODE_N (0.03 ±0.14), and positive with PED (0.17 ±0.18), PAN (0.22 ±0.15), NODE_L (0.24 ±0.14). Finally, GWGH was generally correlated with YIELD (0.27 ±0.18), with a stronger correlation in Kenin-Keni 2012 (0.46 ±0.01). A correlation analysis also helped us to identify the five ECs that were the most correlated with the phenotype and the time window where the association was the strongest (Figures S5 and Tables S4).

**Figure 4.**
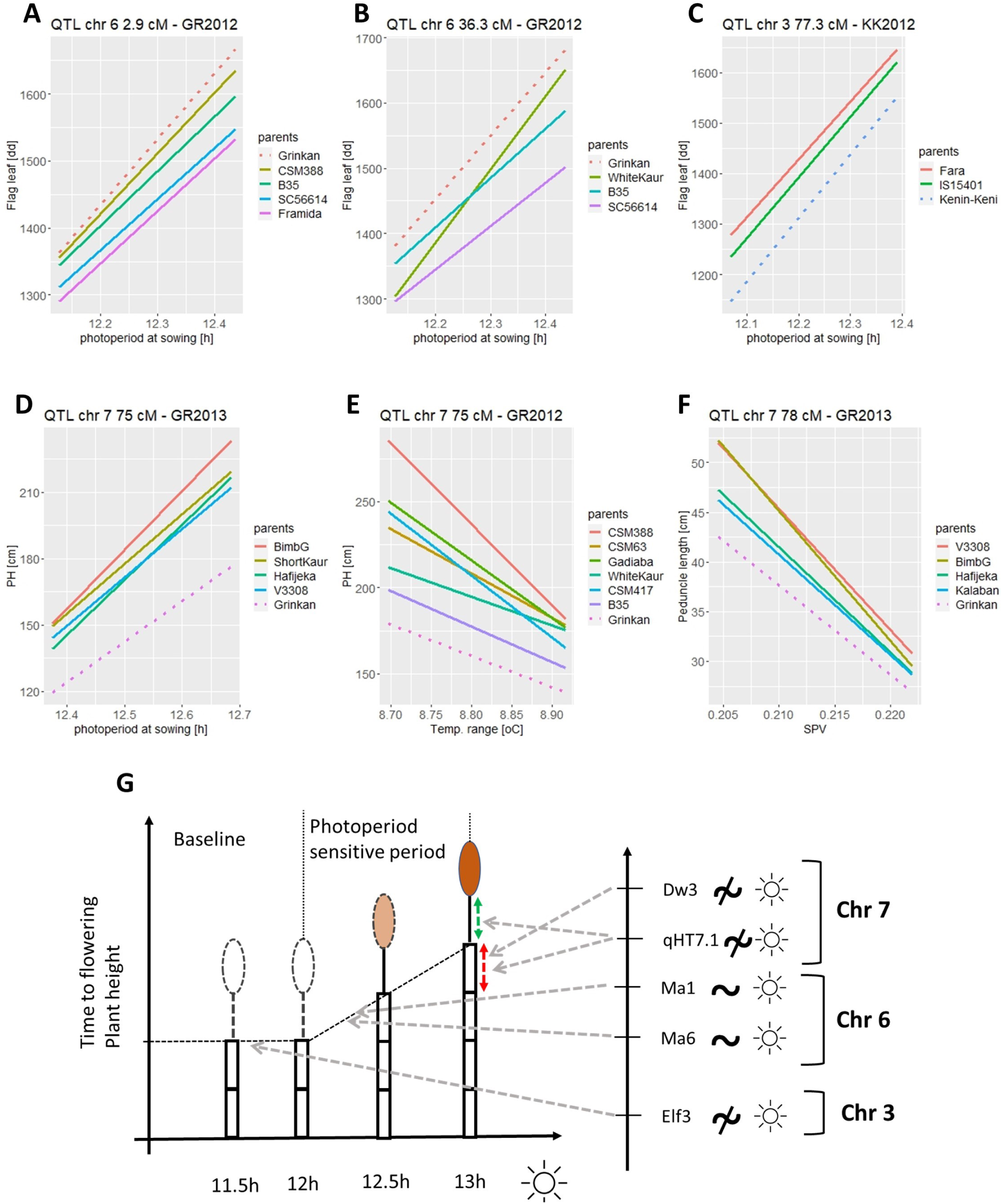
Sensitivity of QTL parental alleles effects to environmental covariables. The plain lines represent the regression between the value of parental allele effect in 4 environments and the value of an environment covariate in the same environments. A) QTL chr 6 (3 cM) in Grinkan 2012; B) QTL chr 6 (36.3 cM) in Grinkan 2012; and C) QTL chr 3 (77.3 cM) in Kenin-Keni 2012. D) Significant parental allelic effects on plant height given photoperiod at QTL chr 7 (76 cM) in Grinkan 2013. E) Significant parental allelic effects on plant height given VPD at QTL chr 7 (76 cM) in Grinkan 2013. F) Significant parental allelic effect on peduncle length given SVP at QTL chr 7 (78 cM) in Grinkan 2013. G) Photoperiodism illustration and summary of the candidate gene effects on plant cycle (time to flowering) and plant height represented as a function of photoperiod (day length [h]). Flowering time is a broken linear function with a constant baseline period and a slope describing the sensitivity to photoperiod. PH is decomposed into number of internodes, internode length, peduncle and panicle lengths.

**Figure 5.**
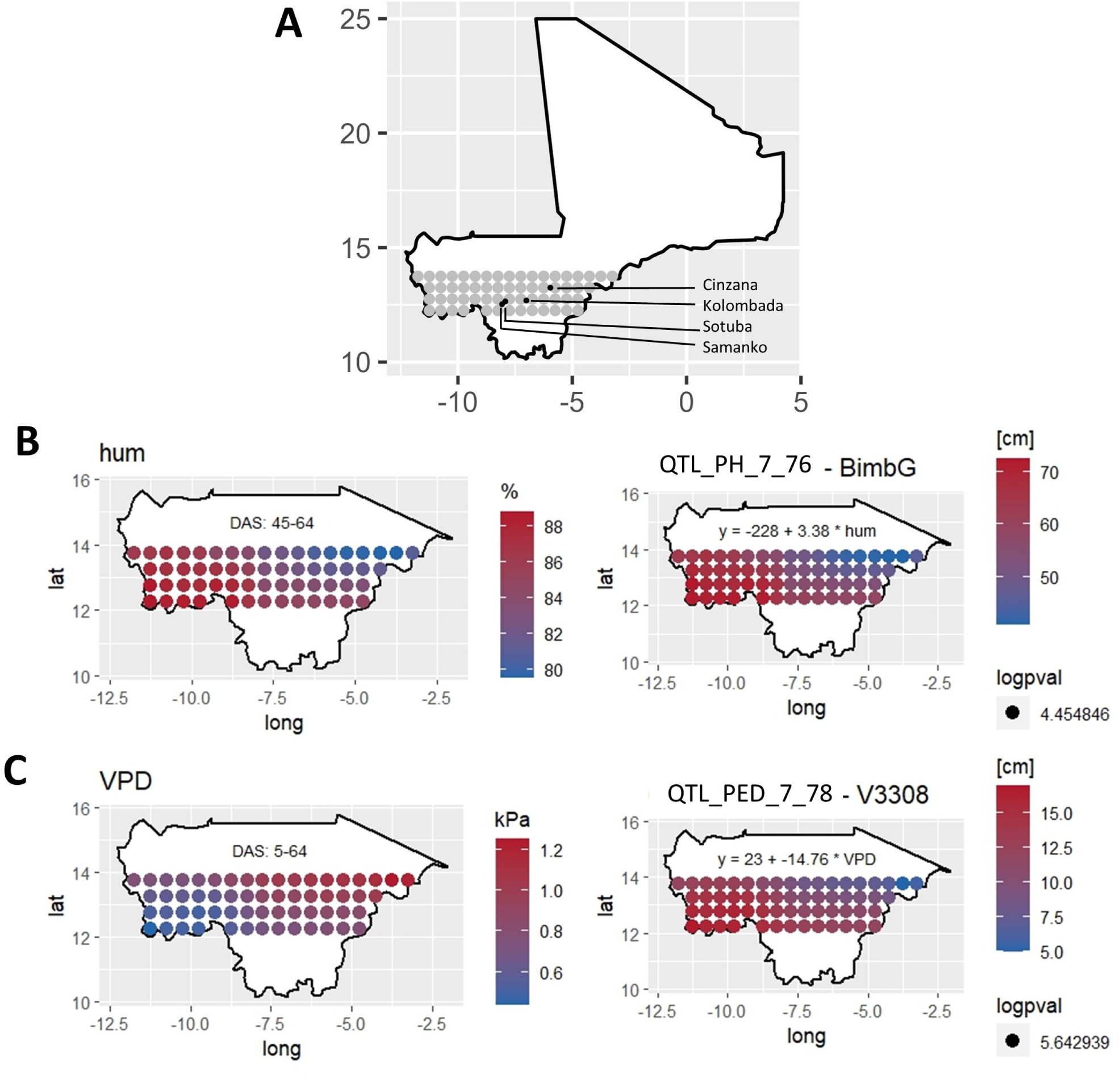
A) Map of Mali with testing locations and neighbouring areas of projection. B-C) Projections of QTL parental allelic effects for plant height and peduncle length in the environment of interest given sensitivity to humidity and VPD, respectively. The two QTL positions and parental allele were B) QTL_PH_7_76 BimbG x humidity; and C) QTL_PED_7_78 V33/08 x VPD. The projections were obtained by substituting observed environmental covariable in a grid of 60 points of the Malian environment in the QTL allele by environmental covariable sensitivity equation.

### F. QTL detection - general results

The total length of the consensus genetic map was 1412 cM with a number of cross-over equal to 47’669, 20’343, and 20’120 in the Grinkan, Kenin-Keni, and Lata3 populations, respectively (Table S1). Overall, we detected 100 significant QTL over the five populations for eight traits, which represented 64 unique QTL (Table S2, Figures S6). Consistently with the heritability estimates, the total variance explained by the QTL effects was rather large for FLAG (32-53), PH (10-48), and NODE_L (11-47), moderate for PED (10-32), NODE_N (10-22), and GWGH (8-30), and low for YIELD (4-14) and PAN (5-9).

**Table 1.** List of large and medium effect QTL with trait, chromosome, position, average *R*_2_, QTLxE effect range, number of parental alleles with significant effects, and candidate genes

| QTL ID | trait | chr | range [cM] | range [Mbp] | $R^2$ | QxE range | Npar | Candidate genes |
| --- | --- | --- | --- | --- | --- | --- | --- | --- |
| Q_FL_3_78 | FLAG | 3 | 77.34-78.36 | 5.11-5.15 | 17.1 | [-123; 144] [dd] | 24 | Elf3, SbCN12 |
| Q_NN_3_78 | NODE_N | 3 | 78.13-78.75 | 5.14-5.17 | 9.3 | [-1.8;2.1] [n] | 16 | Elf3, SbCN12 |
| Q_FL_6_3 | FLAG | 6 | 1.49-2.94 | 0.04-0.08 | 19.4 | [-178; 130] [dd] | 24 | Ma6 |
| Q_NN_6_2 | NODE_N | 6 | 1.49-2.73 | 0.04-0.08 | 7.9 | [-2.6;1.8] [n] | 19 | Ma6 |
| Q_FL_6_38 | FLAG | 6 | 36.32-39.4 | 4.04-4.12 | 6.3 | [-192; -27] [dd] | 19 | Ma1 |
| Q_FL_9_105 | FLAG | 9 | 103.7-106.7 | 5.46-5.54 | 2.7 | [-40;91] [dd] | 23 | SbFL9.1 |
| Q_PH_7_76 | PH | 7 | 74.28-76.69 | 5.47-5.52 | 21.9 | [-37; 69] [cm] | 20 | qHT7.1, (Dw3) |
| Q_NL_7_78 | NODE_L | 7 | 76.29-79.59 | 5.51-5.58 | 29.9 | [-0.1;4.2] [cm] | 16 | qHT7.1, (Dw3) |
| Q_PED_7_78 | PED | 7 | 74.8-82.1 | 54.84-56.26 | 12.6 | [-8; 10] [cm] | 24 | qHT7.1, (Dw3) |
| Q_NL_7_98 | NODE_L | 7 | 96.1-100.7 | 5.83-5.91 | 11.8 | [-5.8-4.2] [cm] | 8 | Dw3 |
| Q_PH_7_106 | PH | 7 | 102-108.3 | 5.94-6.07 | 7.6 | [-14;58] [cm] | 15 | Dw3 |

### G. QTLxEC extend

The 100 significant QTL covered 1056 parental alleles for which we could estimate the significance of the main and GxE additive effects (Tables S5). Around 60% of the parental alleles were significant and around 25% interacted with the environment. Overall, around 15% of the parent alleles interacted with at least one EC. The FLAG, PH and PED QTL were more significantly affected by the EC than the one for PAN and YIELD. For example, photoperiod strongly influenced FLAG, NODE_N and PH QTL. Atmospheric EC like VPD influenced PED and NODE_L QTL sensitivity while YIELD QTL were sensitive to humidity. PAN and NODE_L QTL were sensitive to minimum temperature.

### H. QTL with large effects and candidate genes

Eleven QTL showed medium to large effect with strong significance and consistency over several populations and environments (Table 1 and Figures S6). Their parental effect could go up to 300 dd for the FLAG QTL, or up to 1.07 m for PH. On chromosome three, we detected a strong QTL for FLAG (QTL_FL_3_78) significant in all populations and environments. Almost at the same position, we also detected a large effect QTL for NODE_N (QTL_NN_3_78). QTL_FL_3_78 and QTL_NN_3_78 are probably linked to the early flowering (Elf3) candidate gene (Guitton et al., 2018) or SbCN12 (Yang et al., 2014). Another FLAG QTL (QTL_FL_6_3) with a consistent effect in all populations and environments was detected at the beginning of chromosome six. It colocalized with a QTL for NODE_N (QTL_NN_6_2). Those QTL could be related to the Ma6 gene (Rooney and Aydin 1999; Murphy et al., 2014). We also detected a QTL with medium effects on FLAG on chromosome six around 36 cM (QTL_FL_6_38) falling in the region of the Ma1 gene (Murphy et al., 2011) and another on chromosome nine around 105 cM (QTL_FL_9_105) potentially close to the SbFL9.1 gene (Bouchet et al., 2017). A strong QTL for PH was detected on chromosome seven around 75 cM (QTL_PH_7_76) with significance in the Grinkan and Kenin-Keni (2013) populations. This QTL colocalized with a highly significant QTL for NODE_L (QTL_NL_7_78) and a strong and highly consistent QTL for PED (QTL_PED_7_78). Nearby this QTL, another QTL (QTL_PH_7_106) also had a large effect on PH and colocalized with a large effect QTL for NODE_L (QTL_NL_7_98). The QTL region of chromosome seven could be related to one or two genes. The main candidate gene is Dw3 (Multani et al., 2003; Brown et al., 2008). However, according to Li et al. (2015), chromosome seven could harbour two genes: qHT7.1 positioned before Dw3 would influence both the stem and the peduncle length while Dw3 would only influence the stem length.

### I. Complex QTL effect pattern at large effect QTL

The large effect QTL showed a complex pattern with effects distributed over many parents and a wide range of effects modulated by the genetic background and the environment. Those QTL represented 40% of the QTLxEC effects. Almost all donor parents’ alleles had at least one significant EC interaction. Such a complex pattern can be illustrated for QTL_FL_3_78 (Figure 3). First, we noticed the contrasting parental effects with parents like CSM417 or CSM388 whose alleles reduced maturity while IS15401 alleles increased it. Then, we observed differences of expression due to the genetic backgrounds. For example, the allele of Fara-Fara had low effect in a Grinkan background while it strongly increased maturity in the Lata3 and Kenin-Keni backgrounds. Finally, we could also observe environmental differences like the stronger cycle reduction of CSM388 allele in 2012 compared to 2013.

### J. QTL effect on photoperiodism

The plots of Figures 4 (A-C) represent QTL alleles effect given photoperiod compared to the recurrent (reference) parent score. QTL_FL_6_3 (Ma6) was the most photoperiod sensitive QTL (Figure 4A). At that position compared to Grinkan, the alleles of CSM388 and B35 reduced photoperiodism. For the second sowing date characterized by shorter photoperiod ( 12.1 h), the difference with Grinkan was small but it increased with longer photoperiod ( 12.5 h, first sowing). The slope of CSM388 or B35 is therefore less steep than the one of Grinkan. This photoperiod sensitivity reduction was observed in all genetic backgrounds.

QTL_FL_6_38 (Ma1) was also sensitive to photoperiod with five parental alleles interacting significantly with the photoperiod over the different genetic backgrounds. For example, in Grinkan population (2012), the allele of White Kaura increased the photoperiod sensitivity compared to the recurrent parent (Figure 4B). At QTL_FL_3_78 (Elf3), the parental alleles were mostly insensitive to photoperiod. We only detected a reduced photoperiod sensitivity for the Fara-Fara and IS15401 alleles compared to Kenin-Keni (Figure 4C). The alleles of QTL_FL_9_105 (SbFL9.1) were also insensitive to the photoperiod with only two significant interactions out of 15 possible.

### K. Dissecting plant height genetic determinism

The phenotypic data for PH and its components (NODE_N, NODE_L, PED, PAN) allowed us to dissect PH genetic architecture. PH can be expressed as PH = (NODE_N * NODE_L) + PED + PAN. Since the phenotypic values of NODE_N are strongly correlated with FLAG, it was not surprising to find overlapping QTL for the two traits on chromosomes three and six (QTL_NN_3_78, QTL_NN_6_2). In terms of photoperiod sensitivity, the QTL for NODE_N followed a similar pattern than the ones from FLAG. QTL_NN_3_78 (Elf3) was rather insensitive to photoperiod with only two parental alleles having a significant interaction while QTL_NN_6_2 (Ma6) was more sensitive with five significant alleles (e.g. Malisor).

Concerning PH, we also observed a strong agreement between the QTL positions detected for PH and NODE_L on chromosome seven. QTL_PH_7_76 and QTL_NL_7_78 (qHT7.1) colocalized while QTL_PH_7_106 and QTL_NL_7_98 (Dw3) were separated by less than 10 cM. The QTL influencing NODE_L were not sensitive to photoperiod, but other ECs like VPD or potential evapotranspiration modulated the parental allelic effects at those positions. For example, at QTL_PH_7_76, the effects of parents like Hafijeka or Short Kaura were reduced when VPD increased (Figure 4E). Surprisingly, the corresponding QTL (QTL_PH_7_76) detected for PH showed significant interaction with the photoperiod (Figure 4D) in the Grinkan populations. We consider that this apparent effect of photoperiod on QTL_PH_7_76, is due to the fact that PH is proportional to the interaction NODE_N * NODE_L. Therefore, at chromosome seven, the signal is due to the interaction between a photoperiod sensitive component (NODE_N) and a photoperiod insensitive part (NODE_L). The separate analyses of NODE_N and NODE_L helped us to clarify the GxE effects influencing PH.

In terms of PED, QTL_PED_7_78 (qHT7.1) was one of the most environmentally sensitive QTL. This QTL was not photoperiod sensitive but covariables like SVP had a negative effect on the propensity to increase PED compared to the reference parent. This effect was consistent in the Grinkan (2013) population with four parents (BimbG, Hafijeka, Kalaban, V33/08) reducing their propensity to increase PED when SVP increased (Figure 4F). It is interesting to emphasize that drought related ECs (VPD, SVP) influenced both QTL_PH_7_76 and QTL_PED_7_78 with effects going in the same direction.

### L. QTL effects on yield

Few QTL were detected for YIELD. However, some of the YIELD QTL colocalized with large or medium effect QTL for FLAG, which gave us the possibility to analyse the influence of FLAG on YIELD. A first example of collocating FLAG and YIELD QTLs was positioned on chromosome six at 38 cM (QTL_FL_6_38) and 39 cM (QTL_YLD_6_39). Here, we observed that, compared to Grinkan, the allele of parents B35 and SC566-14 reduced the photoperiod sensitivity. Thus, at the first sowing date characterized by a longer photoperiod the plants carrying B35 or SC566-14 alleles had a reduced cycle. Such a reduction could prevent those plants from accumulating biomass that will be reallocated to the grain, which ultimately reduces the yield. Indeed, at QTL_YLD_6_39, we could observe a negative effect on yield for B35 and SC566-14 alleles. Such an indirect QTL effect on yield via the duration of the plant cycle was confirmed by an analysis of the YIELD values residual after regression on FLAG (Table S7) for the B35 allele. For the allele of SC566-14 however we would rather make the hypothesis of an independent effect on both traits due to a unique or two closely located QTL.

In Kenin-Keni 2012, the region of QTL_FL_3_78 contains QTL for FLAG (78 cM) and YIELD (74 cM) with a strong allelic effect of IS15401. Here, the allele of IS15401 increased the cycle length and decreased yield. The analysis of YIELD conditional on FLAG (Table S6) supports the hypothesis of an indirect QTL effect on YIELD through FLAG modulation. At that position, the IS15401 allele increased the cycle length and decreased the yield. The extended maturity given by IS15401 could make the plant falling outside the time when optimal conditions for the Malian agroecology happen. Such a negative effect on yield for varieties flowering outside the optimal time was already observed by Curtis (1968).

### M. Expected QTL effect beyond the tested environments

Between 12 and 13.75 degrees of latitude, the Malian environment is characterized by a Southwest to Northeast gradient (Figure S7). The Southwest is cooler with lower temperature range, higher precipitation, and humidity while the Northeast is drier with higher temperature ranges and lower precipitations. A final extension of our results is the projection of QTL allelic effects having a significant interaction with one of the ECs in the Malian environment. For that we substituted the observed EC values from a grid of 60 points in the estimated allele sensitivity equation. This visualisation allowed us to map the expected effect of an allele given environmental conditions.

On Figures 3, we represented the expected behaviour of the BimbG and V33/08 alleles at QTL_PH_7_76 and QTL_PED_7_78, respectively. The BimbG allele was positively influenced by humidity which increases its effect on PH in the more humid southwest part and reduces it in the drier northeast regions (Figure 5B). Figure 5C illustrates the effect of VPD on the effect of V33/08 allele at QTL_PED_7_78 on PED extension. PED extension was reduced in the Northeast drier regions, while it was increased in the more humid southwest part of Mali. We can emphasize that those two alleles react similarly to the environmental gradient by increasing more the plant height in the southwest part.

## Discussion

### N. Multi-reference BCNAM design properties

A major contribution of this work is the development of a new sorghum genetic resource taking the form of a multi-reference BCNAM design. Its usefulness can be evaluated given criteria like genetic diversity, mapping power and resolution, and potential for genetic gain, which will have different relative importance given usage of the population (research or breeding, Gage et al., 2020, Scott et al., 2020). In terms of breeding, the use of three recurrent parents instead of one like in almost all the (BC)NAM populations (e.g. Jordan et al., 2011, Bouchet et al., 2017, Mace et al., 2021) substantially increases the exploitable genetic diversity generated.

In terms of QTL detection, the usefulness of a design can be evaluated in terms of detection power, capacity to estimate and trace the QTL effect, and resolution. The power gain offered by NAM design compared to bi-parental populations (Li et al., 2011) or association panels (Bouchet et al., 2017) was already demonstrated. This should also be valid for our population. The most relevant advantage of a multi-reference BCNAM design is the possibility to test allelic contribution in several genetic backgrounds. Similarly to Christopher et al. (2021), we could show that parent allelic effect can be strongly modulated by the genetic background (Figure 3). More generally, in terms of QTL effect characterisation, MPPs like the (BC)NAM designs increase the allele frequencies and allow the user to trace back the allelic effect to a specific parent (Myles et al., 2009), which is fundamental to characterize the genetic effects and, if relevant, to mobilize it for breeding purposes. A second important contribution of this work is the capacity to link source of adaptation to specific parental allele. The main disadvantage of (BC)NAM design is the low resolution compared to designs involving further intercrossing like the multiparent advanced generation intercross (MAGIC, Klasen et al., 2012; Garin et al., 2021). Even if (BC)NAM design involves more recombination than bi-parental populations, recombination is still restricted within the cross which considerably extends the linkage disequilibrium decay. Even if the combination of our design to strategies like RapMap (Zhang et al. 2021) could improve the mapping resolution, populations like association panels could already be helpful to better distinguish between one or two QTL scenarios for PH at chromosome seven, for example.

In terms of genetic gain, Bernardo (2021) showed that populations like MAGIC do not have a significant advantage compared to multiple cross populations like the BCNAM. Moreover, populations involving further intercrossing like MAGIC request more resources. The (BC)NAM populations whose design is closer to the standard breeding crossing practice can also have an interest for genomic selection because of the genetic similarity it contains (Scott et al., 2020).

An important question specific to the multi-reference BCNAM design is the need to cross all donor parents to the recurrent parents (full factorial) or only a subset. Such a problem represents a trade-off between a) the advantage of estimating the QTL allelic effect in multiple genetic backgrounds, and b) an increase of the covered genetic diversity (more donor parents) or of the cross size by reducing their number, which increases the QTL detection power (Garin et al., 2021). Even if the quality of the QTL effect estimation is important we consider that it is conditioned on the covered genetic diversity and the QTL detection power. Therefore, performing only a selected number of crosses in a multi-reference (BC)NAM design could be an interesting strategy. More definitive answers to this question could be obtained by simulations.

### O. Statistical methodology properties

At the moment of writing this article, no statistical package for MPP-ME QTL detection was publicly available, which made a direct comparison impossible. Nevertheless, we can compare our methodology to general properties from similar methods. The most important criterion is the precision of the QTL effect estimation, which influences the detection power and the interpretation of the effect. This criterion is often balanced with computational power requirements.

We used a mixed model with unstructured variance covariance (VCOV) structure to control for the genetic (co)variance. We could reduce the scanning time using approximation, but we estimated the final QTL effect with a full model. Considering QTL as fixed allowed us to use well established mixed model estimates and Wald test for significance. A small disadvantage of fixed QTL effect is the need to interpret the effect with respect to a reference. The use of a random QTL effect like in Li et al. (2022) with parental effect expressed as deviation from zero could facilitate the QTL effect comparison but it creates other challenges for significance estimation and computational speed. The procedure from De Walsche et al. (2022) proposes to perform MPP-ME QTL detection as a meta-analysis of single environment analysis. Such an approach is very fast, but we expect the joint multi-environment analysis to be more precise concerning the QTL effect estimation because it jointly accounts for multiple sources of variations (Malosetti et al. 2013).

The main innovation of our methodology was the extension with envirotyping data like the ones generated by Nasapower. The benefit of extending the genetic models with environmental covariates to increase their prediction ability was already demonstrated (e.g. Westhues et al. 2022). This opens the door to dynamic predictions beyond the tested environments that should be used with caution because the estimated sensitivity equations are potentially strongly influenced by the tested environments. The possibility to increase the number and the representativity of the testing sites should improve the model prediction ability.

### P. Developing climate-smart varieties

An important contribution of this study is to demonstrate the connection between maturity and plant height at the molecular level via large effects QTLs and underlying genes showing various degrees of photoperiodism (Figure 4G). This understanding could support the development of climate-smart varieties in terms of maturity and biomass accumulation.

The expected rise in the variability and intensity of the African climate in terms of temperature, drought and flood events should affect flowering time (WMO 2022). To develop varieties with a wide geographical adaptation, sorghum breeding programs have mainly selected early flowering and photoperiod insensitive genotypes. This approach failed to produce efficient varieties because, in the sub-Saharan African context, photoperiod sensitivity is the main adaptation trait to climate variability (Sultan et al., 2013). Photoperiod sensitivity ensures the synchronization of flowering with the probable end of the rainy season independently of the sowing date (Kouressy et al., 2008). As we could see, at least two QTL regions influencing the cycle length might have an indirect effect on yield by synchronizing or desynchronizing the plant with the optimal flowering time.

Contrary to Mace et al. (2013) who strongly constrained the maturity of their population genotypes, we could identify QTL with large effects on flag leaf appearance. Those QTL could be used in breeding programs. The QTLxEC analysis revealed that the parental alleles of QTL_3_FL_78 (Elf3) and QTL_FL_9_105 (SbFL9.1) were mostly insensitive to photoperiod. Those QTL mostly affect the flowering baseline duration (Guitton et al., 2018) while the QTL on chromosome six (QTL_FL_6_3 and QTL_FL_6_38) linked to the Ma6 and Ma1 regions influenced both the baseline flowering duration and the photoperiod sensitivity. The photoperiod sensitive nature of the Ma6 was also identified by Takai et al. (2012). The availability of different effects allows the development of alternative breeding strategies: a) varying the duration of the cycle without affecting the photoperiod sensitivity (e.g. QTL_FL_3_78 CSM417: -80 dd); b) influencing only the photoperiod sensitivity (e.g. QTL_FL_6_3 CSM388 : -62 dd/h), or c) influencing both the baseline duration and the photoperiod sensitivity (e.g. QTL_FL_6_38 White Kaura: -49 dd and +47 dd/h).

We could also gain knowledge about important genetic regions affecting plant height. Plant height is connected to plant cycle via the Ma6, Ma1, and Elf3 genes that affect both flag leaf appearance and the number of internodes (Figure 4G). The genetic association between FLAG and NODE_N makes sense because the internode organogenesis is a function of the plant cycle (Takai et al., 2012). Given sufficient nutrients, longer maturity allows the plant to accumulate more internodes.

The other important genetic determinants of plant height were located on chromosome seven with very strong effects on the length of the internode and on the peduncle length. Our data support the results from Li et al. (2015) concerning the existence of two distinct genes (qHT7.1 and Dw3) because the QTL effects of chromosome seven were detected at different positions (around 75 cM and 100 cM) in the different populations. The phenotypic effects of those positions was also consistent with Li et al. (2015) observations because the 75 cM position influenced both the internode and the peduncle lengths while the 100 cM position only influenced internode length.

The different genes controlling plant height could also be mobilized for different breeding strategies: a) vary plant height via the internode length independently of the environment (e.g. QTL_NL_7_76 IS23540: +3.1 cm); b) vary plant height via the number of internodes conditionally on photoperiod (e.g. QTL_NN_6_2 SC566-14 -1.1 n +22.7 n/h) or unconditionally (e.g. QTL_NN_3_78 Sangatigui + 1.5 n). Those QTL could help to develop dual-purpose varieties given a parallel improvement of the fodder quality.

The environmental sensitivity of the peduncle length could also help to design climate-smart varieties with resistance to grain mold and pest attack on the panicle given local conditions. Indeed, on Figure 5C we showed that the allele of V33/08 was sensitive to VPD (-15 cm/KPa). V33/08 allele increased more the peduncle length in the more humid southwest region than in the drier northeast zones. Since peduncle length is one of the traits that can reduce grain mold and pest attack by avoiding a too close contact between the panicle and the flag leaf, we could use the climate dependent PED sensitivity of V33/08 to improve cultivar resistance. Those examples illustrate the strong application potential resulting from our design and methodology.

## Material and Methods

### Q. Plant material

The WCA-BCNAM is composed of three BCNAM populations produced after the crossing of the Grinkan, Kenin-Keni, and Lata3 recurrent parents with 24 donor parents representative of the Western African sorghum diversity with lines from Central and East Africa (Table 2, Methods S5: list of parents name synonyms). The whole population contains 3901 BC1F4 genotypes from 41 crosses (GR: 2109 genotypes, 19 crosses; KK: 896 genotypes 10 crosses; Lata: 896 genotypes, 12 crosses, Figure S1). The Lata3 population crosses involved male sterile sister lines of Lata3 to produce BC1 generation, while the BC1 generations of Grinkan and Kenin-Keni crosses were produced using manual emasculation. The recurrent parents are elite lines selected in Mali through farmer variety testing. Grinkan was developed through pedigree breeding methods. Kenin-Keni was derived from a directed recurrent selection population involving local parents of different botanical types (Leroy et al., 2014). Lata3 was selected from a random mating population of Guinea parents (Diallo et al., 2019). The recurrent parents were chosen for their productivity, their adaptation to soil and climate, as well as their resistance to major biotic and abiotic stresses. The recurrent lines also have weaknesses like poor grain quality and mold susceptibility (Grinkan), suboptimal glume opening and/or susceptibility to Striga (Lata3), and low productivity and yield stability (Kenin-Keni).

The 24 donor parents cover diverse racial (Guinea, Caudatum, Durra) and geographical origins (Table 2). They are characterized by key adaptive traits like height, maturity, or photoperiod sensitivity (Kp3). Those parents were also selected for presenting more positive specific traits like tolerance to Striga, to soil phosphorus deficiency and/or to drought, good grain quality, or panicle desirability that could increase farmer acceptance. Several donor parents were tested in multiple genetic backgrounds. Fara-Fara, IS15401, and IS23540 were crossed with the three recurrent parents. Ten donor parents were tested in two genetic backgrounds. During the development of the populations, a moderate selection pressure was applied at BC1F2 generation against too early flowering and too high genotypes.

**Table 2.**
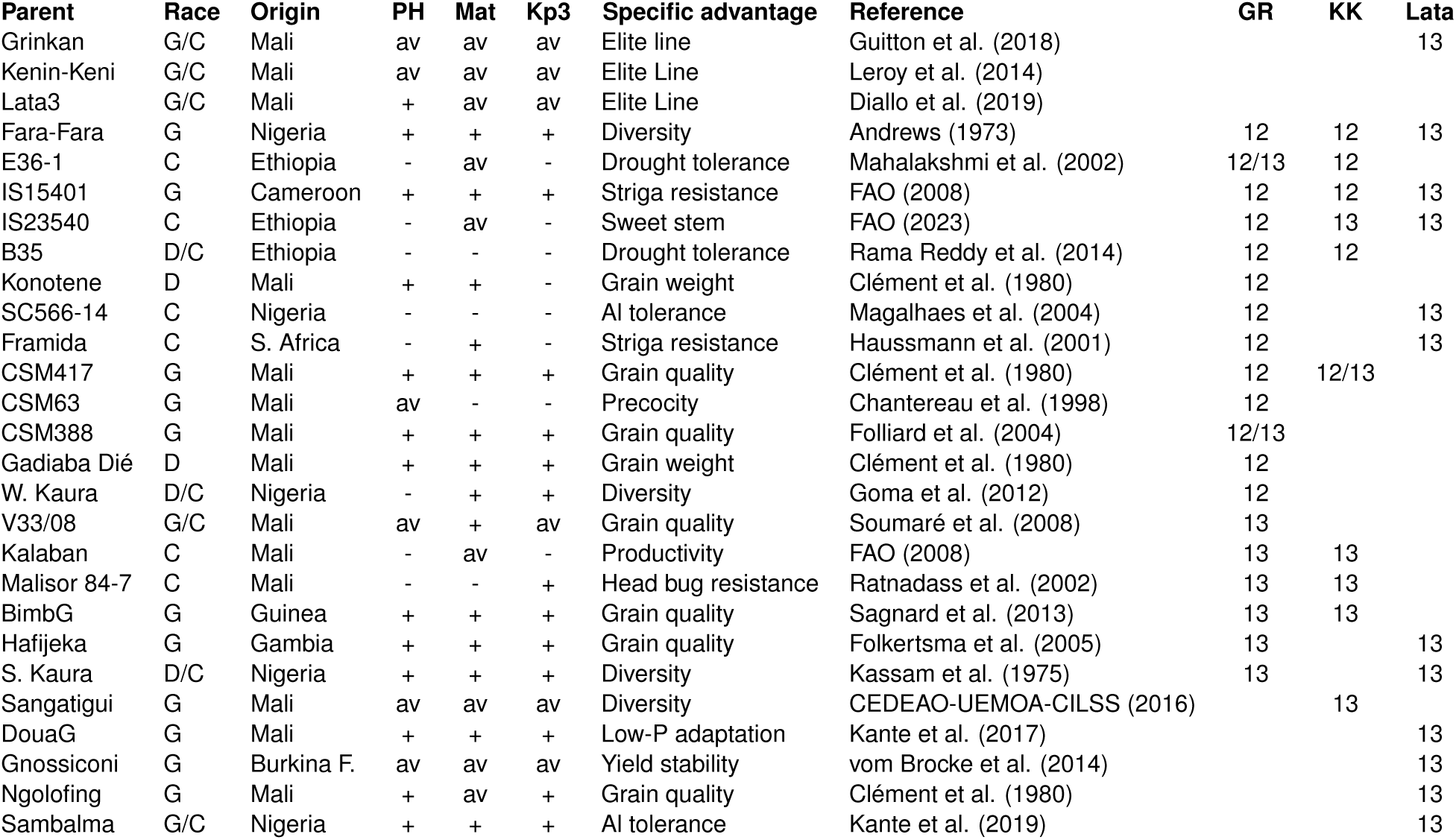
WCA-BCNAM parents with racial classification, origin, relative height (PH), relative maturity, reaction to photoperiod sensitivity (Kp3), and specific advantages. The last three columns specify the crossing scheme with the year when the cross was phenotyped (2012 and/or 2013)

#### Q.1. Genotypic data

The 3901 offspring and their parents were genotyped using genotype by sequencing (GBS, Elshire et al., 2011) with 384-plex libraries on an Illumina HiSeq 2000 sequencer. The offspring were genotyped at generation BC1F3. The sequence data were analysed running the reference genome-based TASSEL GBS pipeline (Glaubitz et al., 2014). Unique tags (3’844’911) were aligned on the sorghum reference genome v2.1 (Paterson et al., 2009). After the filtering of raw genotype data for minor allele frequencies (MAF < 0.05) and single marker missing data (<0.9), 51’545 segregating SNPs were identified between the parents with between 11’856 to 26’128 SNPs segregating in the individual crosses. Missing values in the parents were imputed using Beagle (Browning et al., 2018). Missing values in the offspring genotypes were imputed using FSFHap (Bradbury et al., 2007). We determined a unique genetic consensus map (Table S1) by projecting the 51’545 markers physical distance on a high-quality genetic consensus map (Guindo et al., 2019) using the R package ziplinR (https://github.com/jframi/ziplinR).

### R. Phenotypic data

The Grinkan and Kenin-Keni populations were partly phenotyped in 2012 and partly in 2013 (Table 2, Figure S1). We considered each population within a year as a separate population (GR2012, GR2013, KK2012, KK2013). Each population was phenotyped at a combination of two locations (Sotuba and Cinzana, Figure 5A) and two sowing dates (Sowing 1: end of June, Sowing 2: three to four weeks later, Table S8), which gave a total of four environments (SB1, SB2, CZ1, CZ2; Figure S8). The Sotuba environment is characterized by around 900 mm/year of precipitation and lower temperatures while the Cinzana environment is characterized by lower precipitation (600-700 mm/year) and warmer temperature. In both environments, the second sowing date had a lower level of precipitation and humidity and higher temperatures (Figure S7, Table S9). In each environment, the progenies of Grinkan and Kenin-Keni populations were laid out as augmented design (Kempton, 1984) with the three recurrent parents used as checks. The Lata3 population was entirely phenotyped in 2013 in three environments defined by two levels of phosphorus fertilization (Low-P and high-P) at the Samanko station and standard conditions at Kolombada station (Figure 5A). In each environment, the genotypes were laid out as an alpha-lattice design (Kempton, 1984) with two replications (Diallo et al., 2019).

We measured eight traits listed with crop ontology (CO) code. Flag leaf appearance (FLAG, CO_324:0000631) was the number of days after sowing when half of the plot had their ligulated flag leaves visible. For the QTL analysis FLAG data were converted into degree day. Plant height (PH, CO_324:0000623) was the distance in cm between the soil and the panicle top. The number of internodes (NODE_N, CO_324:0000605) was the number of nodes on the main stem minus one and the average length of the internodes (NODE_L) the main stem length divided by NODE_N. The peduncle length (PED, CO_324:0000622) was the distance in cm between the final node and the panicle bottom. The panicle length (PAN, CO_324:0000620) was the distance in cm from the end of the peduncle to the panicle top. GWGH (CO_324:0000424) was the weight in grams of 1000 grains. Finally, YIELD (CO_324:0000403) was measured in kg/ha at the plot level. All traits except FLAG were measured at harvest. Some traits like GWGH and PAN were not measured in all environments (Table S10).

### S. environmental covariables - EC

We complemented the data by 15 daily observed ECs (Table S11) to evaluate the environmental influence on plant adaptation of the Grinkan and Kenin-Keni populations. With four environments (SB1, SB2, CZ1, CZ2) we could observe environmental gradients, which was not possible for Lata3 data because the low-P and high-P trials were performed at the same time and location which reduce the number of environments to two. The ECs were divided in three categories. The atmospheric ECs contained: cumulated rain [mm], humidity [%], vapour pressure deficit (VPD - kPa), slope of saturation VP curve (SVP – kPa/d), potential evapotranspiration (ETP – mm/day), atmospheric water deficit (PETP – mm/day). The temperature ECs covered: maximum and minimum temperatures [d], temperature range [d], the effect of temperature on radiation use efficiency (FRUE; 0-1), and cumulated degree day (DD). The radiation ECs were the cumulated hours of sun (hSun), the photoperiod (day length) [h], and the solar radiation [MJ/*m*_2_/day]. We also included the photothermal time as the product between photoperiod and DD. A principal component analysis of environments based on the EC value showed a good coverage of the EC space by the four Sotuba and Cinzana environments (Figures S8).

The ECs values came from weather stations at the locations complemented by Nasapower satellite observations (Spark 2018) and transformation using the R package EnvRtype (Costa-Neto et al., 2021). We projected the genetic effects beyond the tested environments, using compiled environmental data from various sources extended with EnvRtype for a grid of 60 points between 12.25 and 13.75 degrees of latitude and -12.5 and -2 of longitude.

### Phenotypic data analysis

We estimated the genotypic variance component and broad-sense heritability (*h*_2_) using the following mixed model

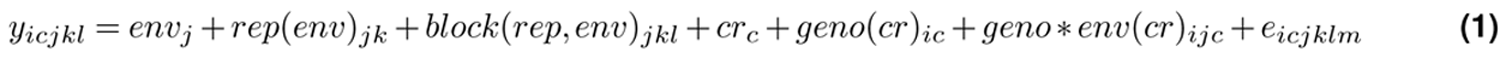

Where, *y_icjkl_* = plot phenotypic observation of the ith genotype from cross c in environment j, replication k, and block l; *env_j_* = environment effect; *rep*(*env*)*_jk_* = replication effect within-environment (only for Lata sub-population); *block*(*rep, env*)*_jkl_* = block effect within replication and environment; *cr_c_* = cross effect; *geno*(*cr*)*_ic_* = genotype effect conditional on cross; *geno env*(*cr*)*_ijc_* = GxE effect conditional on cross. The underlined terms were considered as random, the other ones as fixed.

The genotype, GxE and error terms were normally distributed with cross-specific variance 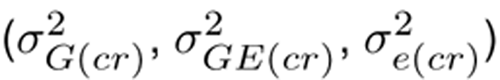. We estimated the model components using Genstat 18 (VSN International 2022). Given those, we calculated the broad sense *h*_2_ using the formula of Hung et al. (2012):

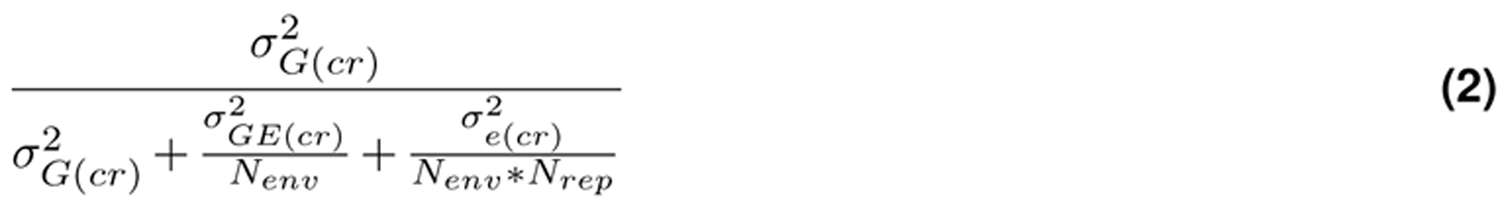

Where, *N_env_* is the number of environments and *N_rep_* the number of replications. For the multi-environment QTL analysis, we calculated within-environment best linear unbiased estimates by removing the environment and cross term from model 1 and by considering the genotype term as fixed. We used the plot field coordinates to model the spatial variation using a 2D P-spline (SpATS model, Rodriguez-Alvarez et al., 2018). For each configuration of population (GR2012, GR2013, KK2012, KK2013) by trait, we selected the five most influential ECs and time window for which the EC effect were maximal to be introduced in the QTLxEC analysis later using the method developed by Li et al. (2018) (Methods S2).

### T. MPP QTLxEC modeling

To detect QTL, we extended the linear mixed model 3 proposed by Garin et al. (2020):

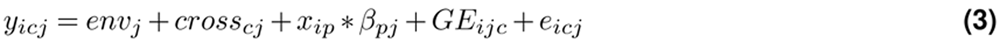

Where, *y_icj_* = BLUE of genotype i from cross c in environment j. *env_j_* = environment effect and *cross_cj_* = cross within-environment effect. *x_ip_* is the number of alleles from parent p carried by genotype i at the QTL position and *β_pj_* represents the QTL allelic effect of parent p in environment j. We assumed that each parent carried a different allele at the QTL position (Garin et al., 2017). The *GE_icj_* term is the residual genetic variation and *e_icj_* the plot error term that cannot be estimated separately due to the unreplicated nature of the BLUEs. To model the (*GE_icj_* + *e_icj_* ) term we extended the model from Garin et al. (2020) using an unstructured variance covariance structure (Boer et al., 2007). The unstructured model estimates one (co)variance (*σ*_2_ *_′_* ) for each pair of environments.

We detected QTL by performing a simple interval mapping followed by a composite interval mapping. The final list of QTL contained the positions significant at a -log10(p-val) detection threshold accounting for multiple testing (Li and Ji 2005). For further details see Methods S3. We grouped QTL detected for the same trait but in different populations into unique QTL positions if those positions were distant by less than 10 cMs. We searched for candidate genes using the sorghum QTL atlas (Mace et al., 2019).

We extended the QTL analysis by fitting a QTL by EC (QTLxEC) model for the QTL that showed a larger significance when modelled with a QTLxE term compared to a QTL main effect term. For those QTL, we extended the *x_ip_ β_pj_* term from model 2 to *x_ip_* (*β_p_* + *EC_e_ S_p_* + *l_pe_*) where *β_p_* is the main parental allelic effect across environments, *EC_e_* is the EC value (e.g. humidity) in each environment, *S_p_* is the parental allele sensitivity to *EC_e_* change and *l_pe_* the residual effect. We estimated the sensitivity to the five most influential ECs previously determined (*Q_eff_* = *β*^^^*_p_* + *EC_e_ S*^^^*_p_*). We could estimate the expected allelic effect beyond tested environments by substituting in the QTL sensitivity equation the average *EC_e_* values over the next seasons (2014-2020) for the grid of 60 points. For the QTL detection, the effect and significance were estimated using and approximate Wald test similar to the generalized least square strategy implemented in Kruijer et al. (2015; Methods S4) while for the final QTL effects model we used an exact restricted maximum likelihood solution. The methodology was added to the mppR package (Garin et al., 2023).

## U. Authors contributions

Data collection: CD, MLT, KT, MV, BG, JFR, KD, AD, MK, WL, IS, MS, NT

Statistical methodology, software, and application: VG Data analysis: VG, CD, MLT, KT, NT, JFR, MV, DP, JF

Manuscript writing, edition: VG, MLT, CD, KT, DP, MV, JK, NT, JFR

Project coordination: JFR, NT, EW, MV, FR, BN, AT, JK, VG

## V. Acknowledgements

The authors thank Madina Diancoumba and Jan Jarolímek for providing the part of the environmental data that were used to realize the projection in the Malian environment.

## Y. Data availability statement

The genotypic and phenotypic data are available here (…) [the link to the data repository will be added on acceptance of the manuscript]. The scripts to reproduce the results are available here: https://github.com/vincentgarin/sorghum_BCNAM_analysis. The results were gathered into an interactive database available here https://github.com/vincentgarin/SQE.

## Supporting figures

**Figure S1:**
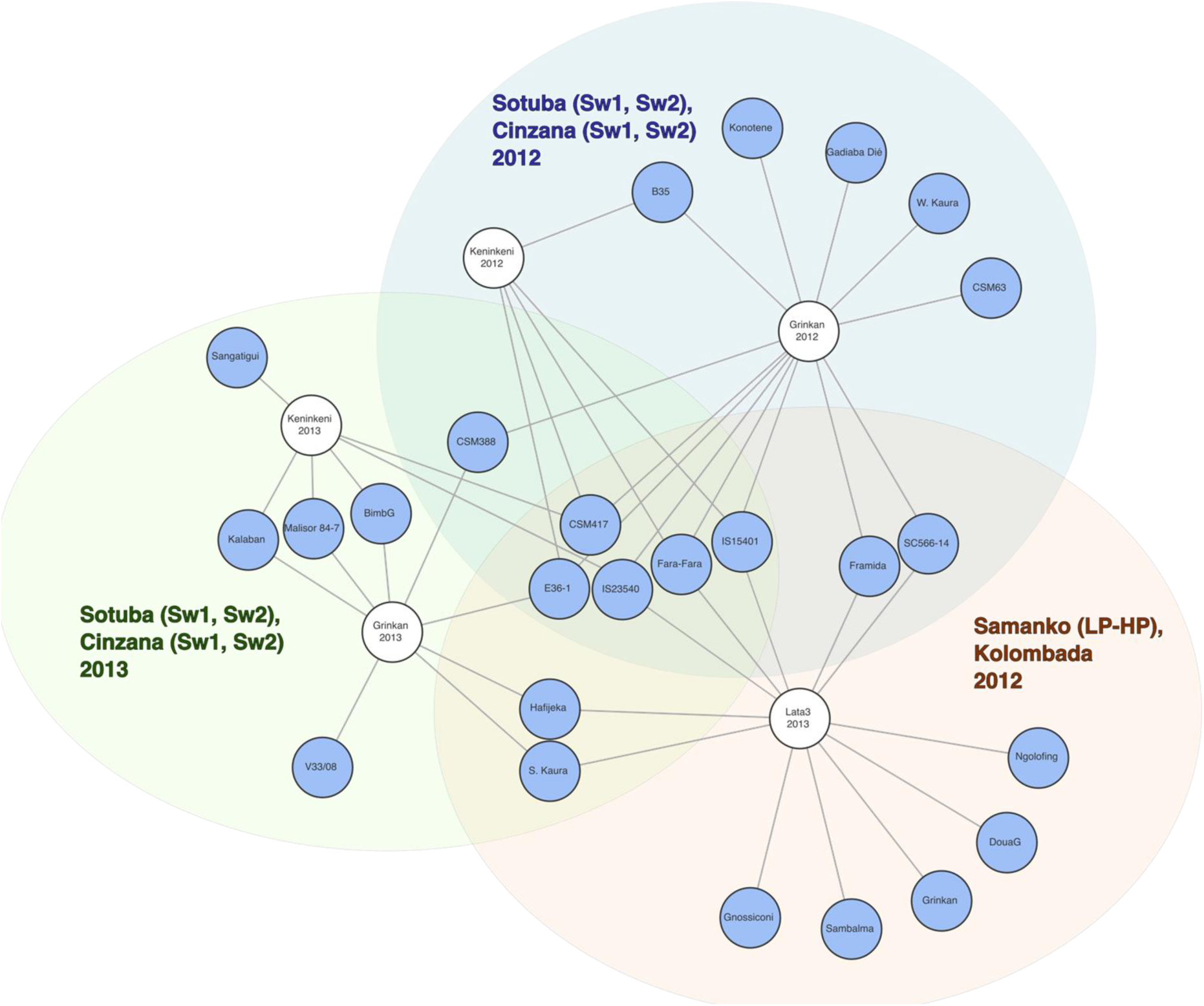
Illustration of the crossing scheme with each parent represented as a circle and cross as a line. The larger ellipses represent the phenotyping experiments

**Figures S2:**
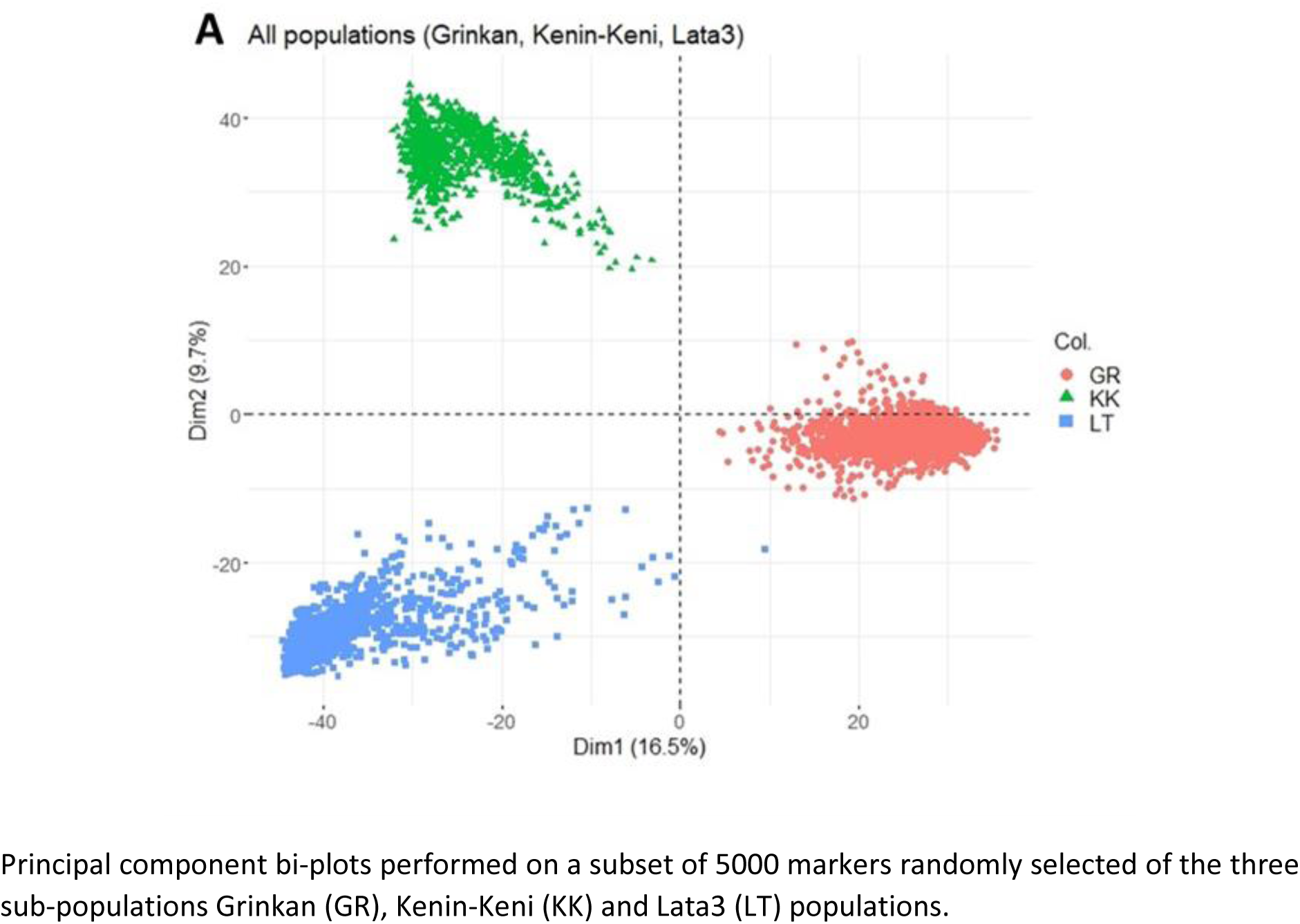
Principal component analysis of genetic information

**Figures S3:**
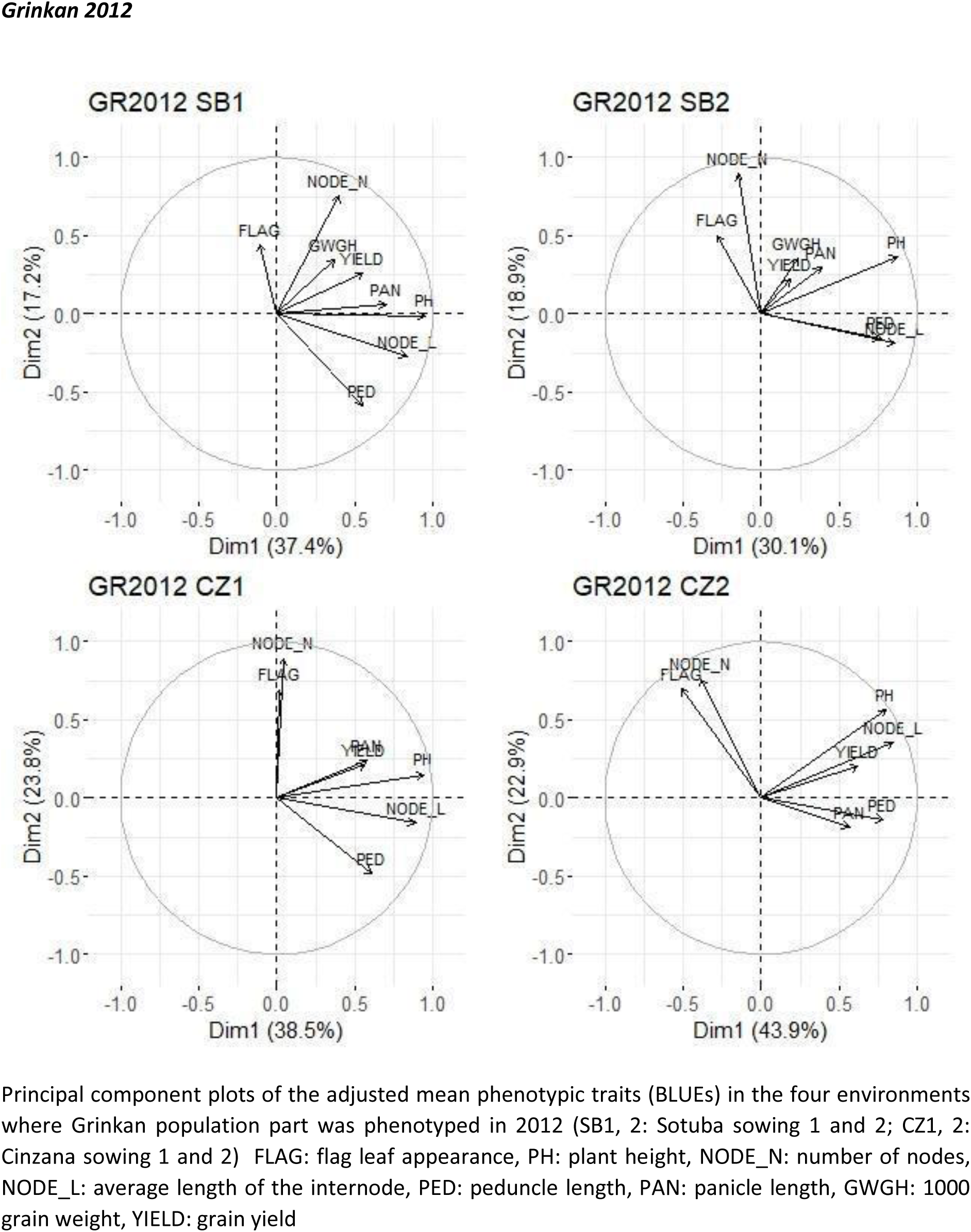
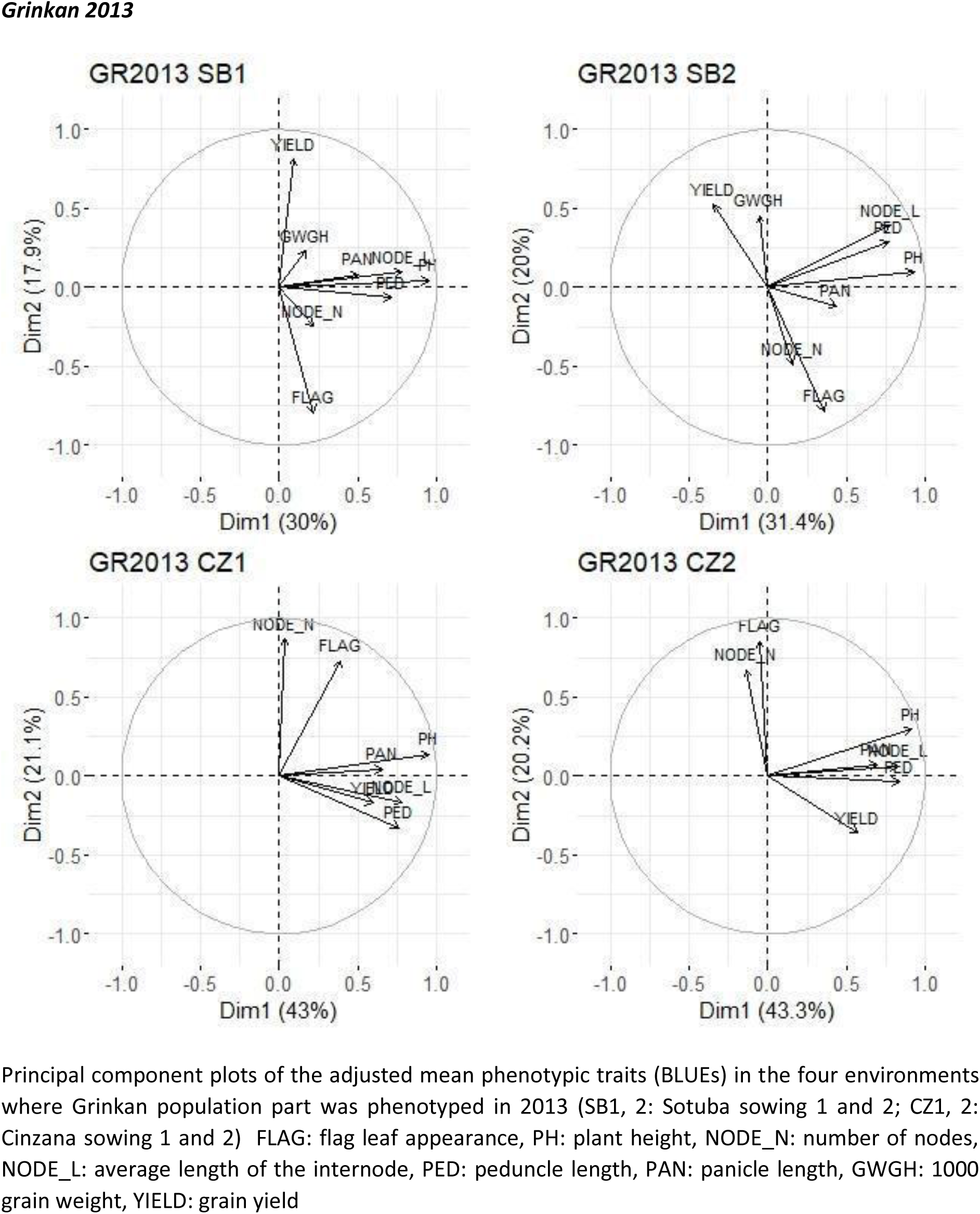
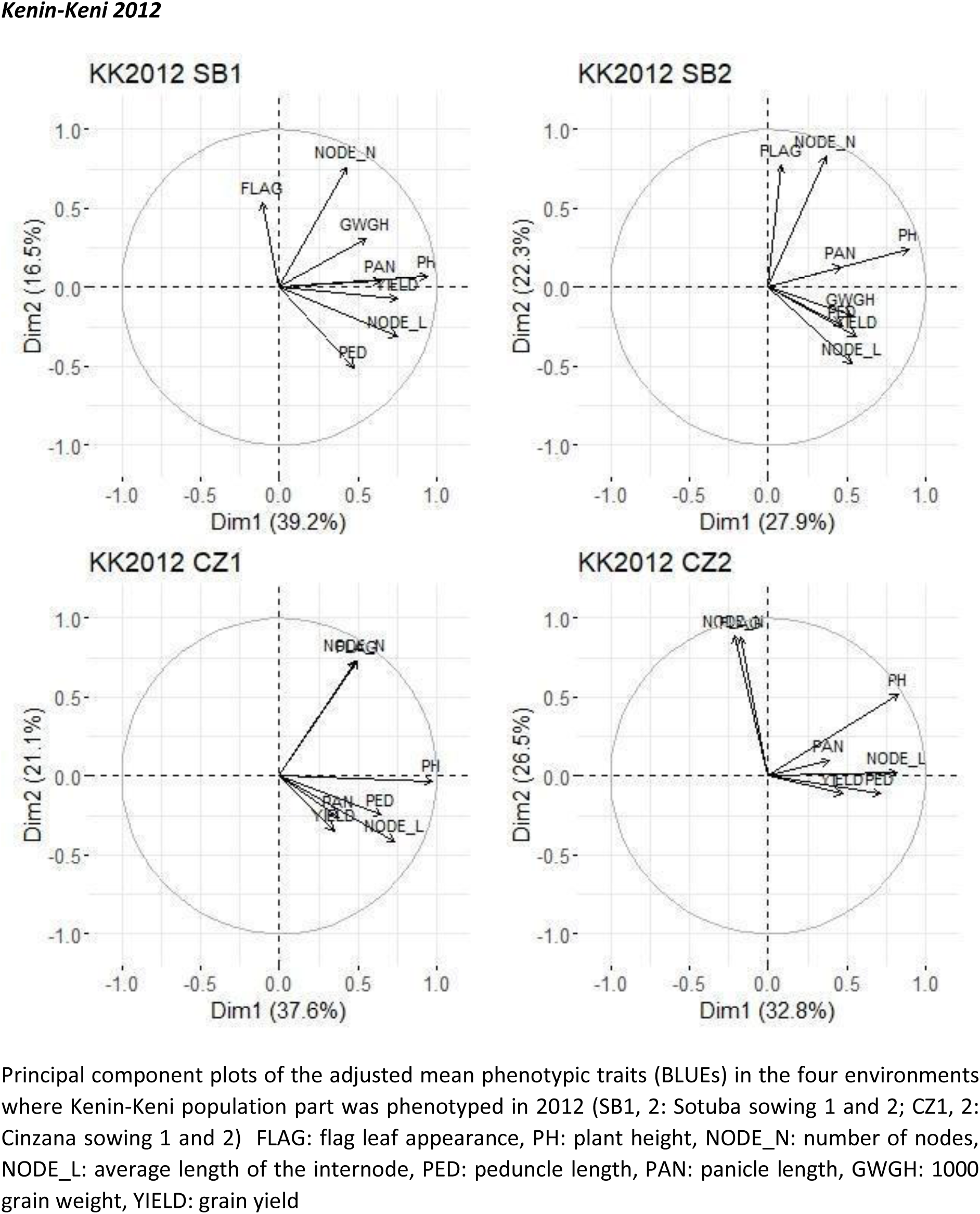
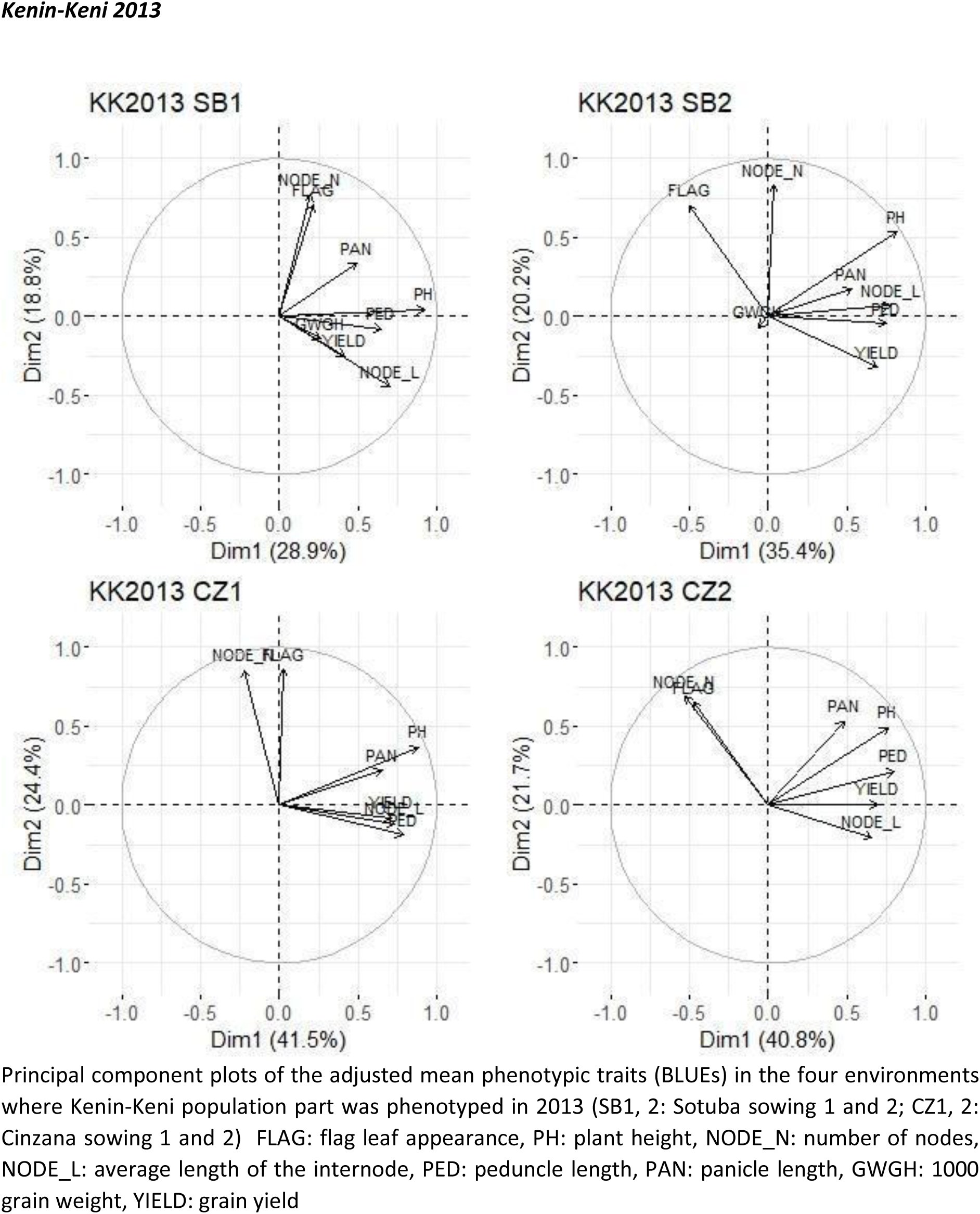
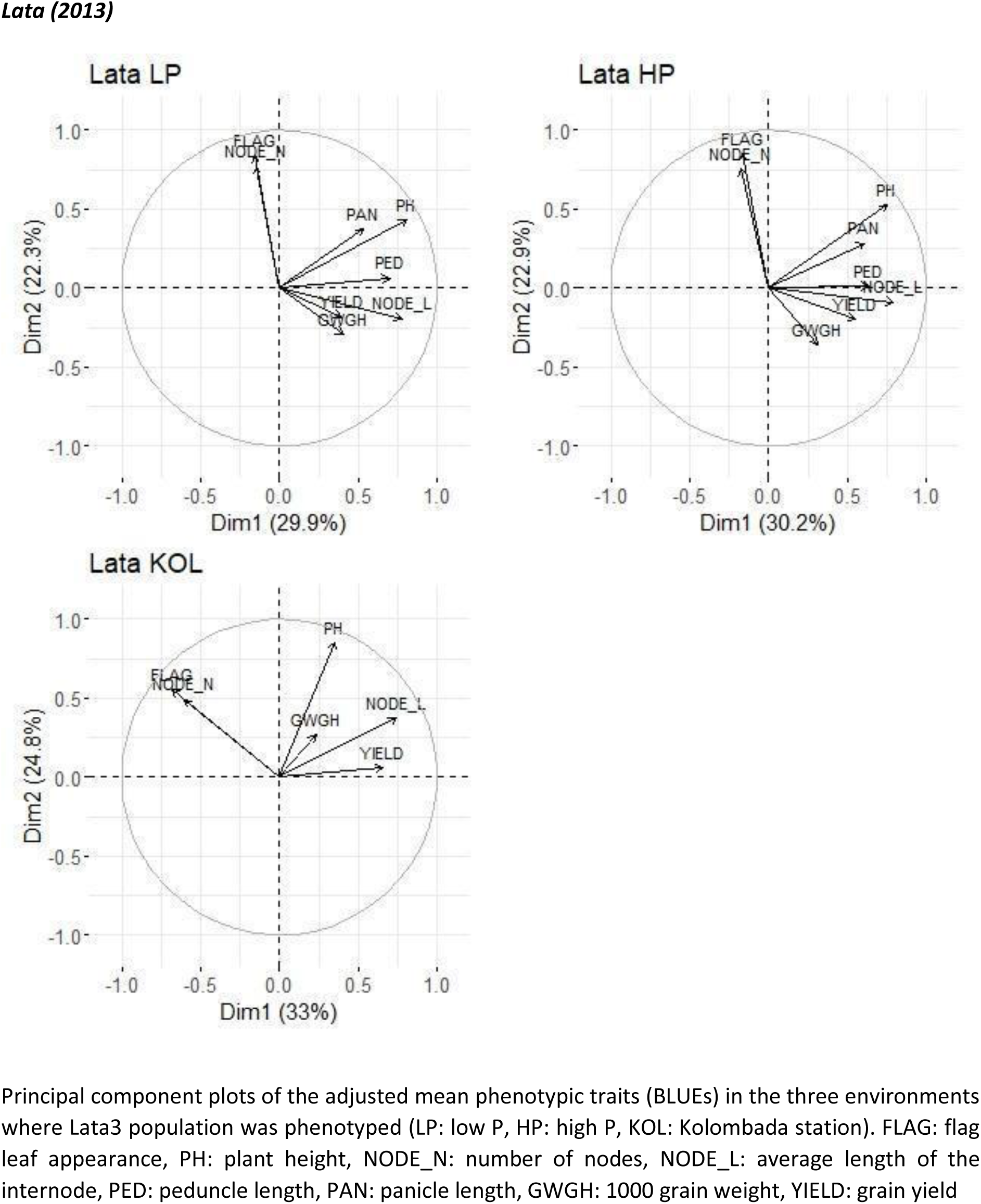
Principal component plots of the phenotypic traits for each population and year of phenotyping combination

**Figures S4:**
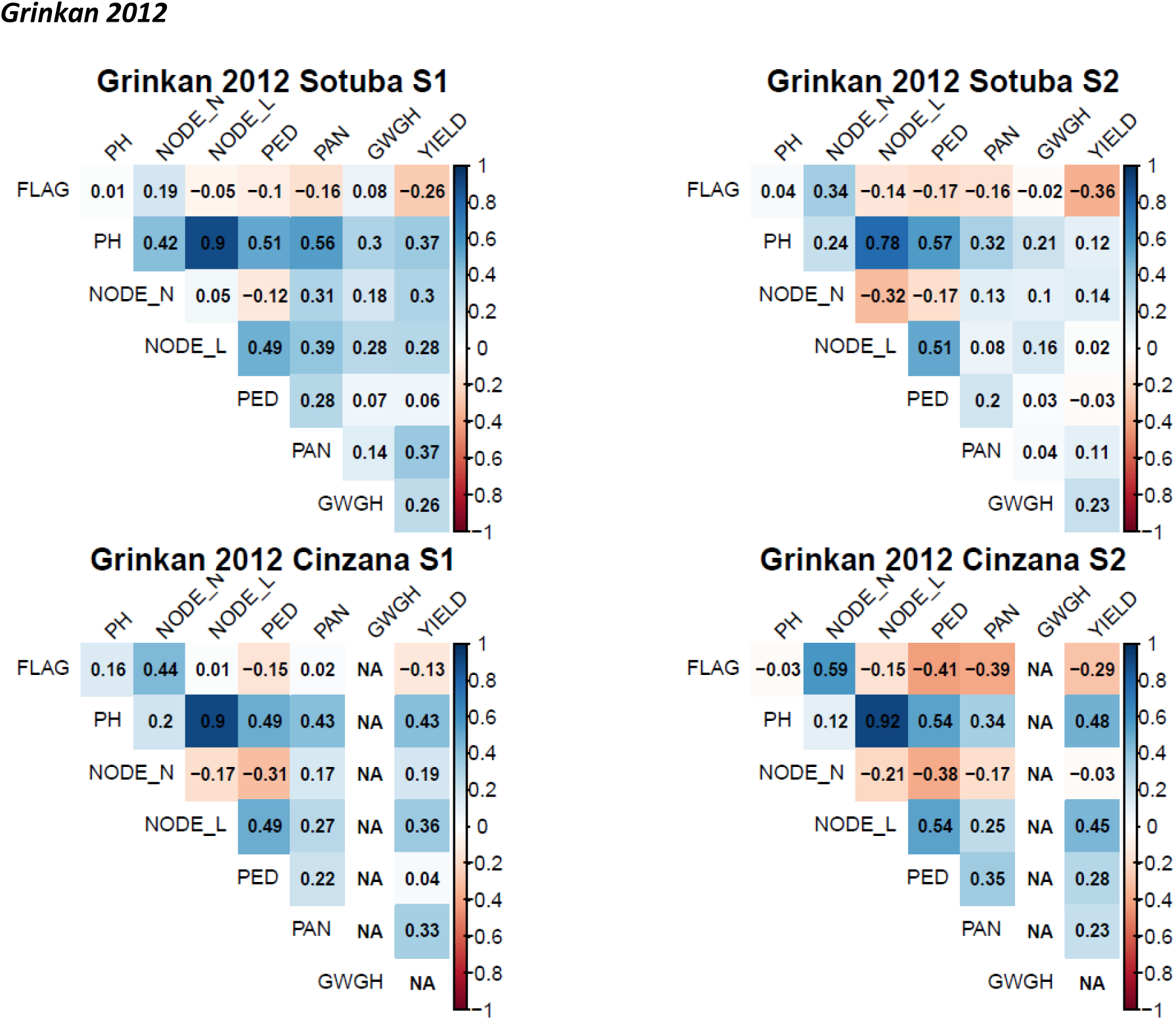
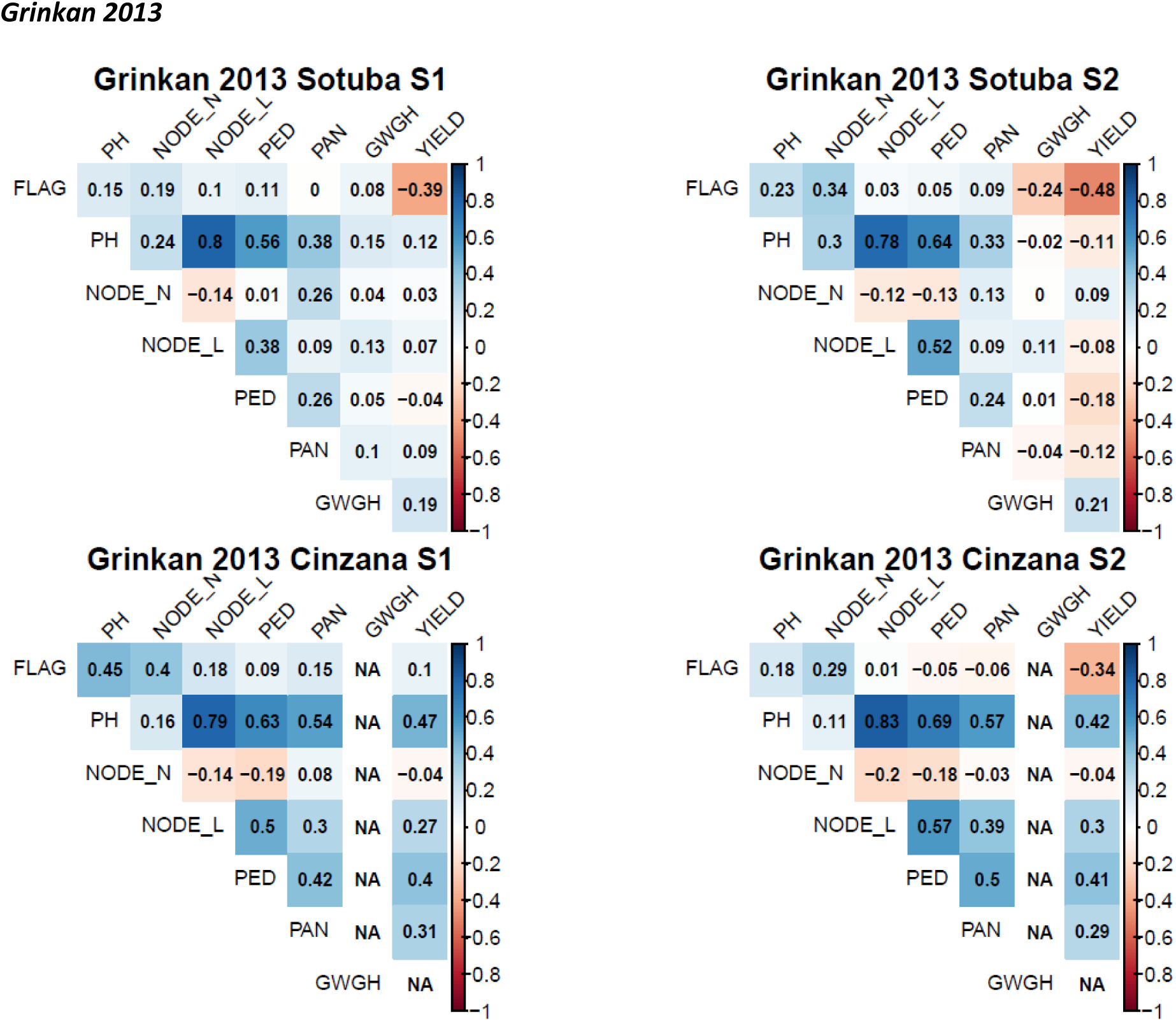
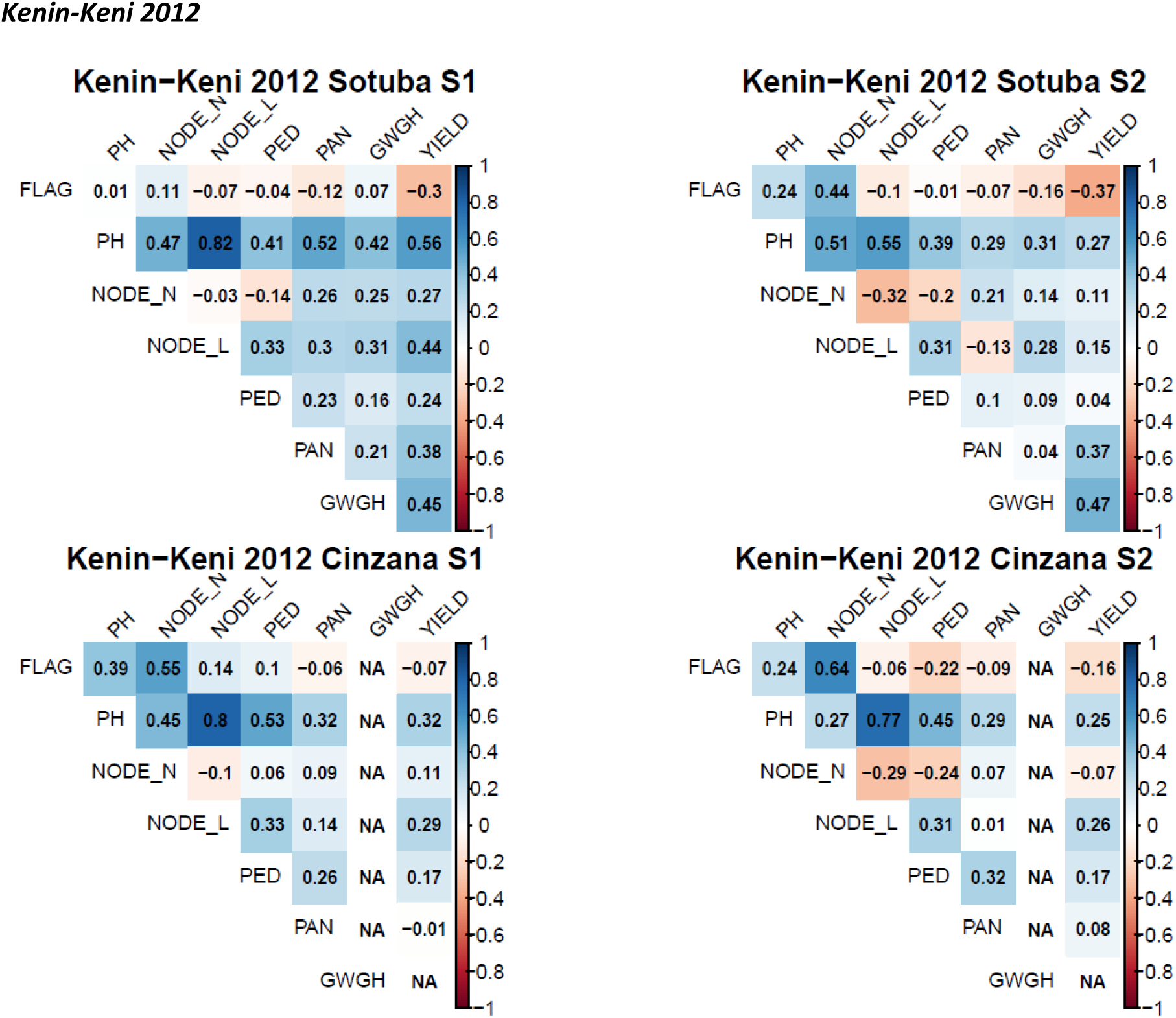
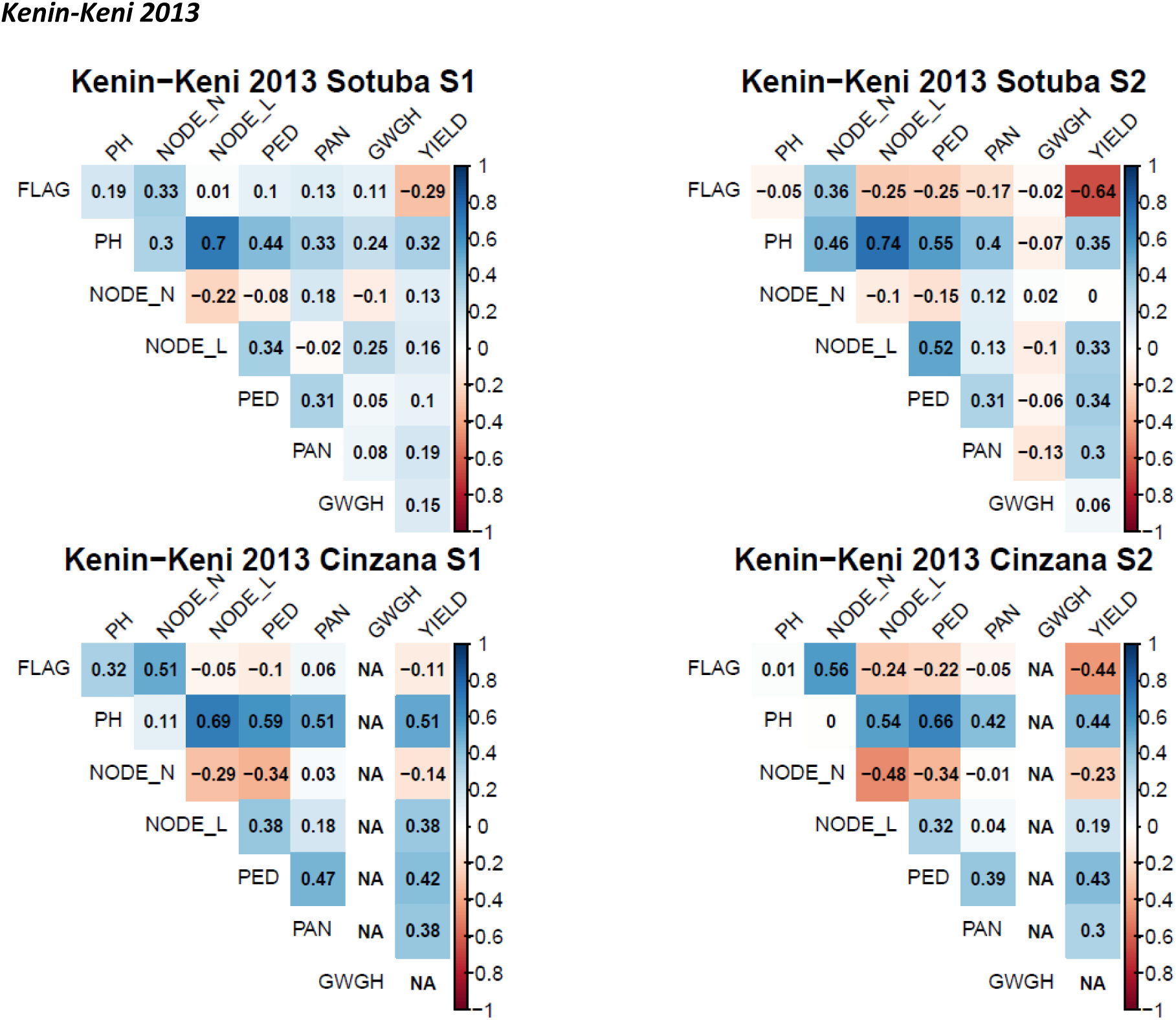
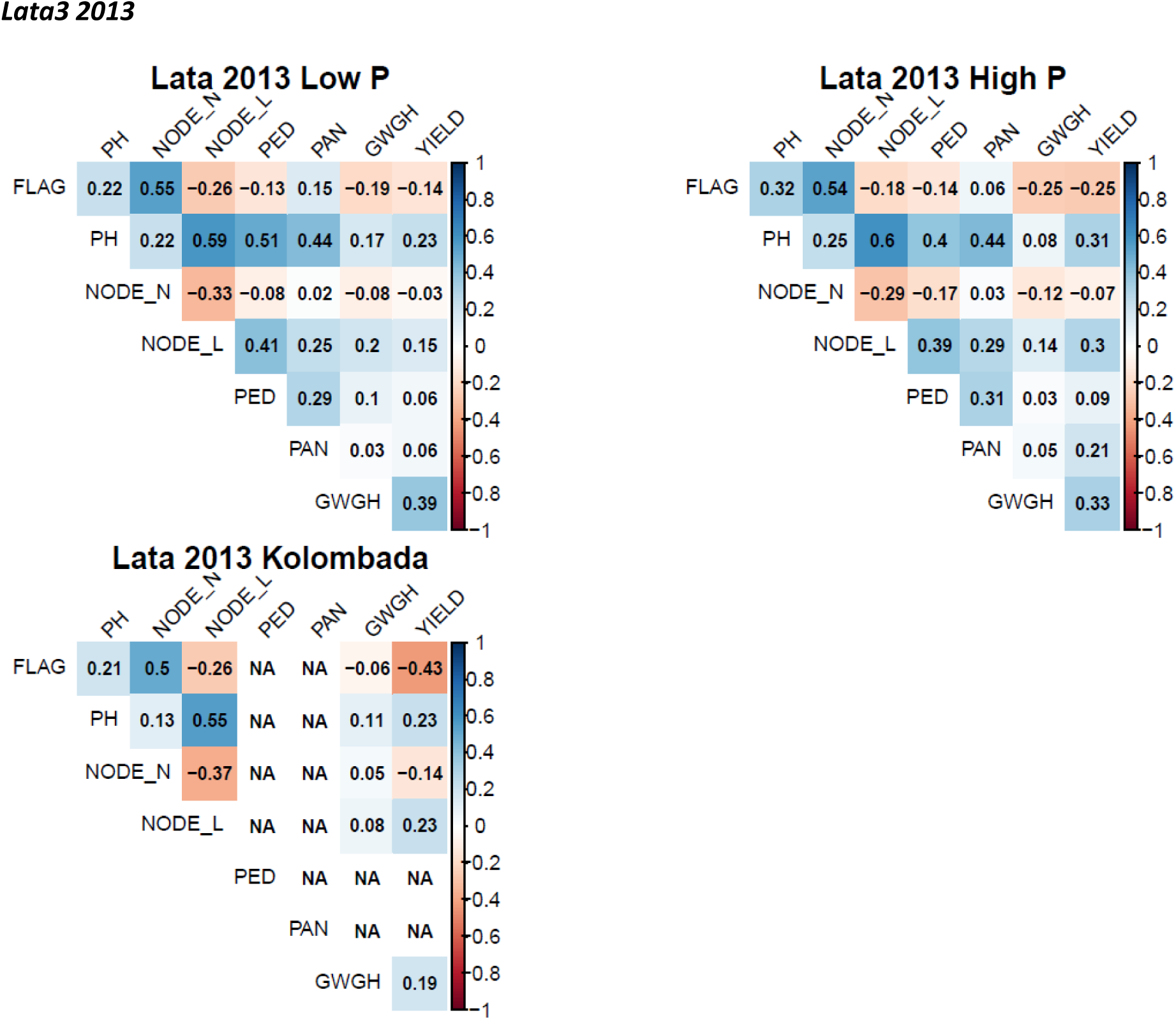
Pearson correlation matrix plots of the phenotypic traits for each population and year of phenotyping combination

**Figures S5:**
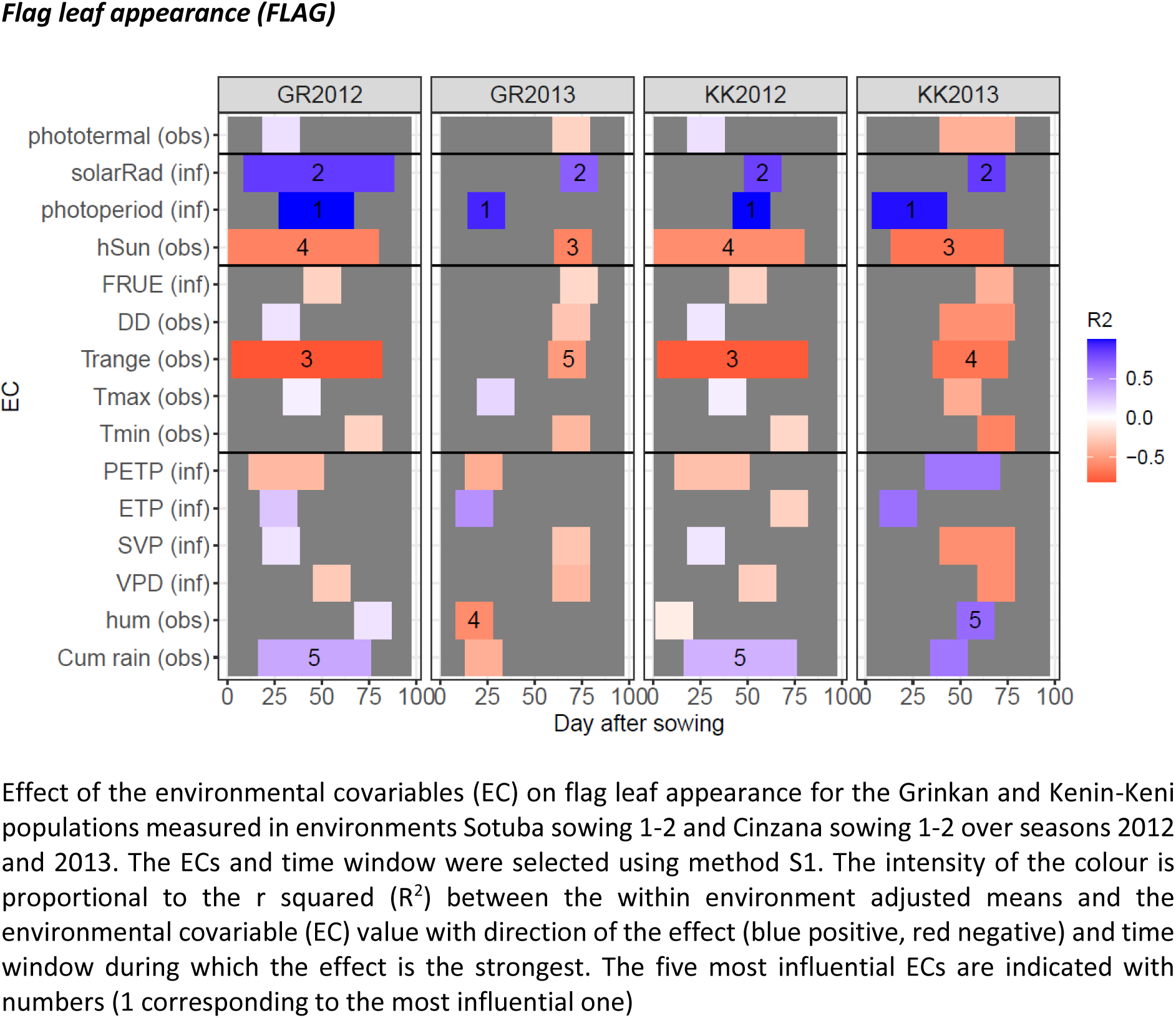
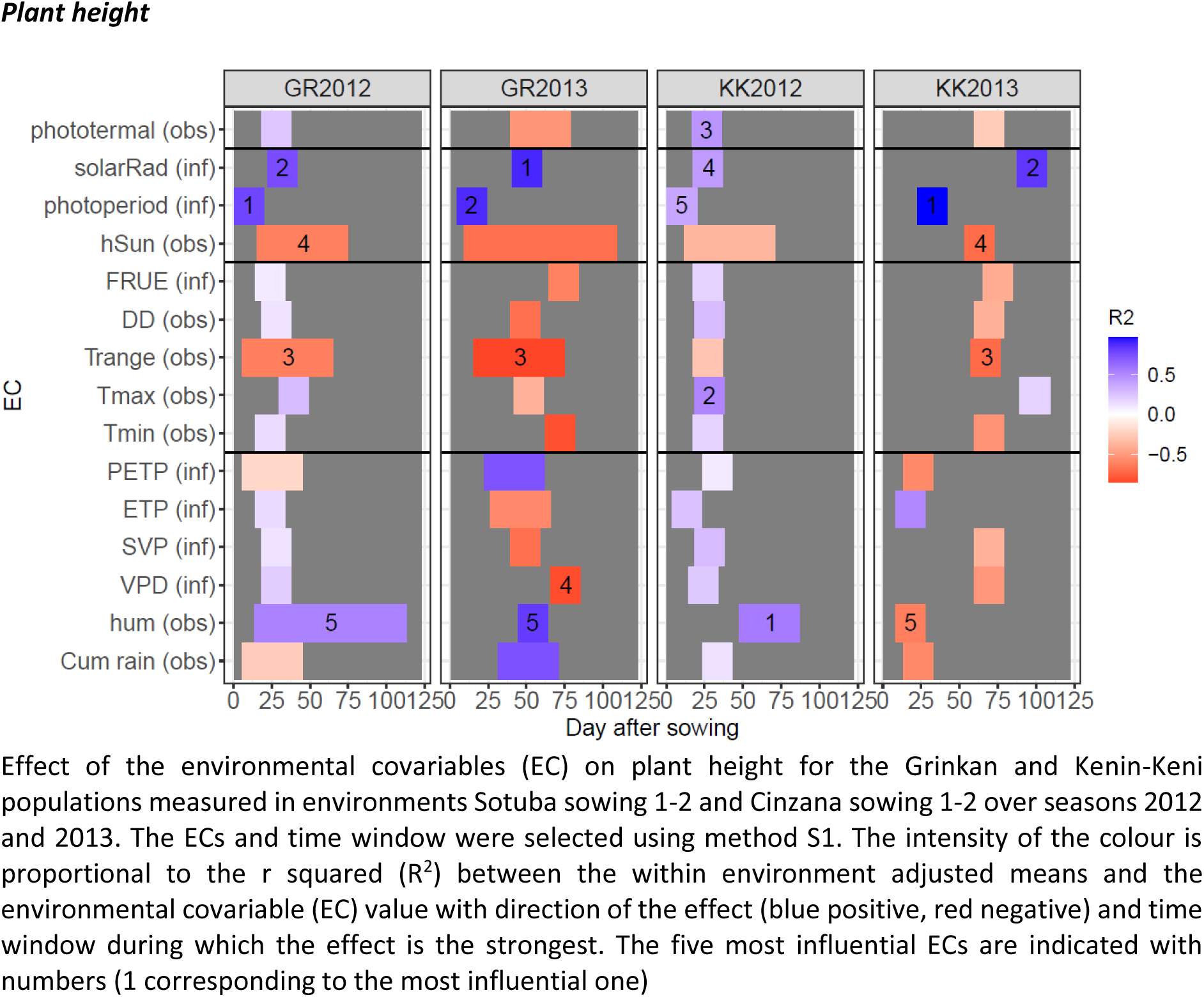
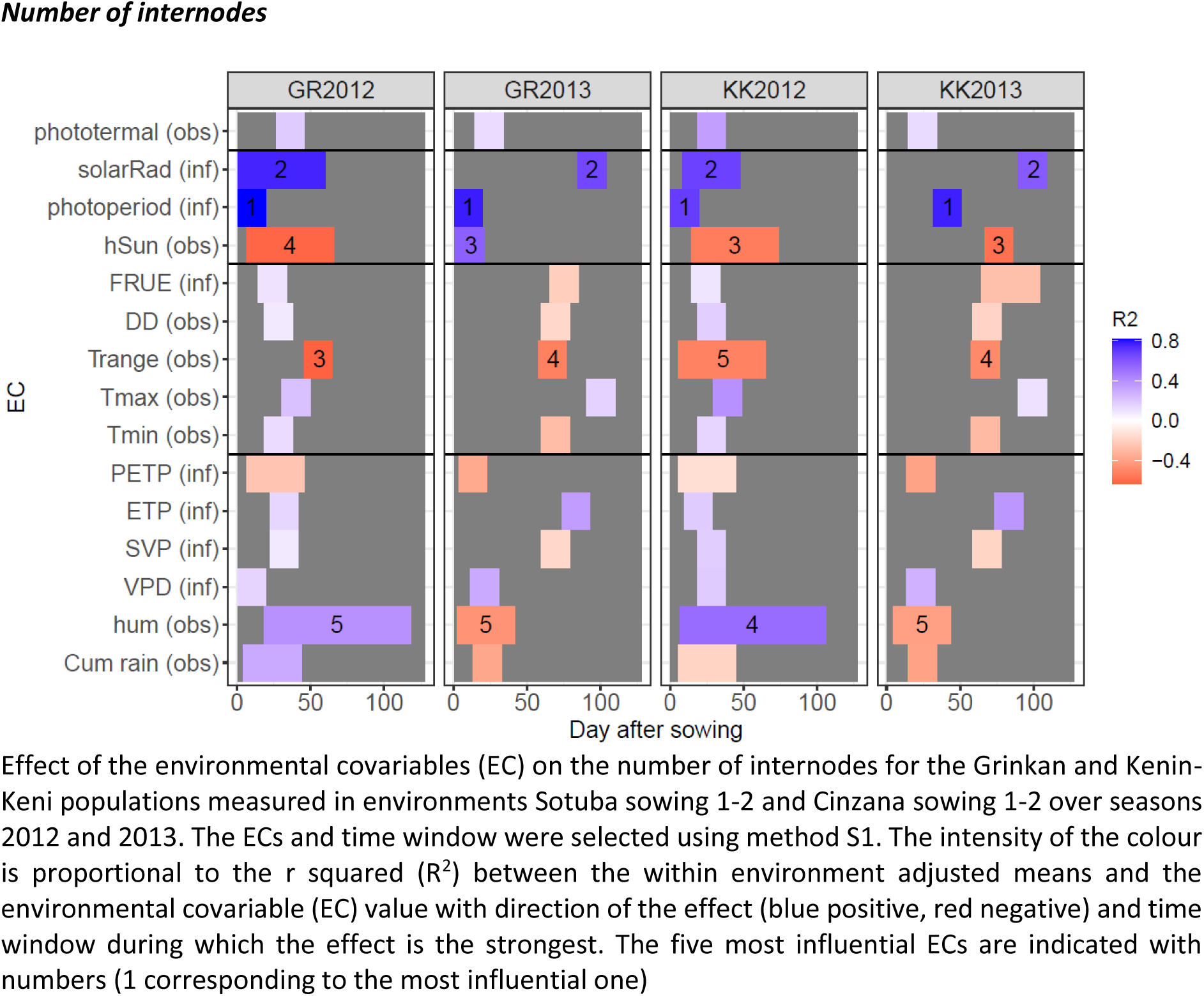
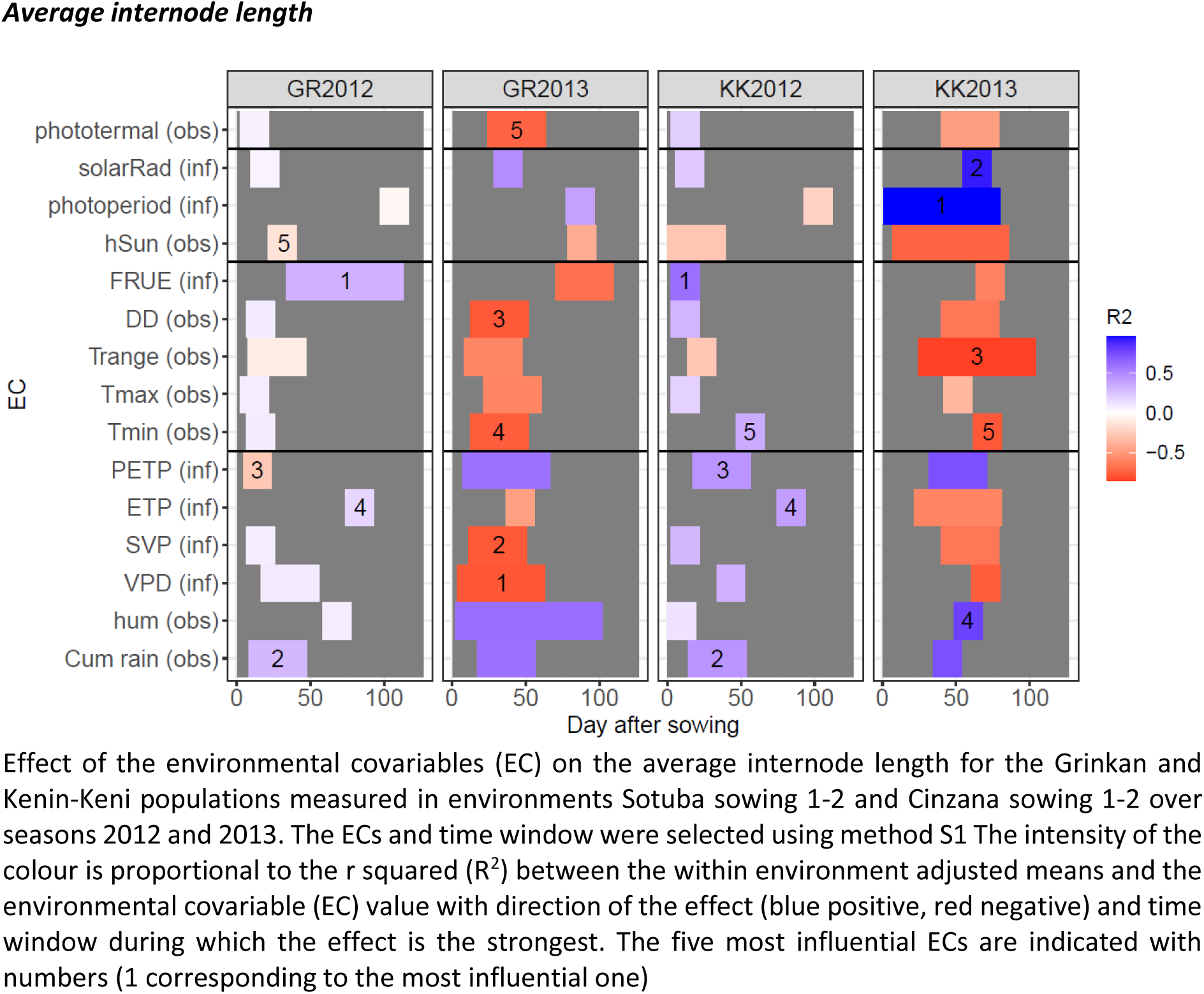
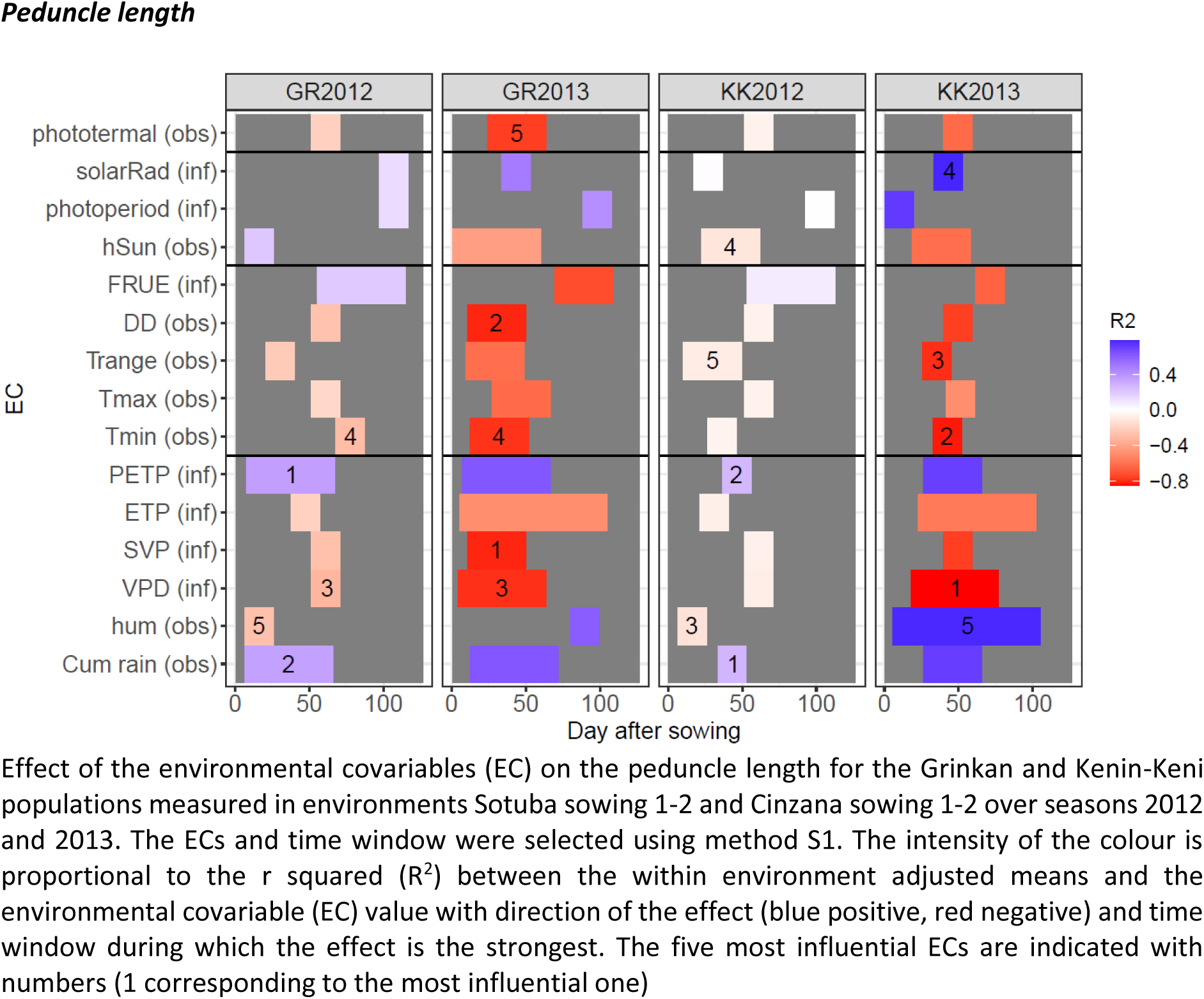
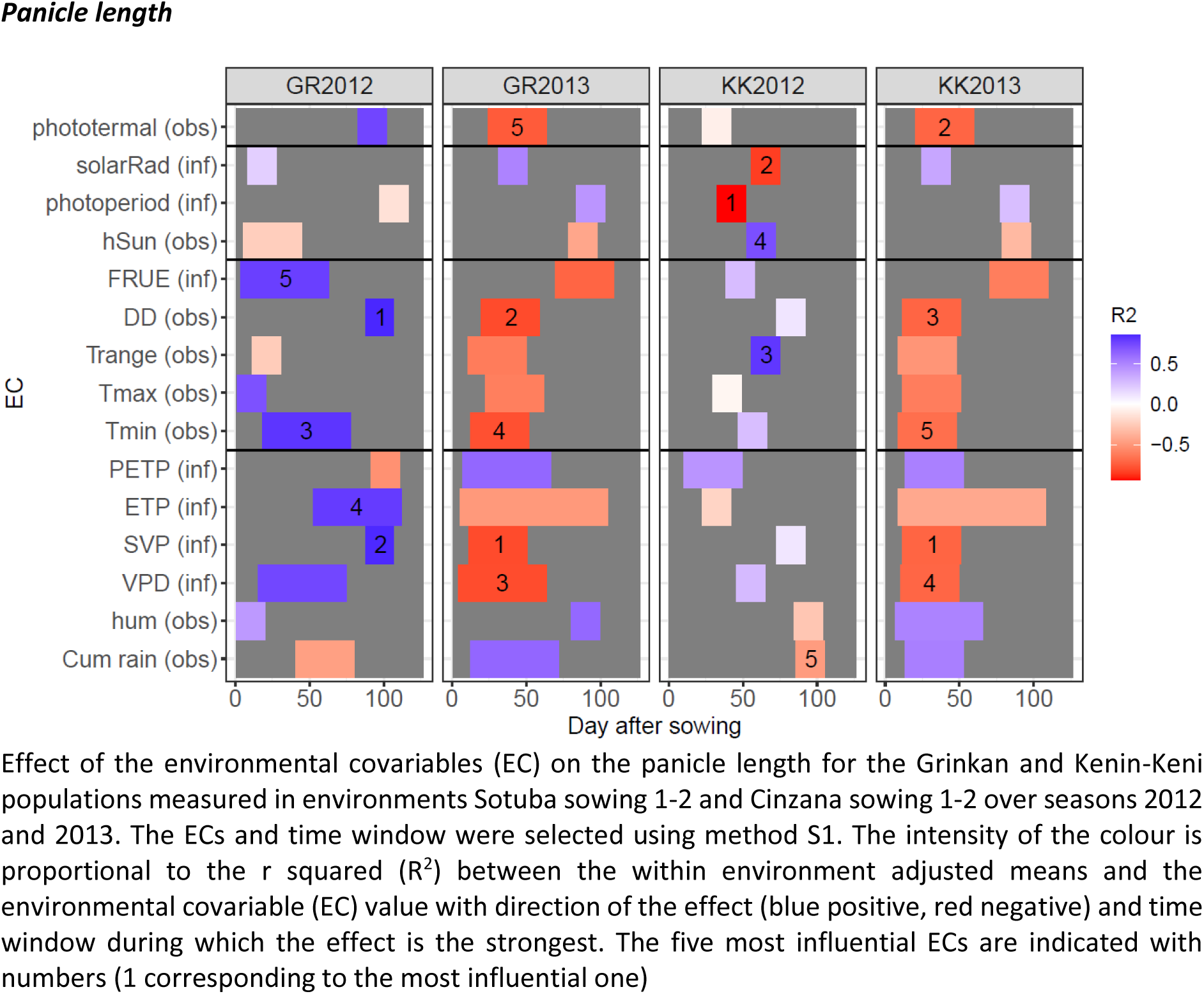
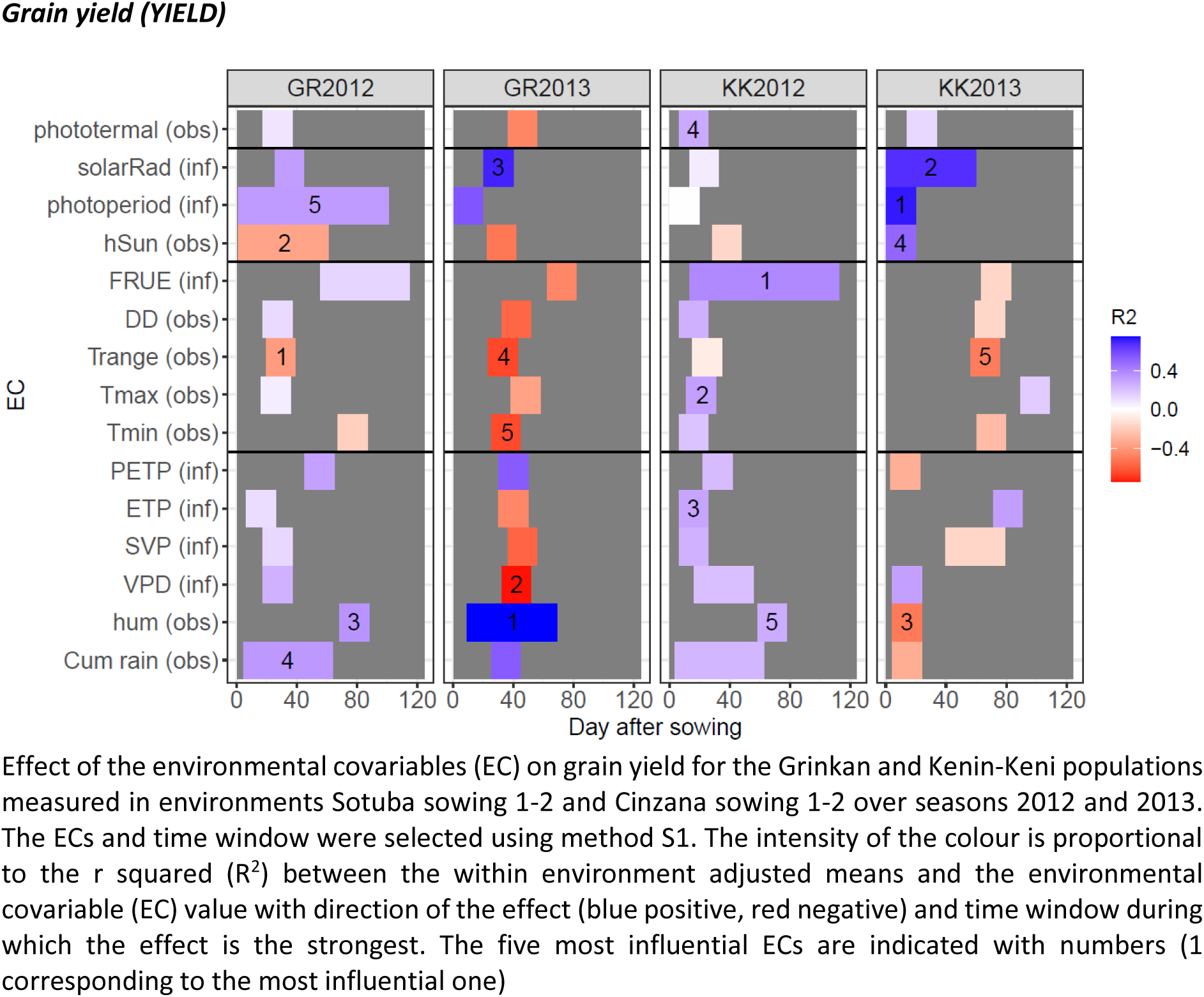
Phenotype by environmental covariables analysis visualisation

**Figures S6:**
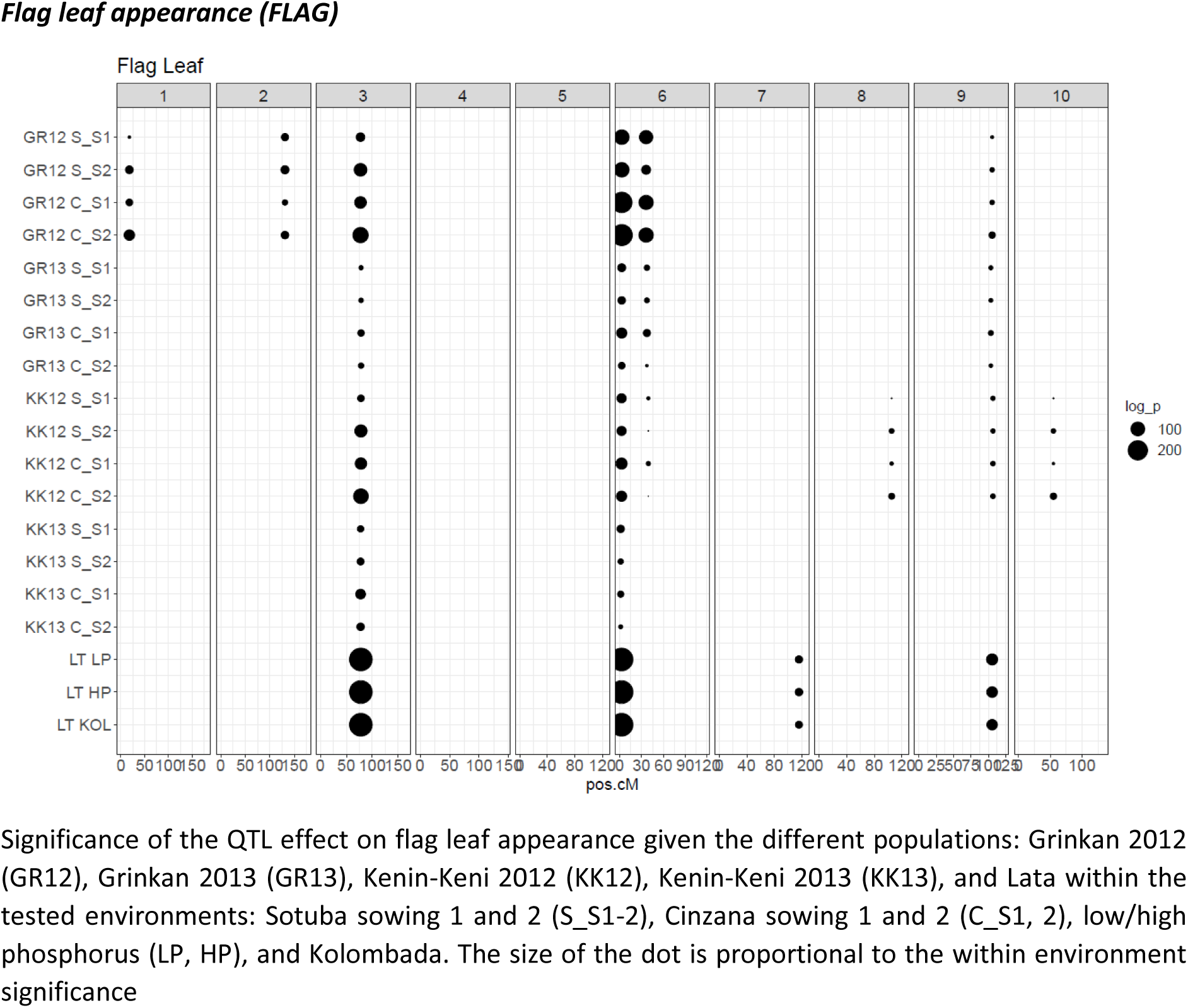
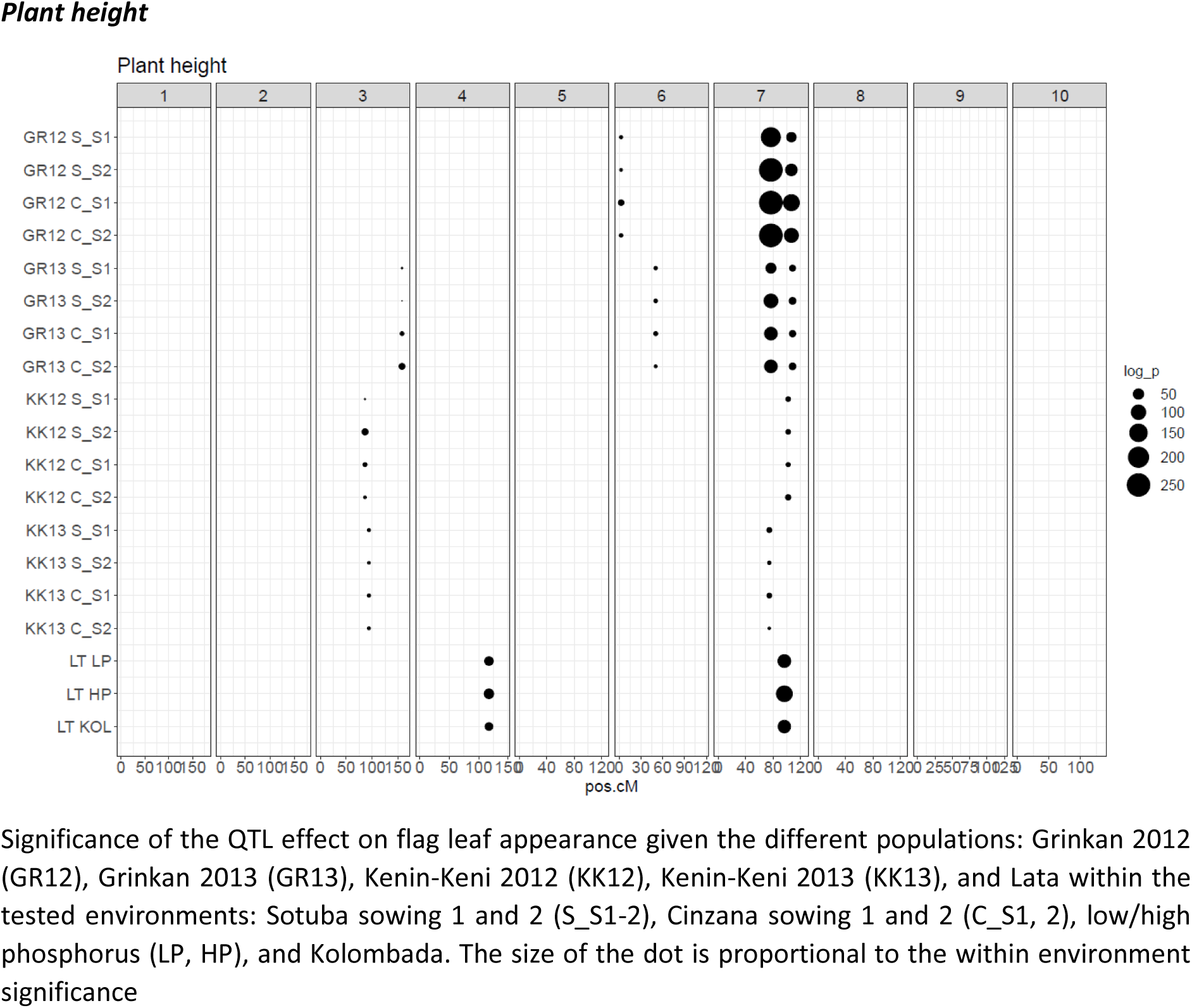
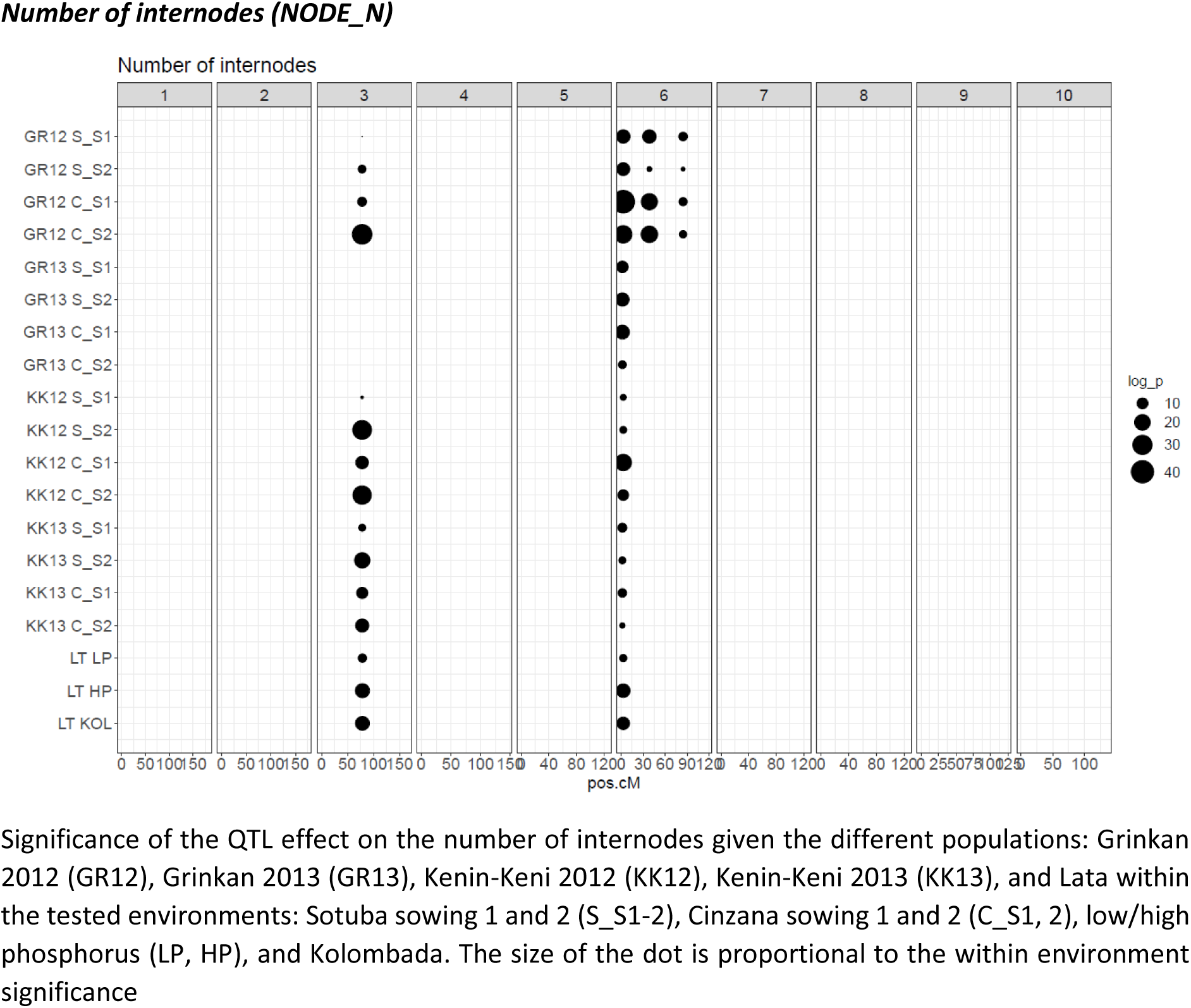
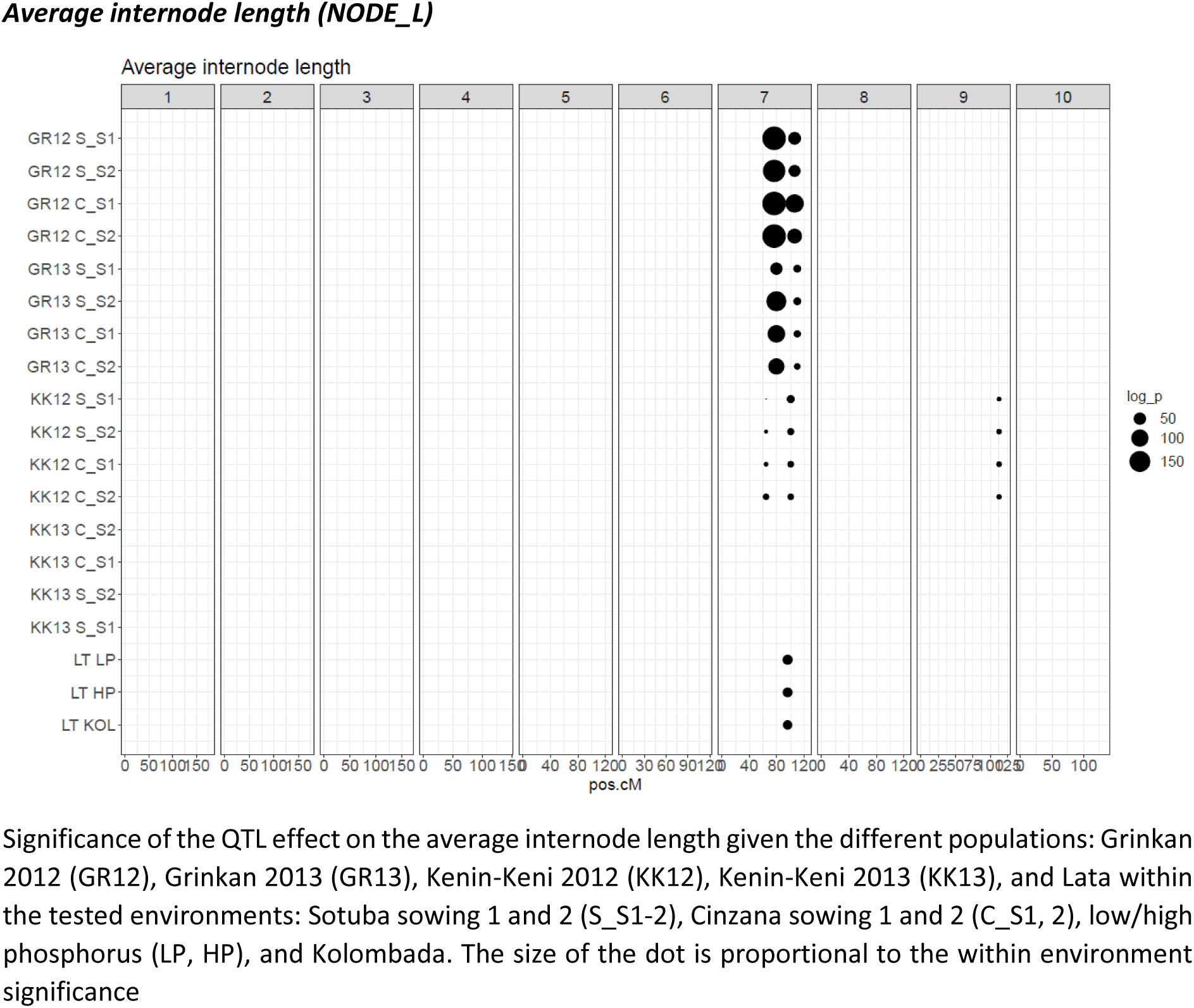
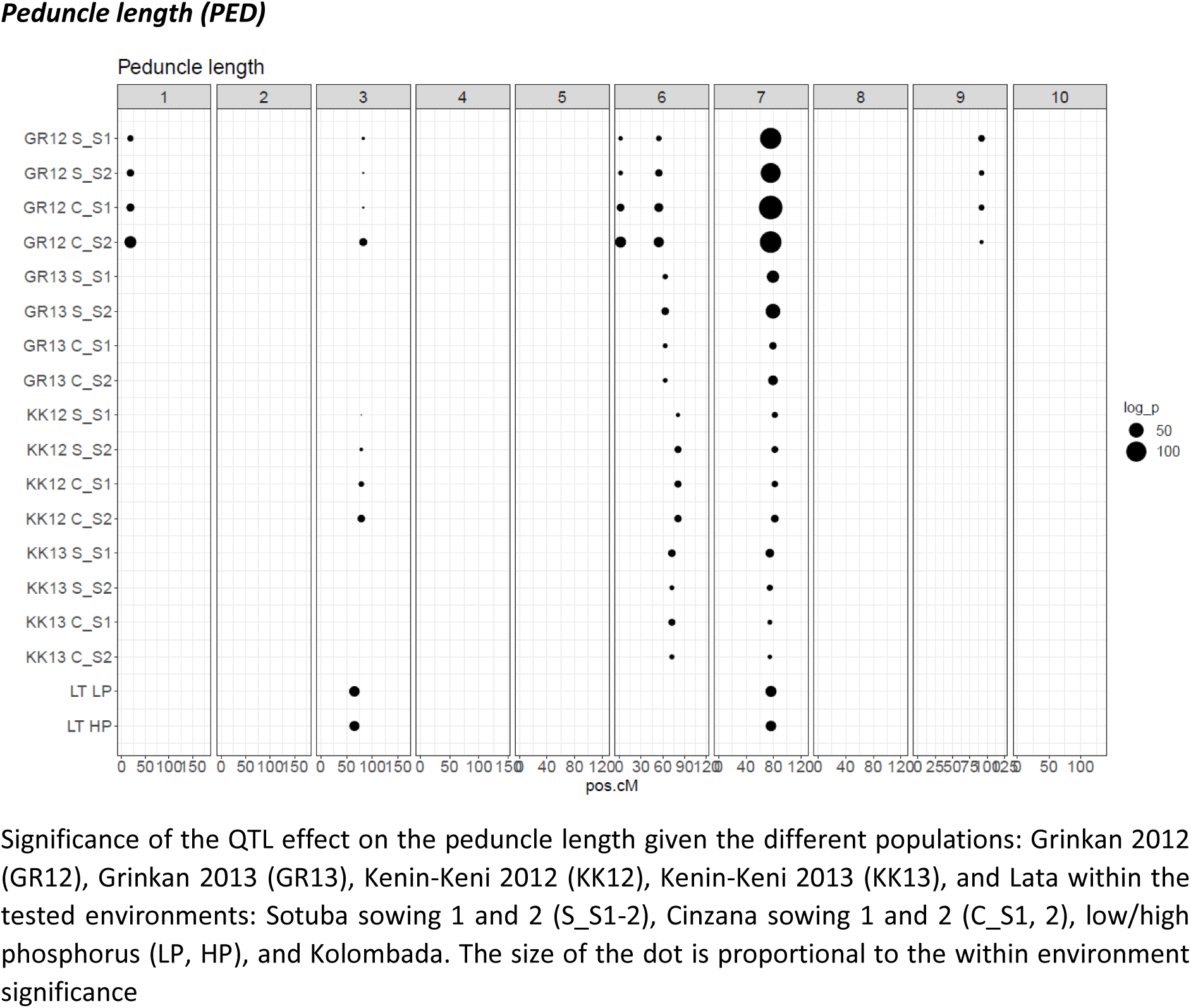
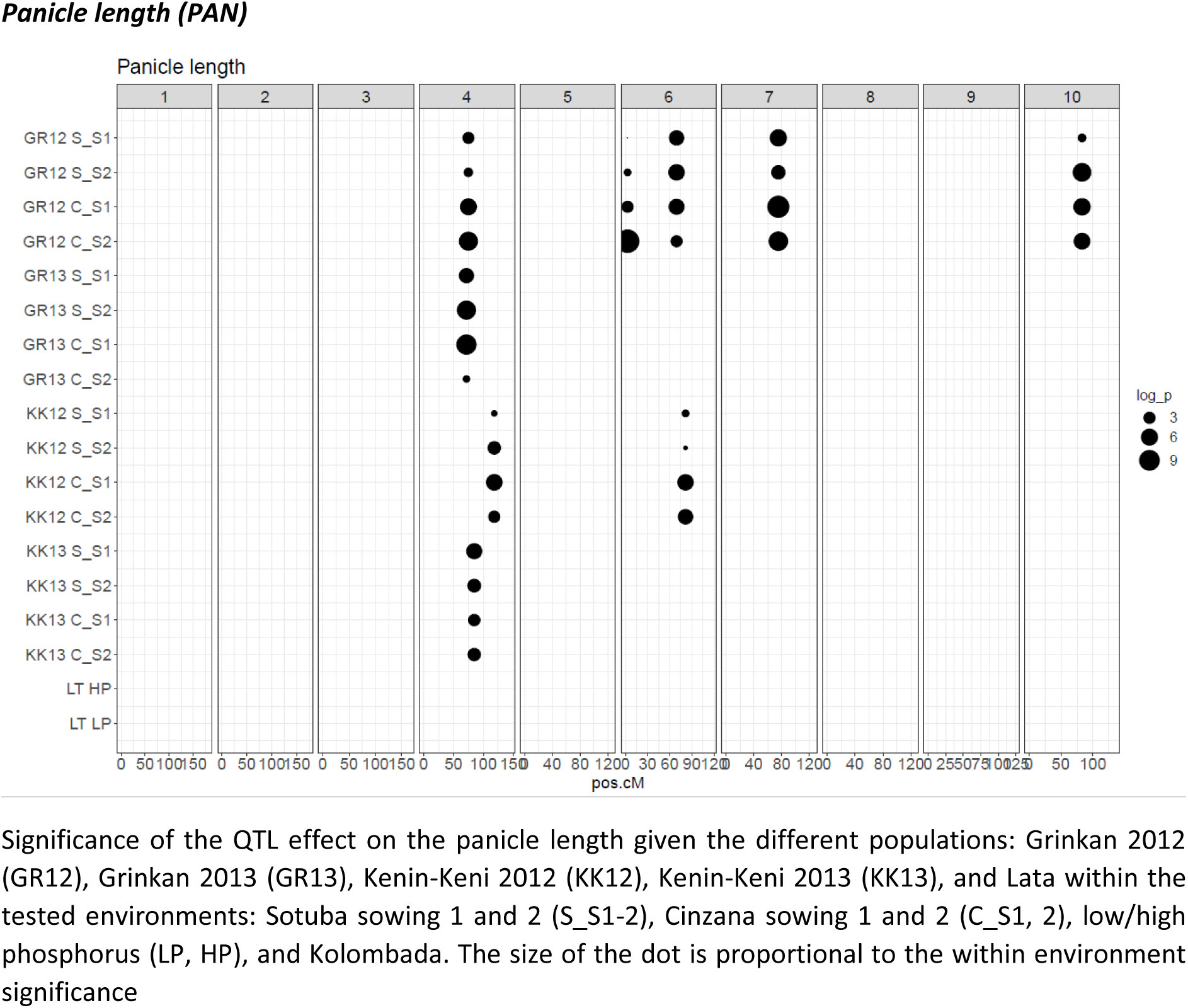
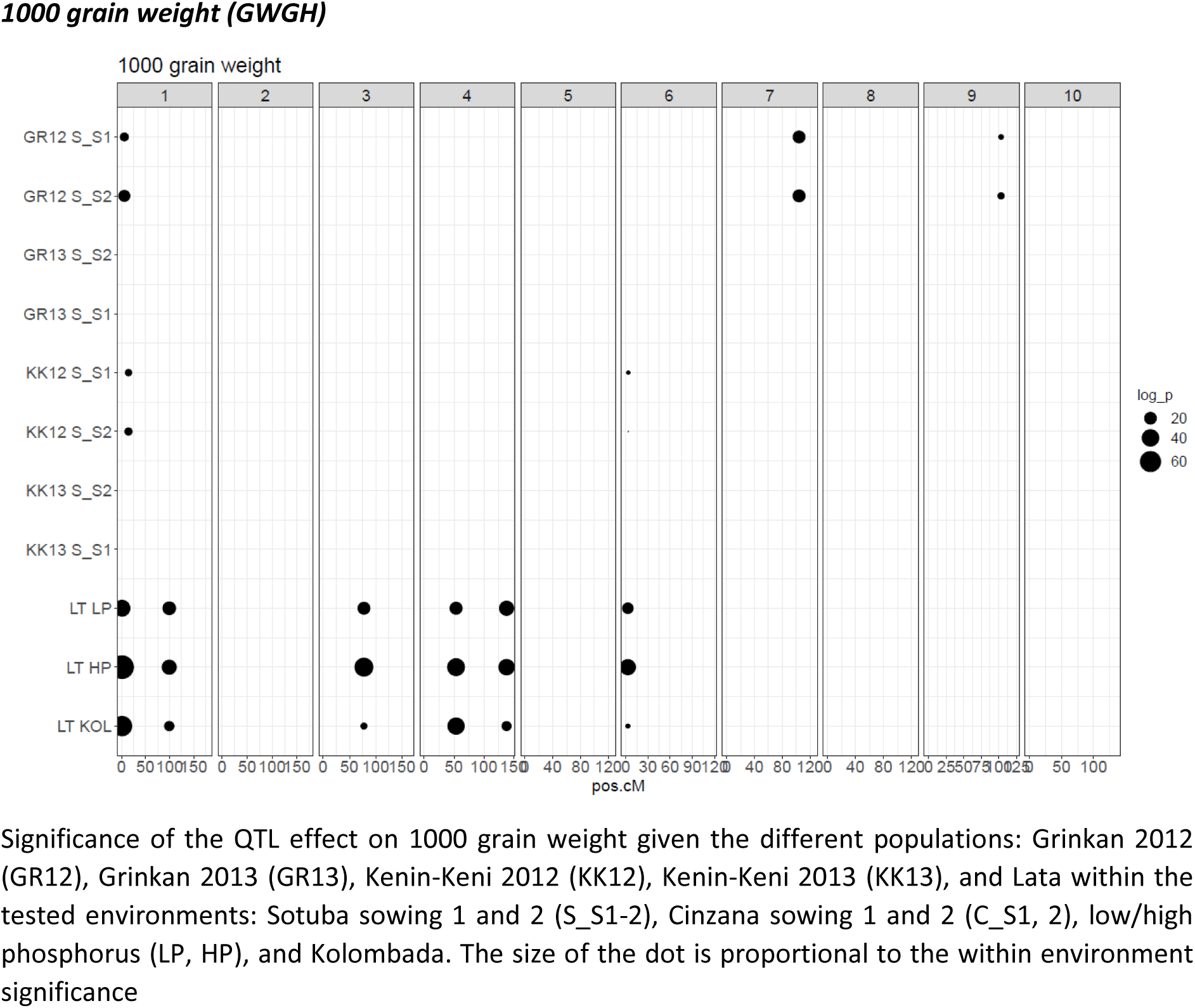
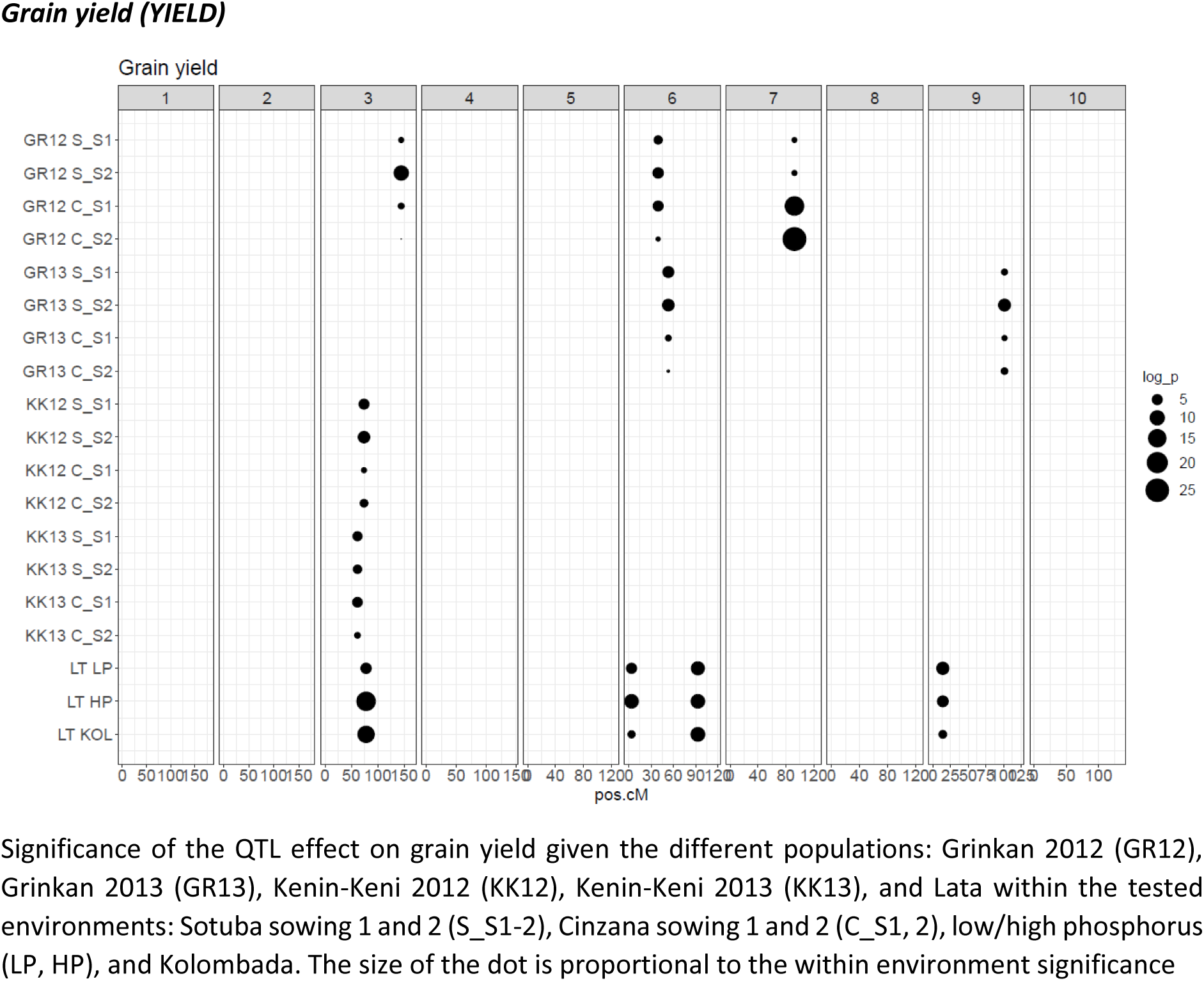
Effect plot of the detected QTLs over trait, populations and environments

**Figures S7:**
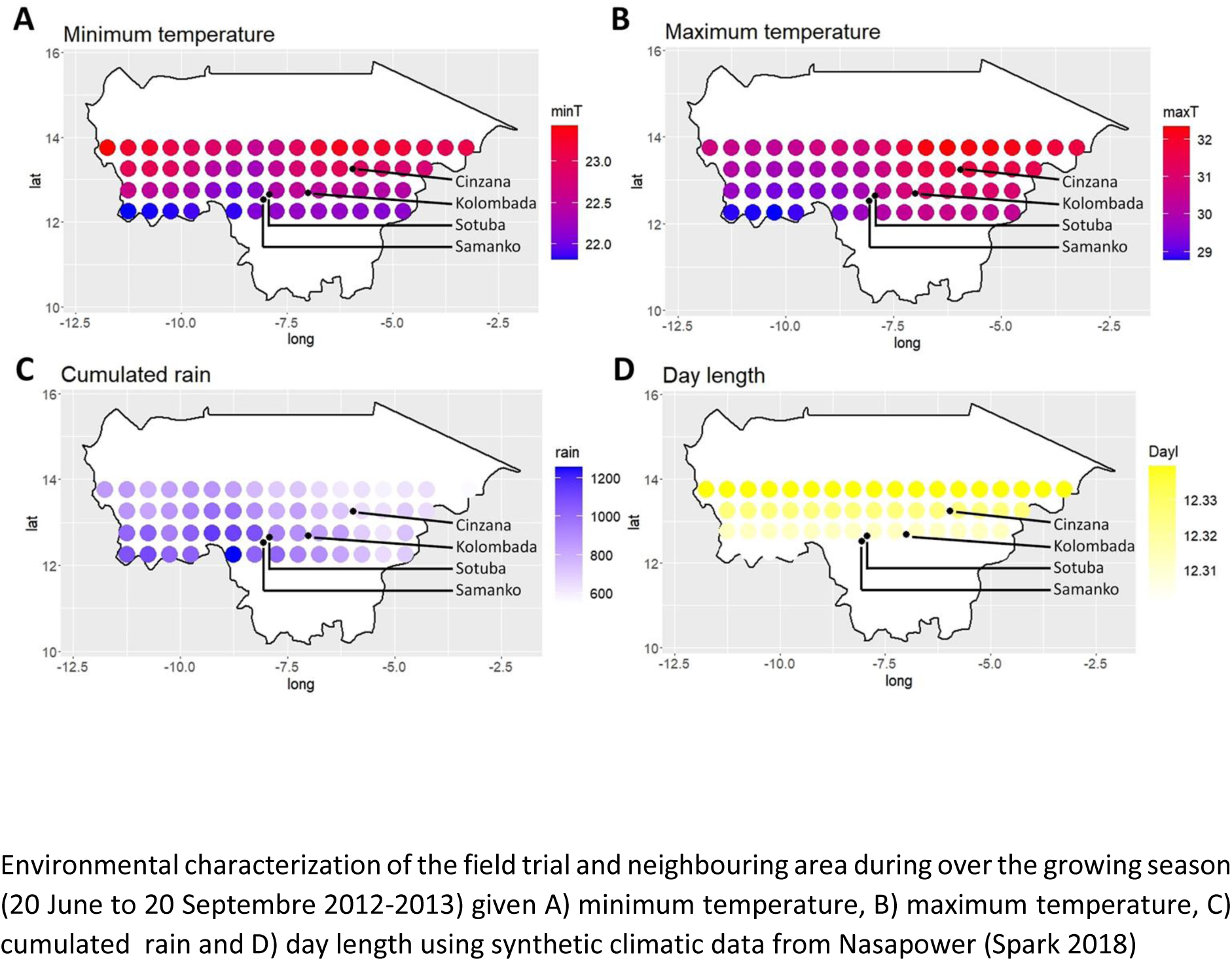
Field trial locations, sowing dates and environmental description

**Figures S8:**
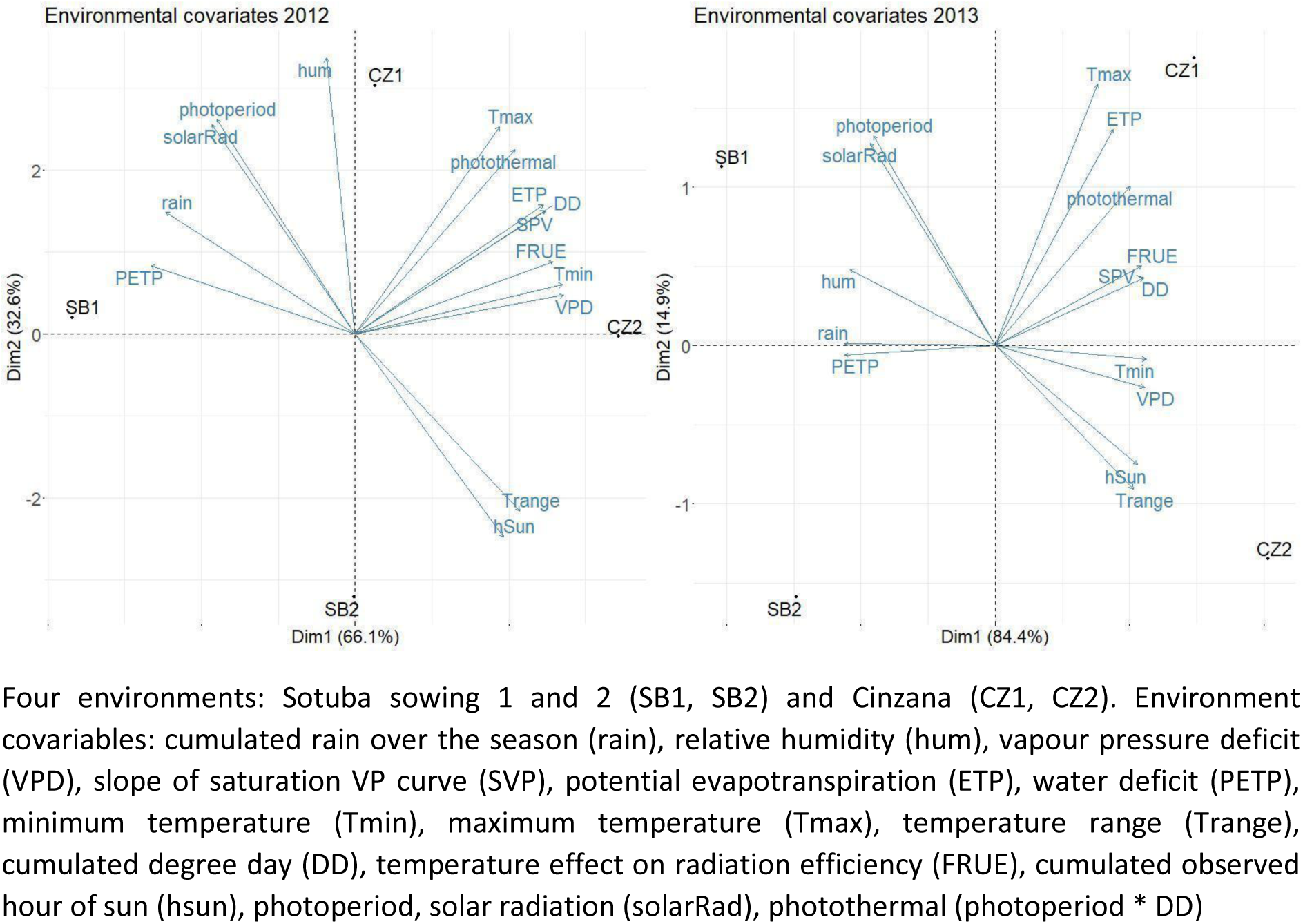
Cinzana and Sotuba environments principal component analysis based on ECs

## Supporting tables

**Table S1:** Consensus map statistics

| Chr | N. markers<sup>1</sup> | Length [cM] | N. CO<sup>2</sup> Grinkan | N. CO Kenin-Keni | N. CO. Lata3 |
| --- | --- | --- | --- | --- | --- |
| 1 | 8154 | 179.8 | 6562 | 2591 | 2570 |
| 2 | 6717 | 173.8 | 5470 | 2425 | 2158 |
| 3 | 7258 | 164.8 | 5571 | 2450 | 2534 |
| 4 | 5593 | 144.9 | 4887 | 2035 | 2183 |
| 5 | 3940 | 124.6 | 4577 | 2327 | 1960 |
| 6 | 4380 | 117.2 | 4548 | 2076 | 1908 |
| 7 | 3674 | 124.9 | 3894 | 1474 | 1574 |
| 8 | 3533 | 124 | 3895 | 1648 | 1733 |
| 9 | 4463 | 123.1 | 3967 | 1693 | 1784 |
| 10 | 3833 | 134.9 | 4298 | 1624 | 1716 |
| Total | 51545 | 1411.9 | 47669 | 20343 | 20120 |

1. Number of polymorphic markers in the consensus map 2. N. CO: total number of crossing over estimated in the different single reference BCNAM (Grinkan, Kenin-Keni, Lata3)

**Table S2:** Number of QTL detected for the traits and reference genotype by year combinations. Total R_2_ explained by the QTL are provided in parenthesis.

|  | FLAG | PH | NODE_N | NODE_L | PED | PAN | GWGH | YIELD |
| --- | --- | --- | --- | --- | --- | --- | --- | --- |
| Grinkan 2012 | 6 (48.9) | 3 (48) | 4 (17.6) | 2 (47.1) | 6 (31.2) | 5 (8.5) | 3 (13.5) | 3 (5.4) |
| Grinkan 2013 | 4 (32.1) | 4 (44.5) | 1 (9.6) | 2 (42.6) | 2 (21.1) | 1 (4.6) | 0 | 2 (7.3) |
| Kenin-Keni 2012 | 6 (53.4) | 2 (9.6) | 2 (16.8) | 3 (18.8) | 3 (14.3) | 2 (5.2) | 2 (7.9) | 1 (3.9) |
| Kenin-Keni 2013 | 2 (35.5) | 2 (12.7) | 2 (22.3) | 0 | 2 (16.9) | 1 (5.8) | 0 | 1 (5.9) |
| Lata3 | 4 (50.3) | 2 (20.1) | 2 (13.7) | 1 (11) | 2 (10.2) | 0 | 6 (30.4) | 4 (14.3) |

**Tables S3:** Within cross variance components: genotypic variance (σ_g_^2^) genotype by environment variance (σ_ge_^2^ ), error variance (σ_e_^2^) and heritability (h2) of the different populations. Empty cells correspond to traits that have not been evaluated in a specific population.

|  |  | Flag leaf [d] |  |  |  | Plant height [cm] |  |  |  | Peduncle length [cm] |  |  |  | Panicle length [cm] |  |  |  | Grain weight [g] |  |  |  | Grain yield [ton/ha] |  |  |  |
| --- | --- | --- | --- | --- | --- | --- | --- | --- | --- | --- | --- | --- | --- | --- | --- | --- | --- | --- | --- | --- | --- | --- | --- | --- | --- |
| **Grinkan** | N | Av | $\sigma_g^2$ | $\sigma_{ge}^2$ | $h^2$ | Av | $\sigma_g^2$ | $\sigma_{ge}^2$ | $h^2$ | Av | $\sigma_g^2$ | $\sigma_{ge}^2$ | $h^2$ | Av | $\sigma_g^2$ | $\sigma_{ge}^2$ | $h^2$ | Av | $\sigma_g^2$ | $\sigma_{ge}^2$ | $h^2$ | Av | $\sigma_g^2$ | $\sigma_{ge}^2$ | $h^2$ |
| Fara | 117 | 83.6 | 15.9 | 13.8 | 0.82 | 175.7 | 1360.1 | 662.6 | 0.89 | 36.8 | 23.2 | 33.3 | 0.74 | 30.7 | 8.7 | 22.6 | 0.61 | 25.3 | 3.2 | 3.1 | 0.67 | 1.8 | 0.3 | 0.9 | 0.55 |
| E36-1 | 125 | 76.3 | 35.0 | 14.7 | 0.91 | 166.6 | 738.3 | 645.0 | 0.82 | 34.6 | 20.7 | 35.1 | 0.7 | 28.0 | 4.3 | 11.3 | 0.6 | 25.4 | 2.9 | 2.8 | 0.68 | 2.5 | 0.2 | 1.1 | 0.41 |
| IS15401 | 113 | 80.4 | 8.0 | 7.8 | 0.8 | 166.8 | 615.9 | 624.3 | 0.8 | 34.3 | 22.0 | 33.6 | 0.72 | 28.9 | 3.4 | 14.9 | 0.48 | 23.5 | 3.9 | 2.5 | 0.76 | 2.0 | 0.2 | 0.8 | 0.44 |
| IS23540 | 61 | 77.8 | 15.9 | 9.4 | 0.87 | 176.3 | 1098.6 | 641.6 | 0.87 | 40.2 | 25.4 | 26.5 | 0.79 | 28.3 | 1.6 | 12.7 | 0.34 | 24.4 | 5.0 | 2.2 | 0.82 | 2.3 | 0.0 | 1.3 | 0.04 |
| B35 | 118 | 79.1 | 25.4 | 19.9 | 0.84 | 156.7 | 888.6 | 489.9 | 0.88 | 38.1 | 36.4 | 36.0 | 0.8 | 27.6 | 5.7 | 13.0 | 0.64 | 23.7 | 4.6 | 1.4 | 0.86 | 1.5 | 0.2 | 0.5 | 0.6 |
| Kono. | 134 | 78.9 | 6.8 | 6.2 | 0.81 | 180.7 | 1837.9 | 823.9 | 0.9 | 35.0 | 36.8 | 25.5 | 0.85 | 29.1 | 8.0 | 12.7 | 0.72 | 26.8 | 6.1 | 3.7 | 0.77 | 2.1 | 0.2 | 0.9 | 0.44 |
| SC566-14 | 142 | 74.5 | 53.5 | 23.9 | 0.9 | 173.2 | 1335.7 | 564.3 | 0.9 | 39.7 | 60.6 | 39.5 | 0.86 | 28.5 | 5.5 | 16.5 | 0.57 | 24.1 | 2.4 | 2.0 | 0.7 | 2.0 | 0.2 | 0.7 | 0.46 |
| Framida | 140 | 76.2 | 19.5 | 6.5 | 0.92 | 172.6 | 1077.2 | 445.4 | 0.91 | 36.5 | 24.3 | 19.4 | 0.83 | 28.6 | 6.8 | 9.5 | 0.74 | 24.5 | 2.0 | 3.6 | 0.52 | 2.2 | 0.1 | 0.7 | 0.35 |
| CSM417 | 160 | 75.9 | 16.8 | 5.8 | 0.92 | 179.1 | 1974.4 | 946.5 | 0.89 | 40.4 | 45.2 | 35.9 | 0.83 | 30.8 | 5.1 | 22.3 | 0.48 |  |  |  |  | 2.1 | 0.2 | 0.9 | 0.41 |
| CSM63 | 67 | 75.3 | 28.8 | 15.0 | 0.89 | 187.1 | 1697.7 | 1250.4 | 0.84 | 41.4 | 57.0 | 32.2 | 0.88 | 31.6 | 3.9 | 23.8 | 0.4 | 23.1 | 1.6 | 4.0 | 0.44 | 1.9 | 0.0 | 1.0 | 0 |
| CSM388 | 149 | 76.6 | 24.1 | 4.8 | 0.95 | 201.7 | 1779.5 | 1283.5 | 0.85 | 40.5 | 30.6 | 41.5 | 0.75 | 32.0 | 5.2 | 21.0 | 0.5 | 23.2 | 1.7 | 3.1 | 0.52 | 2.2 | 0.1 | 1.0 | 0.28 |
| Gadiaba | 135 | 80.1 | 9.5 | 4.6 | 0.89 | 192.6 | 2466.6 | 1211.0 | 0.89 | 38.3 | 41.8 | 42.0 | 0.8 | 27.5 | 7.7 | 11.1 | 0.74 | 23.7 | 2.7 | 2.6 | 0.67 | 2.2 | 0.1 | 0.7 | 0.35 |
| Wth Kaur | 137 | 79.1 | 12.4 | 6.2 | 0.89 | 171.8 | 1780.4 | 663.7 | 0.91 | 39.5 | 87.6 | 31.2 | 0.92 | 28.7 | 4.6 | 12.8 | 0.59 | 24.3 | 4.7 | 3.7 | 0.72 | 2.2 | 0.2 | 0.7 | 0.54 |
| Average | 123 | 78.0 | 20.9 | 10.6 | 0.88 | 177.0 | 1434.7 | 788.6 | 0.87 | 38.1 | 39.3 | 33.2 | 0.81 | 29.3 | 5.4 | 15.7 | 0.57 | 24.3 | 3.4 | 2.9 | 0.68 | 2.1 | 0.1 | 0.9 | 0.37 |

***Kenin-Keni part characterized in 2012***
|  |  | Flag leaf [d] |  |  |  | Plant height [cm] |  |  |  | Peduncle length [cm] |  |  |  | Panicle length [cm] |  |  |  | Grain weight [g] |  |  |  | Grain yield [ton/ha] |  |  |  |
| --- | --- | --- | --- | --- | --- | --- | --- | --- | --- | --- | --- | --- | --- | --- | --- | --- | --- | --- | --- | --- | --- | --- | --- | --- | --- |
| ***Keni-Keni*** | N | Av | $\sigma_g^2$ | $\sigma_{ge}^2$ | $h^2$ | Av | $\sigma_g^2$ | $\sigma_{ge}^2$ | $h^2$ | Av | $\sigma_g^2$ | $\sigma_{ge}^2$ | $h^2$ | Av | $\sigma_g^2$ | $\sigma_{ge}^2$ | $h^2$ | Av | $\sigma_g^2$ | $\sigma_{ge}^2$ | $h^2$ | Av | $\sigma_g^2$ | $\sigma_{ge}^2$ | $h^2$ |
| Fara | 134 | 79.1 | 34.0 | 7.0 | 0.95 | 245.7 | 705.2 | 689.7 | 0.8 | 59.7 | 20.1 | 55.4 | 0.59 | 37.3 | 3.5 | 26.7 | 0.34 | 22.9 | 3.8 | 3.0 | 0.72 | 2.0 | 0.3 | 0.7 | 0.6 |
| E36-1 | 144 | 70.0 | 37.8 | 11.1 | 0.93 | 207.9 | 1044.0 | 758.4 | 0.85 | 51.3 | 69.0 | 43.0 | 0.87 | 35.1 | 7.9 | 23.6 | 0.57 | 21.0 | 2.8 | 2.2 | 0.71 | 2.1 | 0.1 | 0.7 | 0.39 |
| IS15401 | 138 | 75.0 | 23.3 | 9.0 | 0.91 | 223.1 | 642.9 | 577.3 | 0.82 | 55.7 | 25.2 | 45.3 | 0.69 | 35.6 | 4.3 | 29.6 | 0.37 | 22.7 | 2.3 | 2.5 | 0.65 | 2.2 | 0.1 | 0.8 | 0.27 |
| B35 | 79 | 71.6 | 17.0 | 19.3 | 0.78 | 205.4 | 898.9 | 1324.1 | 0.73 | 55.1 | 51.4 | 61.6 | 0.77 | 36.6 | 3.4 | 39.4 | 0.26 | 20.3 | 4.0 | 4.2 | 0.65 | 1.7 | 0.2 | 0.4 | 0.65 |
| CSM417 | 80 | 71.4 | 1.7 | 6.3 | 0.52 | 241.7 | 441.5 | 1040.8 | 0.63 | 64.4 | 10.1 | 44.8 | 0.47 | 40.0 | 5.7 | 41.4 | 0.36 |  |  |  |  | 2.4 | 0.0 | 1.1 | 0 |
| Average | 115 | 73.4 | 22.8 | 10.5 | 0.82 | 224.8 | 746.5 | 878.1 | 0.77 | 57.2 | 35.2 | 50.0 | 0.68 | 36.9 | 5.0 | 32.1 | 0.38 | 21.7 | 3.2 | 3.0 | 0.68 | 2.1 | 0.1 | 0.8 | 0.38 |

**Grinkan part characterized in 2013**
|  |  | Flag leaf [d] |  |  |  | Plant height [cm] |  |  |  | Peduncle length [cm] |  |  |  | Panicle length [cm] |  |  |  | Grain weight [g] |  |  |  | Grain yield [ton/ha] |  |  |  |
| --- | --- | --- | --- | --- | --- | --- | --- | --- | --- | --- | --- | --- | --- | --- | --- | --- | --- | --- | --- | --- | --- | --- | --- | --- | --- |
| | N | Av | $\sigma_g^2$ | $\sigma_{ge}^2$ | $h^2$ | Av | $\sigma_g^2$ | $\sigma_{ge}^2$ | $h^2$ | Av | $\sigma_g^2$ | $\sigma_{ge}^2$ | $h^2$ | Av | $\sigma_g^2$ | $\sigma_{ge}^2$ | $h^2$ | Av | $\sigma_g^2$ | $\sigma_{ge}^2$ | $h^2$ | Av | $\sigma_g^2$ | $\sigma_{ge}^2$ | $h^2$ |
| E36-1 | 46 | 70.2 | 31.7 | 7.1 | 0.95 | 155.29 | 395.35 | 262 | 0.86 | 31.6 | 13.8 | 15.4 | 0.78 | 27.1 | 5.07 | 5.83 | 0.78 |  |  |  |  | 1.9 | 0.04 | 0.42 | 0.29 |
| V33/08 | 67 | 73.1 | 6.5 | 6.1 | 0.81 | 157.43 | 655.28 | 555.2 | 0.83 | 36.1 | 35.06 | 44.9 | 0.76 | 29.9 | 7.63 | 9.1 | 0.77 |  |  |  |  | 1.5 | 0.05 | 0.46 | 0.29 |
| Kalaban | 115 | 70.4 | 19.2 | 7.1 | 0.92 | 153.5 | 461.43 | 318.5 | 0.85 | 35.2 | 11.75 | 28.9 | 0.62 | 26.7 | 2.02 | 9.09 | 0.47 |  |  |  |  | 1.68 | 0.05 | 0.44 | 0.31 |
| Malisor | 95 | 72.3 | 14.7 | 17.6 | 0.77 | 142.77 | 139.03 | 259.7 | 0.68 | 31.9 | 18.73 | 19.4 | 0.79 | 26.7 | 2.53 | 8.9 | 0.53 | 17.1 | 5.6 | 42.1 | 0.21 | 1.68 | 0.02 | 0.41 | 0.18 |
| BimbG | 100 | 79 | 15.2 | 10.9 | 0.85 | 177.86 | 1001.7 | 603.9 | 0.87 | 35.9 | 23.85 | 28.3 | 0.77 | 30.5 | 10.9 | 10.44 | 0.81 | 19.3 | 16.2 | 23.4 | 0.58 | 1.09 | 0.22 | 0.35 | 0.71 |
| Hafijega | 84 | 71.6 | 25.2 | 10.3 | 0.91 | 165.26 | 1310.7 | 614.2 | 0.9 | 35.7 | 34.89 | 26.8 | 0.84 | 31.2 | 9.57 | 11.31 | 0.77 |  |  |  |  | 1.27 | 0.1 | 0.38 | 0.53 |
| CSM_388 | 47 | 69.5 | 24.5 | 6.5 | 0.94 | 185.67 | 1540.3 | 661 | 0.9 | 37.9 | 44.31 | 27.7 | 0.86 | 30.1 | 5.53 | 8.04 | 0.73 |  |  |  |  | 1.73 | 0.07 | 0.57 | 0.33 |
| Sh Kaur | 50 | 73.9 | 5.1 | 5.6 | 0.79 | 161.13 | 843.1 | 618.5 | 0.85 | 34.8 | 35.49 | 41.8 | 0.77 | 29 | 3.46 | 12.68 | 0.52 |  |  |  |  | 1.39 | 0.19 | 0.45 | 0.63 |
| average | 75.5 | 72.5 |  |  | 0.87 | 162.36 |  |  | 0.84 | 34.9 |  |  | 0.77 | 28.9 |  |  | 0.67 | 18.2 |  |  | 0.39 | 1.53 |  |  | 0.41 |

***Kenin-Keni part characterized in 2013***
|  |  | Flag leaf [d] |  |  |  | Plant height [cm] |  |  |  | Peduncle length [cm] |  |  |  | Panicle length [cm] |  |  |  | Grain weight [g] |  |  |  | Grain yield [ton/ha] |  |  |  |
| --- | --- | --- | --- | --- | --- | --- | --- | --- | --- | --- | --- | --- | --- | --- | --- | --- | --- | --- | --- | --- | --- | --- | --- | --- | --- |
| | N | Av | $\sigma_g^2$ | $\sigma_{ge}^2$ | $h^2$ | Av | $\sigma_g^2$ | $\sigma_{ge}^2$ | $h^2$ | Av | $\sigma_g^2$ | $\sigma_{ge}^2$ | $h^2$ | Av | $\sigma_g^2$ | $\sigma_{ge}^2$ | $h^2$ | Av | $\sigma_g^2$ | $\sigma_{ge}^2$ | $h^2$ | Av | $\sigma_g^2$ | $\sigma_{ge}^2$ | $h^2$ |
| Sangatigui | 35 | 68.2 | 28.3 | 8.7 | 0.93 | 213.36 | 477.75 | 358.7 | 0.84 | 57.2 | 17.01 | 50.2 | 0.58 | 39.99 | 4.97 | 15.58 | 0.56 |  |  |  |  | 1.49 | 0.14 | 0.46 | 0.55 |
| IS23540 | 67 | 70.2 | 25.1 | 12.7 | 0.89 | 210.72 | 297.61 | 430.5 | 0.73 | 48.7 | 40.06 | 41.4 | 0.79 | 34.84 | 6.72 | 12.46 | 0.68 |  |  |  |  | 1.54 | 0.13 | 0.44 | 0.54 |
| Kalaban | 80 | 67.2 | 16.7 | 8.6 | 0.89 | 196 | 928.98 | 367.6 | 0.91 | 50.2 | 47.95 | 42.3 | 0.82 | 34.79 | 10.6 | 13.87 | 0.75 |  |  |  |  | 1.71 | 0.06 | 0.53 | 0.3 |
| Malisor | 66 | 69.8 | 20.9 | 22.8 | 0.79 | 182.62 | 769.19 | 421.5 | 0.88 | 48.7 | 54.06 | 28.5 | 0.88 | 34.85 | 8.45 | 16.75 | 0.67 | 15.3 | 2.2 | 19.8 | 0.18 | 1.4 | 0.12 | 0.53 | 0.47 |
| BimbG | 73 | 74.9 | 36.3 | 10.5 | 0.93 | 218.56 | 546.97 | 414 | 0.84 | 52.3 | 13.23 | 57.8 | 0.48 | 39.82 | 5.1 | 19.71 | 0.51 | 16.4 | 0 | 38.1 | 0 | 1.43 | 0.33 | 0.57 | 0.7 |
| CSM417 | 23 | 66.5 | 0.55 | 6.5 | 0.25 | 223.3 | 407 | 376.8 | 0.81 | 59.6 | 31.63 | 65.2 | 0.66 | 40.91 | 13.2 | 12.68 | 0.81 |  |  |  |  | 2.06 | 0.25 | 0.37 | 0.73 |
| average | 57.3 | 69.5 |  |  | 0.78 | 207.43 |  |  | 0.84 | 52.8 |  |  | 0.7 | 37.53 |  |  | 0.66 | 15.8 |  |  | 0.09 | 1.61 |  |  | 0.55 |

***Lata3 population characterized in 2013***
|  |  | Flag leaf [d] |  |  |  | Plant height [cm] |  |  |  | Peduncle length [cm] |  |  |  | Panicle length [cm] |  |  |  | Grain weight [g] |  |  |  | Grain yield [ton/ha] |  |  |  |
| --- | --- | --- | --- | --- | --- | --- | --- | --- | --- | --- | --- | --- | --- | --- | --- | --- | --- | --- | --- | --- | --- | --- | --- | --- | --- |
| ***Lata*** | N | Av | $\sigma_g^2$ | $\sigma_{ge}^2$ | $h^2$ | Av | $\sigma_g^2$ | $\sigma_{ge}^2$ | $h^2$ | Av | $\sigma_g^2$ | $\sigma_{ge}^2$ | $h^2$ | Av | $\sigma_g^2$ | $\sigma_{ge}^2$ | $h^2$ | Av | $\sigma_g^2$ | $\sigma_{ge}^2$ | $h^2$ | Av | $\sigma_g^2$ | $\sigma_{ge}^2$ | $h^2$ |
| Grinkan | 95 | 81.2 | 24.7 | 0.3 | 0.97 | 238.2 | 1596.8 | 105.8 | 0.93 | 42.8 | 27.2 | 1.4 | 0.88 | 29.2 | 6.1 | 1.3 | 0.77 | 2.1 | 0.04 | 0.01 | 0.86 | 2.47 | 0.31 | 0.05 | 0.8 |
| IS23645 | 54 | 83.2 | 40.3 | 0.9 | 0.97 | 250.6 | 3387.6 | 1.9 | 0.97 | 41.0 | 49.1 | 2.1 | 0.89 | 30.8 | 9.8 | 0.0 | 0.84 | 1.86 | 0.08 | 0.01 | 0.92 | 1.46 | 0.22 | 0.09 | 0.78 |
| SK5912 | 76 | 86.0 | 34.4 | 1.4 | 0.97 | 248.9 | 1170.1 | 74.6 | 0.92 | 43.4 | 27.6 | 0.3 | 0.9 | 28.1 | 6.1 | 1.7 | 0.73 | 2.33 | 0.09 | 0.01 | 0.91 | 2.07 | 0.21 | 0.08 | 0.77 |
| DouaG | 72 | 83.7 | 24.9 | 0.9 | 0.97 | 276.9 | 558.5 | 0.0 | 0.87 | 48.6 | 23.2 | 0.3 | 0.87 | 30.5 | 4.4 | 0.0 | 0.75 | 2.13 | 0.05 | 0 | 0.95 | 2.32 | 0.09 | 0.07 | 0.56 |
| Framida | 60 | 81.6 | 56.6 | 1.3 | 0.98 | 245.2 | 369.7 | 0.0 | 0.78 | 42.2 | 33.1 | 0.8 | 0.92 | 28.0 | 6.4 | 1.0 | 0.76 | 2.28 | 0.07 | 0.01 | 0.91 | 2.01 | 0.14 | 0.16 | 0.56 |
| Gnossiconi | 67 | 79.9 | 18.1 | 0.6 | 0.96 | 269.5 | 269.8 | 64.0 | 0.74 | 46.5 | 14.1 | 0.0 | 0.77 | 29.2 | 3.6 | 0.0 | 0.68 | 2.24 | 0.04 | 0 | 0.89 | 2.37 | 0.05 | 0.05 | 0.4 |
| IS15401 | 89 | 85.2 | 49.1 | 1.6 | 0.97 | 274.7 | 799.7 | 7.5 | 0.91 | 43.8 | 24.3 | 4.3 | 0.83 | 28.5 | 5.9 | 0.0 | 0.87 | 2.36 | 0.06 | 0.01 | 0.9 | 2.45 | 0.19 | 0.13 | 0.67 |
| IS23540 | 68 | 82.0 | 52.9 | 3.6 | 0.96 | 237.7 | 1011.8 | 20.2 | 0.93 | 39.8 | 30.0 | 2.9 | 0.88 | 26.0 | 4.6 | 1.6 | 0.69 | 2.24 | 0.08 | 0.01 | 0.91 | 2.18 | 0.11 | 0.06 | 0.6 |
| Fara | 82 | 90.0 | 42.9 | 0.9 | 0.97 | 295.2 | 924.6 | 70.8 | 0.86 | 47.2 | 22.9 | 0.0 | 0.86 | 32.2 | 7.6 | 0.5 | 0.78 | 2.25 | 0.07 | 0.01 | 0.91 | 2.07 | 0.18 | 0.05 | 0.74 |
| Ngolofing | 76 | 82.0 | 2.1 | 0.0 | 0.8 | 275.8 | 331.2 | 0.0 | 0.82 | 44.9 | 12.9 | 0.0 | 0.81 | 30.8 | 3.1 | 0.0 | 0.66 | 2.2 | 0.02 | 0.01 | 0.76 | 2.28 | 0.03 | 0.09 | 0.31 |
| Sambalma | 88 | 86.3 | 31.2 | 0.0 | 0.97 | 265.9 | 628.5 | 32.5 | 0.88 | 43.0 | 18.0 | 1.7 | 0.84 | 32.7 | 7.3 | 0.9 | 0.78 | 2.27 | 0.04 | 0.01 | 0.84 | 2.46 | 0.14 | 0.07 | 0.65 |
| SC566-14 | 69 | 82.5 | 74.7 | 5.8 | 0.96 | 253.5 | 1465.7 | 124.1 | 0.92 | 43.0 | 28.3 | 0.3 | 0.89 | 28.1 | 10.9 | 0.1 | 0.88 | 2.22 | 0.04 | 0.01 | 0.88 | 2.21 | 0.28 | 0.07 | 0.78 |
| average | 74 | 83.6 | 37.7 | 1.43 | 0.95 | 261 | 1042.8 | 41.8 | 0.88 | 43.9 | 25.9 | 1.2 | 0.9 | 29.5 | 6.3 | 0.6 | 0.77 | 2.21 | 0.06 | 0.01 | 0.89 | 2.2 | 0.16 | 0.08 | 0.64 |

**Tables S4:** Lists of the five most influential environmental covariables on each trait Table of most influential EC on traits with recurrent parent, year of phenotyping, trait, most influential EC in order of influence, average R2 trait-EC of the different tested window, R2 trait-EC of the best window, starting and end day of the best sowing window, value of the EC in the environment during the best window

| RP | year | trait | EC | R2_av | R2_win | start | end | SB1 | SB2 | CZ1 | CZ2 |
| --- | --- | --- | --- | --- | --- | --- | --- | --- | --- | --- | --- |
| Grinkan | 2012 | FLAG | photoperiod | 0.997 | 1 | 28 | 67 | 12.4 | 12.1 | 12.4 | 12.2 |
| Grinkan | 2012 | FLAG | solarRad | 0.844 | 0.999 | 9 | 88 | 3003.5 | 2928 | 3002.5 | 2937.1 |
| Grinkan | 2012 | FLAG | Trange | 0.82 | 1 | 3 | 82 | 8.8 | 9.4 | 8.9 | 9.4 |
| Grinkan | 2012 | FLAG | hSun | 0.629 | 0.997 | 1 | 80 | 512.1 | 542.9 | 511.3 | 538 |
| Grinkan | 2012 | FLAG | rain | 0.378 | 0.959 | 17 | 76 | 573.6 | 417.4 | 540.4 | 408.7 |
| Grinkan | 2012 | PH | photoperiod | 0.775 | 0.895 | 1 | 20 | 12.7 | 12.5 | 12.7 | 12.6 |
| Grinkan | 2012 | PH | solarRad | 0.746 | 0.995 | 23 | 42 | 756.5 | 752.6 | 758.9 | 754.5 |
| Grinkan | 2012 | PH | Trange | 0.628 | 0.985 | 6 | 65 | 8.8 | 8.9 | 8.7 | 8.8 |
| Grinkan | 2012 | PH | hSun | 0.621 | 0.999 | 16 | 75 | 389.4 | 407.2 | 376.7 | 398.8 |
| Grinkan | 2012 | PH | hum | 0.523 | 0.999 | 14 | 113 | 77.2 | 74 | 79.8 | 75.4 |
| Grinkan | 2012 | N_N | photoperiod | 0.819 | 0.915 | 1 | 20 | 12.7 | 12.5 | 12.7 | 12.6 |
| Grinkan | 2012 | N_N | solarRad | 0.758 | 0.901 | 1 | 60 | 2266.3 | 2248.7 | 2272.4 | 2255 |
| Grinkan | 2012 | N_N | Trange | 0.633 | 0.974 | 46 | 65 | 8.8 | 9.8 | 9 | 9.3 |
| Grinkan | 2012 | N_N | hSun | 0.62 | 0.994 | 7 | 66 | 376.3 | 396.7 | 373.6 | 383.1 |
| Grinkan | 2012 | N_N | hum | 0.388 | 0.841 | 19 | 118 | 76.9 | 73.1 | 78.8 | 74.2 |
| Grinkan | 2012 | N_L | FRUE | 0.32 | 0.956 | 34 | 113 | 1 | 1 | 1 | 1 |
| Grinkan | 2012 | N_L | rain | 0.291 | 0.967 | 9 | 48 | 359.8 | 419.7 | 436.6 | 396.7 |
| Grinkan | 2012 | N_L | PETP | 0.275 | 0.963 | 5 | 24 | 4.7 | -0.5 | -1.8 | 1.4 |
| Grinkan | 2012 | N_L | ETP | 0.168 | 0.644 | 74 | 93 | 9.1 | 9.9 | 10.3 | 10.1 |
| Grinkan | 2012 | N_L | hSun | 0.145 | 0.919 | 22 | 41 | 137.2 | 120.2 | 114.8 | 124.6 |
| Grinkan | 2012 | PAN | PETP | 0.356 | 0.999 | 8 | 67 | -0.4 | 0.2 | 0.5 | -1.9 |
| Grinkan | 2012 | PAN | rain | 0.345 | 0.995 | 7 | 66 | 516.6 | 539.1 | 554.9 | 449.7 |
| Grinkan | 2012 | PAN | VPD | 0.322 | 0.634 | 52 | 71 | 0.8 | 1 | 0.9 | 1.3 |
| Grinkan | 2012 | PAN | Tmin | 0.295 | 0.549 | 68 | 87 | 31.3 | 33.2 | 32.4 | 35.8 |
| Grinkan | 2012 | PAN | hum | 0.277 | 0.924 | 7 | 26 | 78.6 | 78.6 | 77.9 | 81.9 |
| Grinkan | 2012 | PED | DD | 0.857 | 0.979 | 88 | 107 | 377 | 396.8 | 433.4 | 452.4 |
| Grinkan | 2012 | PED | SVP | 0.85 | 0.974 | 88 | 107 | 0.2 | 0.2 | 0.2 | 0.3 |
| Grinkan | 2012 | PED | Tmin | 0.809 | 0.993 | 19 | 78 | 30.6 | 31.3 | 31.9 | 32.4 |
| Grinkan | 2012 | PED | ETP | 0.788 | 1 | 53 | 112 | 9.1 | 9.5 | 9.9 | 10.1 |
| Grinkan | 2012 | PED | FRUE | 0.772 | 0.997 | 4 | 63 | 1 | 1 | 1 | 1 |
| Grinkan | 2012 | YIELD | Trange | 0.382 | 0.995 | 20 | 39 | 8.6 | 8.6 | 8.3 | 8.9 |
| Grinkan | 2012 | YIELD | hSun | 0.352 | 0.976 | 2 | 61 | 376.3 | 377 | 371.1 | 381.7 |
| Grinkan | 2012 | YIELD | hum | 0.346 | 0.869 | 69 | 88 | 76.4 | 74.5 | 83.2 | 72.4 |
| Grinkan | 2012 | YIELD | rain | 0.33 | 0.998 | 5 | 64 | 509.8 | 524.2 | 584.4 | 460.2 |
| Grinkan | 2012 | YIELD | photoperiod | 0.32 | 0.321 | 2 | 101 | 12.4 | 12.1 | 12.3 | 12.1 |
| Grinkan | 2013 | FLAG | photoperiod | 0.92 | 0.926 | 15 | 34 | 12.6 | 12.4 | 12.5 | 12.3 |
| Grinkan | 2013 | FLAG | solarRad | 0.693 | 0.866 | 64 | 83 | 741.5 | 710.8 | 708.1 | 674.2 |
| Grinkan | 2013 | FLAG | hSun | 0.623 | 0.991 | 61 | 80 | 145 | 175.7 | 163.2 | 181.5 |
| Grinkan | 2013 | FLAG | hum | 0.584 | 0.993 | 9 | 28 | 75.9 | 80.7 | 78.3 | 81.8 |
| Grinkan | 2013 | FLAG | Trange | 0.514 | 0.954 | 58 | 77 | 8.2 | 10.4 | 9.7 | 12 |
| Grinkan | 2013 | PH | solarRad | 0.887 | 0.961 | 41 | 60 | 755.5 | 742.5 | 741.7 | 718.6 |
| Grinkan | 2013 | PH | photoperiod | 0.874 | 0.884 | 5 | 24 | 12.7 | 12.5 | 12.6 | 12.4 |
| Grinkan | 2013 | PH | Trange | 0.848 | 1 | 16 | 75 | 8.3 | 8.7 | 9.1 | 10.2 |
| Grinkan | 2013 | PH | VPD | 0.822 | 0.998 | 66 | 85 | 1 | 1.3 | 1.7 | 2.6 |
| Grinkan | 2013 | PH | hum | 0.814 | 0.998 | 45 | 64 | 80.7 | 79.1 | 77.9 | 74.6 |
| Grinkan | 2013 | N_N | photoperiod | 0.774 | 0.812 | 1 | 20 | 12.7 | 12.6 | 12.6 | 12.4 |
| Grinkan | 2013 | N_N | solarRad | 0.65 | 0.726 | 85 | 104 | 712.6 | 670 | 665.8 | 630.8 |
| Grinkan | 2013 | N_N | hSun | 0.565 | 0.995 | 2 | 21 | 155.6 | 130.1 | 147.8 | 128 |
| Grinkan | 2013 | N_N | Trange | 0.523 | 0.873 | 58 | 77 | 8.2 | 10.4 | 9.7 | 12 |
| Grinkan | 2013 | N_N | hum | 0.452 | 0.998 | 3 | 42 | 77.1 | 80 | 78.3 | 80.6 |
| Grinkan | 2013 | N_L | VPD | 0.781 | 0.976 | 4 | 63 | 1 | 1 | 1.3 | 1.3 |
| Grinkan | 2013 | N_L | SVP | 0.772 | 0.998 | 12 | 51 | 0.2 | 0.2 | 0.2 | 0.2 |
| Grinkan | 2013 | N_L | DD | 0.772 | 0.998 | 13 | 52 | 764.4 | 742.9 | 786.3 | 807.1 |
| Grinkan | 2013 | N_L | Tmin | 0.754 | 0.998 | 13 | 52 | 31.2 | 30.6 | 32.2 | 32.8 |
| Grinkan | 2013 | N_L | photothermal | 0.728 | 0.997 | 25 | 64 | 9380.2 | 9144.8 | 9741.9 | 9961.7 |
| Grinkan | 2013 | PAN | SVP | 0.814 | 0.994 | 11 | 50 | 0.2 | 0.2 | 0.2 | 0.2 |
| Grinkan | 2013 | PAN | DD | 0.813 | 0.991 | 11 | 50 | 762.5 | 740.9 | 791 | 804 |
| Grinkan | 2013 | PAN | VPD | 0.795 | 0.995 | 5 | 64 | 1 | 0.9 | 1.3 | 1.3 |
| Grinkan | 2013 | PAN | Tmin | 0.79 | 0.998 | 13 | 52 | 31.2 | 30.6 | 32.2 | 32.8 |
| Grinkan | 2013 | PAN | photothermal | 0.765 | 0.998 | 25 | 64 | 9380.2 | 9144.8 | 9741.9 | 9961.7 |
| Grinkan | 2013 | PED | SVP | 0.804 | 0.993 | 12 | 51 | 0.2 | 0.2 | 0.2 | 0.2 |
| Grinkan | 2013 | PED | DD | 0.803 | 0.991 | 20 | 59 | 755.8 | 739.9 | 784.7 | 812.5 |
| Grinkan | 2013 | PED | VPD | 0.801 | 0.989 | 5 | 64 | 1 | 0.9 | 1.3 | 1.3 |
| Grinkan | 2013 | PED | Tmin | 0.788 | 1 | 13 | 52 | 31.2 | 30.6 | 32.2 | 32.8 |
| Grinkan | 2013 | PED | photothermal | 0.75 | 0.998 | 25 | 64 | 9380.2 | 9144.8 | 9741.9 | 9961.7 |
| Grinkan | 2013 | YIELD | hum | 0.746 | 1 | 10 | 69 | 79.2 | 79.6 | 78.9 | 76.9 |
| Grinkan | 2013 | YIELD | VPD | 0.736 | 0.991 | 33 | 52 | 0.9 | 0.9 | 1.1 | 1.5 |
| Grinkan | 2013 | YIELD | solarRad | 0.693 | 0.893 | 21 | 40 | 756.5 | 755 | 755.8 | 744.7 |
| Grinkan | 2013 | YIELD | Trange | 0.651 | 0.986 | 24 | 43 | 8.3 | 7.8 | 8.5 | 9.7 |
| Grinkan | 2013 | YIELD | Tmin | 0.642 | 0.934 | 26 | 45 | 31.1 | 30.4 | 31.8 | 33.1 |
| Kenin-K | 2012 | FLAG | photoperiod | 0.995 | 0.999 | 43 | 62 | 12.4 | 12.1 | 12.4 | 12.1 |
| Kenin-K | 2012 | FLAG | solarRad | 0.817 | 0.997 | 49 | 68 | 753.5 | 728.2 | 752.4 | 730.7 |
| Kenin-K | 2012 | FLAG | Trange | 0.8 | 1 | 3 | 82 | 8.8 | 9.4 | 8.9 | 9.4 |
| Kenin-K | 2012 | FLAG | hSun | 0.58 | 0.994 | 1 | 80 | 512.1 | 542.9 | 511.3 | 538 |
| Kenin-K | 2012 | FLAG | photothermal | 0.553 | 0.997 | 12 | 31 | 2600.1 | 2224.2 | 2598.4 | 2282.5 |
| Kenin-K | 2012 | PH | hum | 0.551 | 0.998 | 48 | 87 | 77.2 | 75.8 | 83.9 | 77.8 |
| Kenin-K | 2012 | PH | Tmax | 0.504 | 0.996 | 19 | 38 | 22.2 | 21.9 | 23.9 | 22.5 |
| Kenin-K | 2012 | PH | solarRad | 0.414 | 0.959 | 18 | 37 | 756 | 754.6 | 758.7 | 756.5 |
| Kenin-K | 2012 | PH | photoperiod | 0.371 | 0.523 | 1 | 20 | 12.7 | 12.5 | 12.7 | 12.6 |
| Kenin-K | 2012 | PH | hSun | 0.365 | 0.985 | 12 | 71 | 393.5 | 396.2 | 375.2 | 392.6 |
| Kenin-K | 2012 | N_N | photoperiod | 0.684 | 0.836 | 1 | 20 | 12.7 | 12.5 | 12.7 | 12.6 |
| Kenin-K | 2012 | N_N | solarRad | 0.668 | 0.994 | 9 | 48 | 1511.7 | 1505.7 | 1516.7 | 1509.6 |
| Kenin-K | 2012 | N_N | hSun | 0.54 | 0.994 | 15 | 74 | 389.3 | 405.2 | 380.1 | 395.2 |
| Kenin-K | 2012 | N_N | hum | 0.52 | 0.991 | 7 | 106 | 77.7 | 74.8 | 80.4 | 76.8 |
| Kenin-K | 2012 | N_N | Trange | 0.517 | 0.97 | 6 | 65 | 8.8 | 8.9 | 8.7 | 8.8 |
| Kenin-K | 2012 | N_L | FRUE | 0.593 | 0.972 | 3 | 22 | 1 | 1 | 1 | 1 |
| Kenin-K | 2012 | N_L | rain | 0.439 | 0.999 | 15 | 54 | 277 | 404.2 | 409.4 | 377.2 |
| Kenin-K | 2012 | N_L | PETP | 0.438 | 0.976 | 18 | 57 | -2.1 | 1 | 0.7 | 0.3 |
| Kenin-K | 2012 | N_L | ETP | 0.397 | 0.895 | 75 | 94 | 9.2 | 9.9 | 10.2 | 10 |
| Kenin-K | 2012 | N_L | Tmin | 0.344 | 0.55 | 47 | 66 | 30.5 | 31.8 | 31.6 | 32.5 |
| Kenin-K | 2012 | PAN | photoperiod | 0.937 | 0.941 | 33 | 52 | 12.5 | 12.2 | 12.5 | 12.2 |
| Kenin-K | 2012 | PAN | solarRad | 0.843 | 0.932 | 56 | 75 | 749.7 | 717.6 | 747.4 | 720.2 |
| Kenin-K | 2012 | PAN | Trange | 0.802 | 0.999 | 56 | 75 | 8.8 | 10.2 | 9.2 | 9.8 |
| Kenin-K | 2012 | PAN | hSun | 0.714 | 0.995 | 53 | 72 | 131.9 | 147.5 | 133.9 | 142.4 |
| Kenin-K | 2012 | PAN | photothermal | 0.544 | 0.986 | 17 | 36 | 2676.2 | 2350.1 | 2577.4 | 2425.7 |
| Kenin-K | 2012 | PED | rain | 0.268 | 1 | 34 | 53 | 162.9 | 255.4 | 262.2 | 131 |
| Kenin-K | 2012 | PED | PETP | 0.261 | 0.999 | 37 | 56 | -2 | 3.1 | 3.9 | -3.5 |
| Kenin-K | 2012 | PED | hum | 0.131 | 0.601 | 7 | 26 | 78.6 | 78.6 | 77.9 | 81.9 |
| Kenin-K | 2012 | PED | photothermal | 0.127 | 0.982 | 27 | 46 | 2525.5 | 2194.5 | 2236.6 | 2627.2 |
| Kenin-K | 2012 | PED | hSun | 0.113 | 0.989 | 23 | 62 | 258.3 | 251.5 | 249.5 | 261.6 |
| Kenin-K | 2012 | YIELD | FRUE | 0.385 | 0.964 | 14 | 113 | 1 | 1 | 1 | 1 |
| Kenin-K | 2012 | YIELD | Tmax | 0.305 | 0.736 | 12 | 31 | 21.8 | 22 | 23.9 | 22.9 |
| Kenin-K | 2012 | YIELD | ETP | 0.288 | 0.886 | 7 | 26 | 8.6 | 8.9 | 10 | 8.8 |
| Kenin-K | 2012 | YIELD | hum | 0.267 | 0.774 | 59 | 78 | 78 | 78 | 84.6 | 79 |
| Kenin-K | 2012 | YIELD | DD | 0.254 | 0.724 | 7 | 26 | 358.5 | 364 | 403.8 | 384.4 |
| Kenin-K | 2013 | FLAG | photoperiod | 0.975 | 0.978 | 4 | 43 | 12.6 | 12.4 | 12.5 | 12.3 |
| Kenin-K | 2013 | FLAG | solarRad | 0.842 | 0.999 | 55 | 74 | 749.1 | 725.3 | 723.4 | 692.8 |
| Kenin-K | 2013 | FLAG | hSun | 0.69 | 0.996 | 14 | 73 | 416.7 | 442.1 | 446.7 | 487.8 |
| Kenin-K | 2013 | FLAG | Trange | 0.686 | 0.998 | 36 | 75 | 8.1 | 9.2 | 9.4 | 10.7 |
| Kenin-K | 2013 | FLAG | hum | 0.645 | 0.999 | 49 | 68 | 80.7 | 77.7 | 77 | 72.8 |
| Kenin-K | 2013 | PH | photoperiod | 0.966 | 0.972 | 23 | 42 | 12.6 | 12.3 | 12.4 | 12.2 |
| Kenin-K | 2013 | PH | solarRad | 0.83 | 0.945 | 88 | 107 | 707.3 | 663.8 | 659.4 | 625.2 |
| Kenin-K | 2013 | PH | Trange | 0.714 | 0.976 | 58 | 77 | 8.2 | 10.4 | 9.7 | 12 |
| Kenin-K | 2013 | PH | hSun | 0.707 | 0.999 | 54 | 73 | 141.3 | 162.2 | 157.5 | 172.8 |
| Kenin-K | 2013 | PH | hum | 0.627 | 0.961 | 9 | 28 | 75.9 | 80.7 | 78.3 | 81.8 |
| Kenin-K | 2013 | N_N | photoperiod | 0.769 | 0.773 | 32 | 51 | 12.5 | 12.2 | 12.3 | 12 |
| Kenin-K | 2013 | N_N | solarRad | 0.584 | 0.737 | 90 | 109 | 703.7 | 659.7 | 655.2 | 621.6 |
| Kenin-K | 2013 | N_N | hSun | 0.581 | 1 | 67 | 86 | 159.1 | 176.1 | 167.9 | 175.9 |
| Kenin-K | 2013 | N_N | Trange | 0.487 | 0.808 | 58 | 77 | 8.2 | 10.4 | 9.7 | 12 |
| Kenin-K | 2013 | N_N | hum | 0.401 | 0.999 | 5 | 44 | 77.4 | 80.2 | 78.8 | 80.2 |
| Kenin-K | 2013 | N_L | photoperiod | 0.956 | 0.96 | 1 | 80 | 12.5 | 12.2 | 12.3 | 12.1 |
| Kenin-K | 2013 | N_L | solarRad | 0.911 | 0.991 | 55 | 74 | 749.1 | 725.3 | 723.4 | 692.8 |
| Kenin-K | 2013 | N_L | Trange | 0.852 | 1 | 25 | 104 | 8.8 | 10.1 | 10.6 | 12.4 |
| Kenin-K | 2013 | N_L | hum | 0.777 | 0.998 | 49 | 68 | 80.7 | 77.7 | 77 | 72.8 |
| Kenin-K | 2013 | N_L | Tmin | 0.776 | 0.99 | 62 | 81 | 31 | 33 | 34.3 | 36.7 |
| Kenin-K | 2013 | PAN | VPD | 0.85 | 1 | 18 | 77 | 1 | 1 | 1.3 | 1.6 |
| Kenin-K | 2013 | PAN | Tmin | 0.832 | 1 | 33 | 52 | 30.6 | 30.7 | 32 | 33.7 |
| Kenin-K | 2013 | PAN | Trange | 0.8 | 0.994 | 26 | 45 | 7.9 | 8 | 8.5 | 9.6 |
| Kenin-K | 2013 | PAN | solarRad | 0.782 | 0.865 | 34 | 53 | 756.7 | 748.4 | 748.2 | 729.3 |
| Kenin-K | 2013 | PAN | hum | 0.774 | 0.998 | 6 | 105 | 78.2 | 77.4 | 72.8 | 67.1 |
| Kenin-K | 2013 | PED | SVP | 0.714 | 0.99 | 12 | 51 | 0.2 | 0.2 | 0.2 | 0.2 |
| Kenin-K | 2013 | PED | photothermal | 0.713 | 1 | 21 | 60 | 9447.8 | 9132.6 | 9683.7 | 9869.8 |
| Kenin-K | 2013 | PED | DD | 0.713 | 0.992 | 12 | 51 | 763.3 | 741.5 | 788.6 | 804.8 |
| Kenin-K | 2013 | PED | VPD | 0.708 | 0.994 | 11 | 50 | 1.1 | 0.9 | 1.1 | 1.2 |
| Kenin-K | 2013 | PED | Tmin | 0.674 | 0.988 | 9 | 48 | 31.2 | 30.4 | 32.1 | 32.5 |
| Kenin-K | 2013 | YIELD | photoperiod | 0.714 | 0.807 | 1 | 20 | 12.7 | 12.6 | 12.6 | 12.4 |
| Kenin-K | 2013 | YIELD | solarRad | 0.664 | 0.727 | 1 | 60 | 2266.2 | 2254.1 | 2256.4 | 2220.1 |
| Kenin-K | 2013 | YIELD | hum | 0.502 | 0.996 | 5 | 24 | 75.4 | 80.3 | 75.7 | 82.1 |
| Kenin-K | 2013 | YIELD | hSun | 0.493 | 0.98 | 1 | 20 | 154.4 | 131.4 | 147.7 | 126.4 |
| Kenin-K | 2013 | YIELD | Trange | 0.491 | 0.867 | 57 | 76 | 8.4 | 10.2 | 9.6 | 12 |

**Tables S5:**
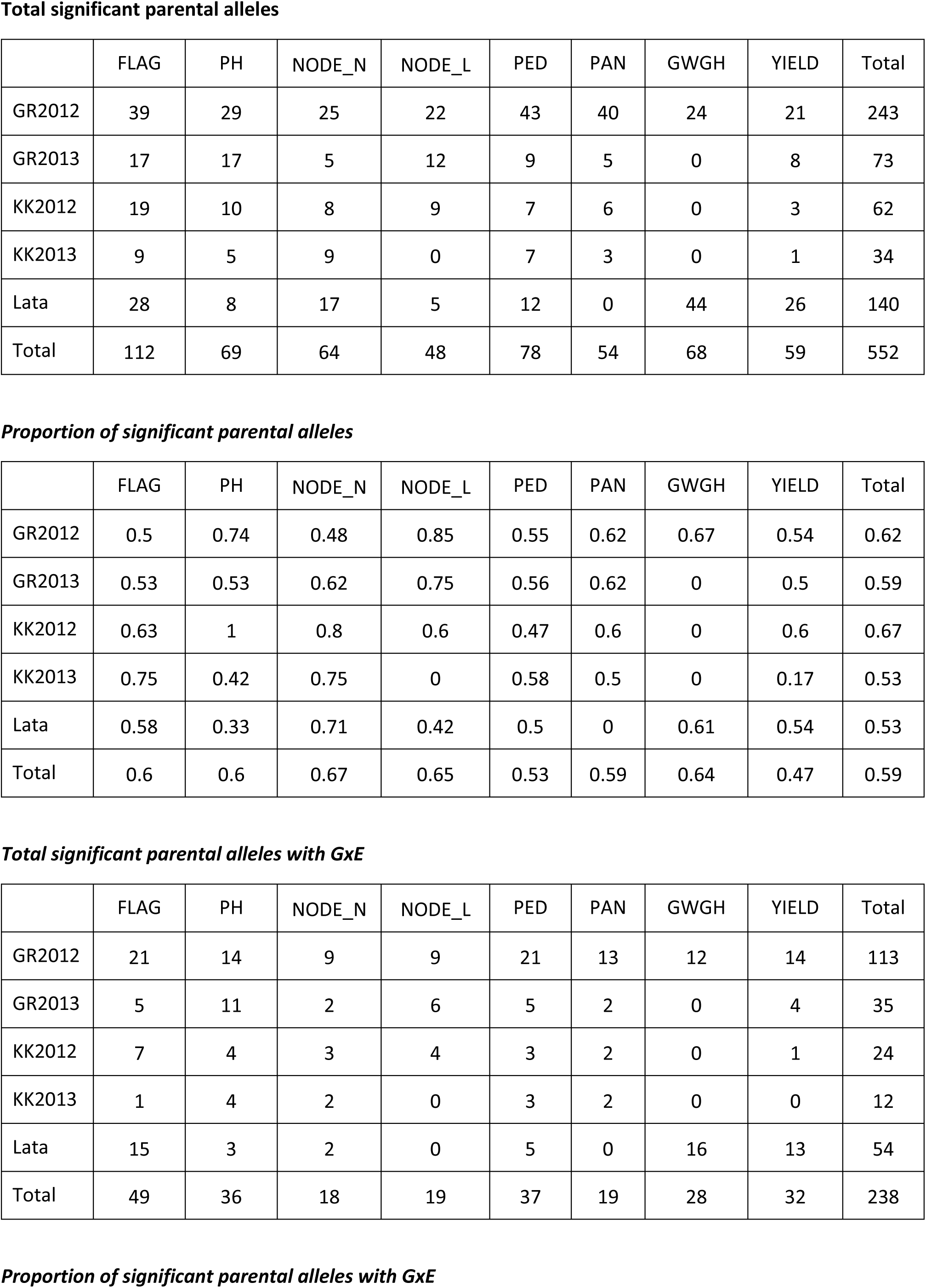
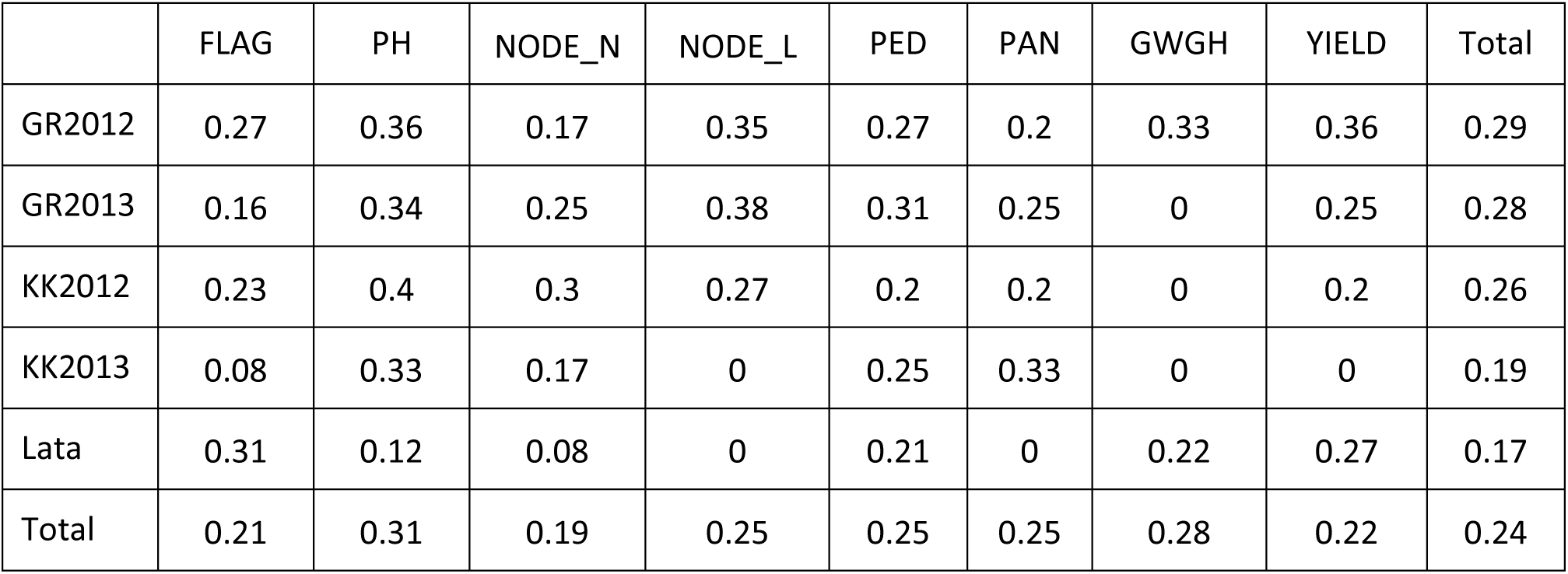
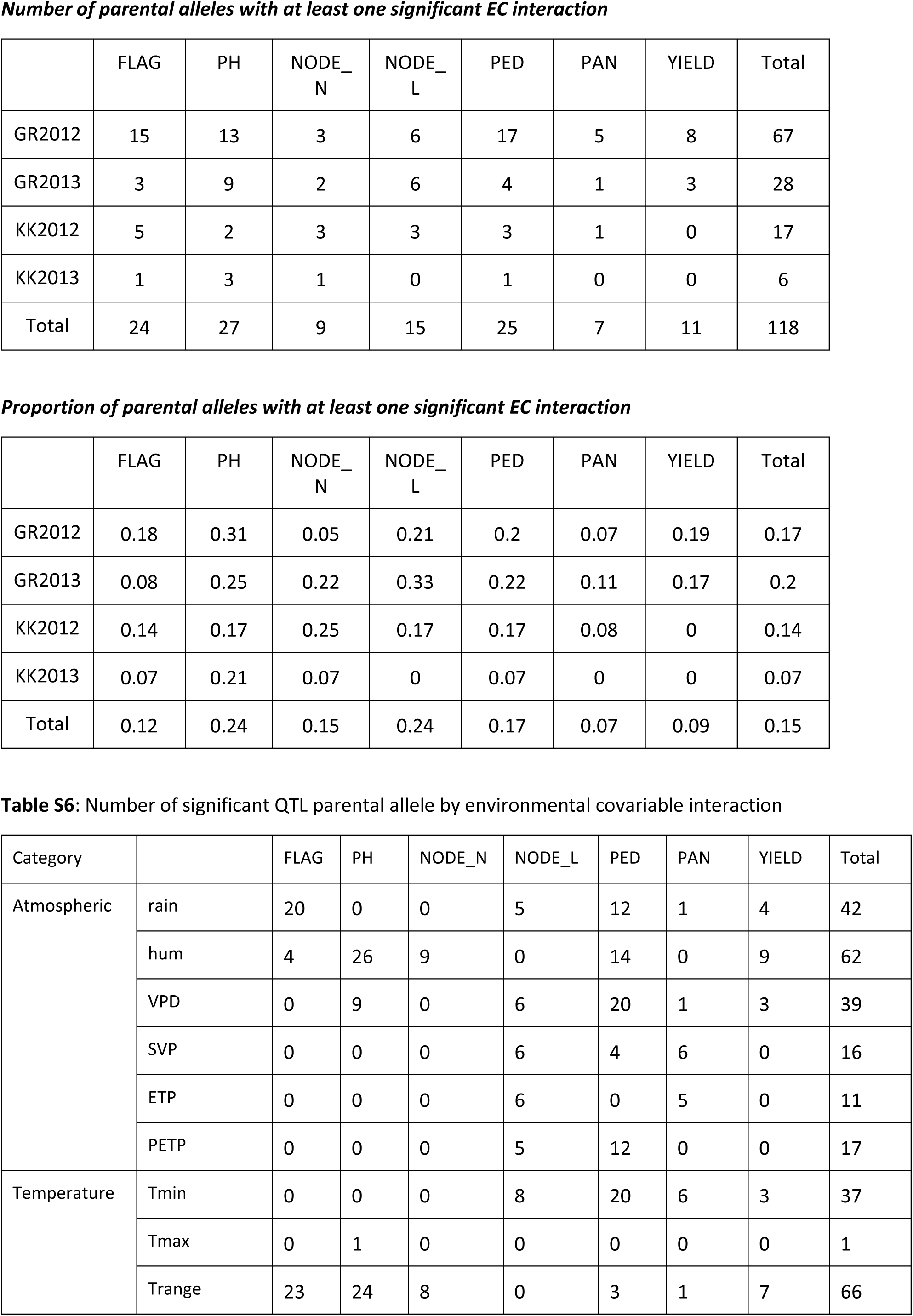
QTL parental allele effects detailed statistics (significant effect, QTLxE effect, QTLxEC effect)

**Table S6:**
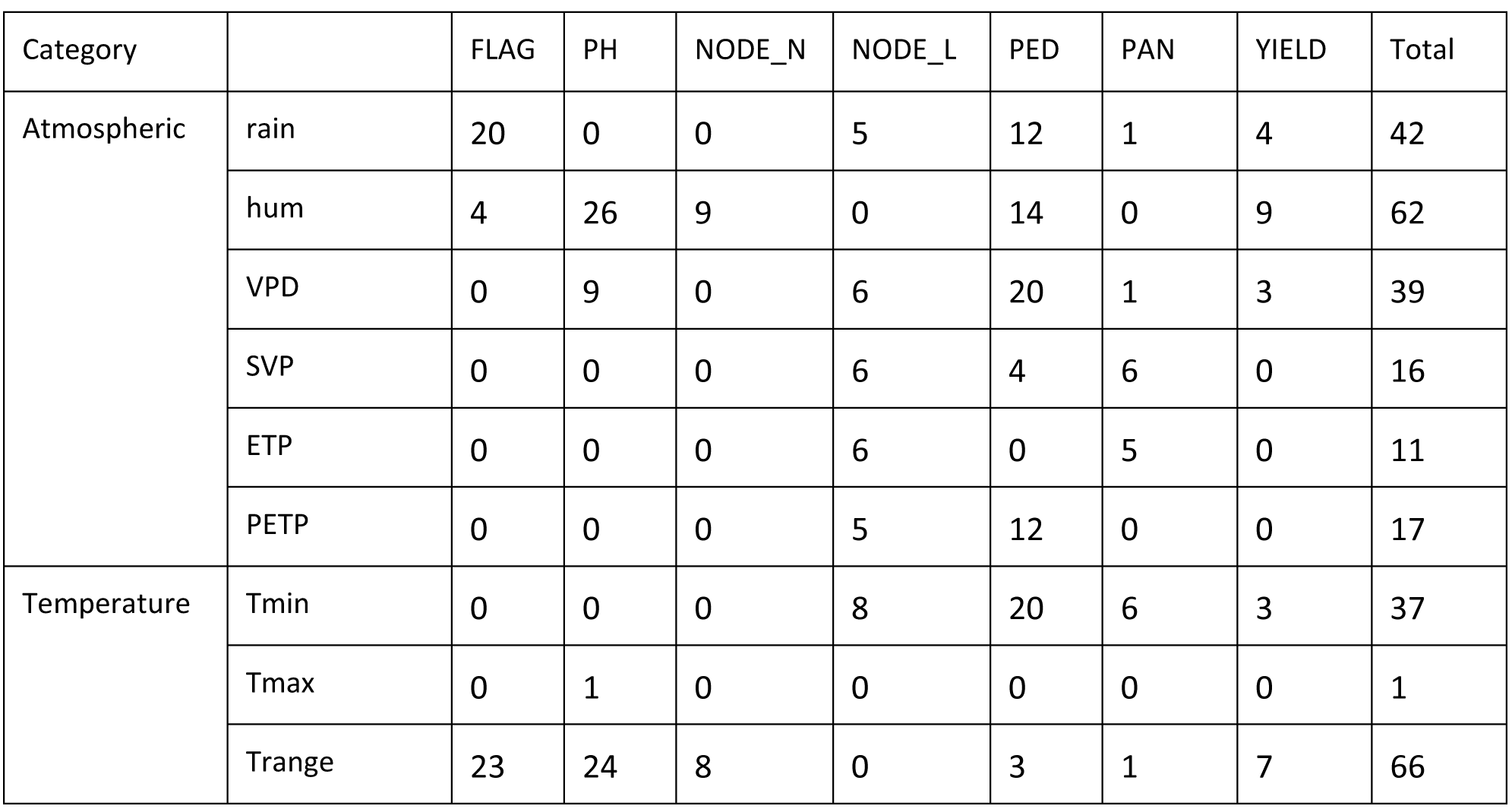
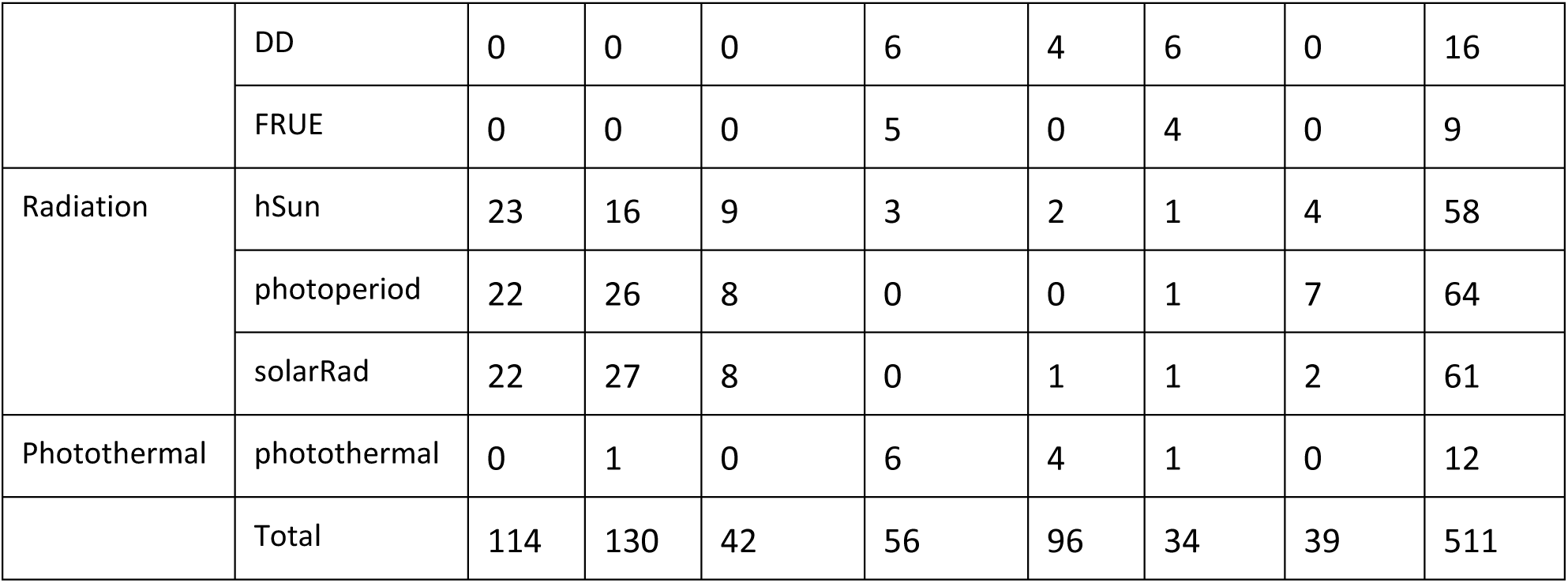
Number of significant QTL parental allele by environmental covariable interaction

**Table S7:** Significance of the QTL effect on yield after correction for a component trait (flag leaf appearance or plant height) We estimated the effect of QTL position on yield an the residual of yield (yield corrected) after correcting for the component trait by using a linear regression.

| Par | env | -log10(pval) |  |
| --- | --- | --- | --- |
|  |  | yield | yield corrected |
| IS15401 | SB1 | 0.37 | 0.01 |
| IS15401 | SB2 | 2.46 | 0.12 |
| IS15401 | CZ1 | 0.96 | 0.72 |
| IS15401 | CZ2 | 0.37 | 1.25 |

***GR2012 QTL chr 6 39 cM Grain yield flag leaf corrected***
| Par | env | log10(pval) |  |
| --- | --- | --- | --- |
|  |  | yield | yield (corrected) |
| B35 | SB1 | 0.08 | 0.9 |
| B35 | SB2 | 0.01 | 0.82 |
| B35 | CZ1 | 0.88 | 1.91 |
| B35 | CZ2 | 2.65 | 0.81 |
| SC566-14 | SB1 | 2.38 | 10.14 |
| SC566-14 | SB2 | 0.15 | 1.57 |
| SC566-14 | CZ1 | 5.48 | 9.21 |
| SC566-14 | CZ2 | 0.01 | 2.65 |

**Table S8:**
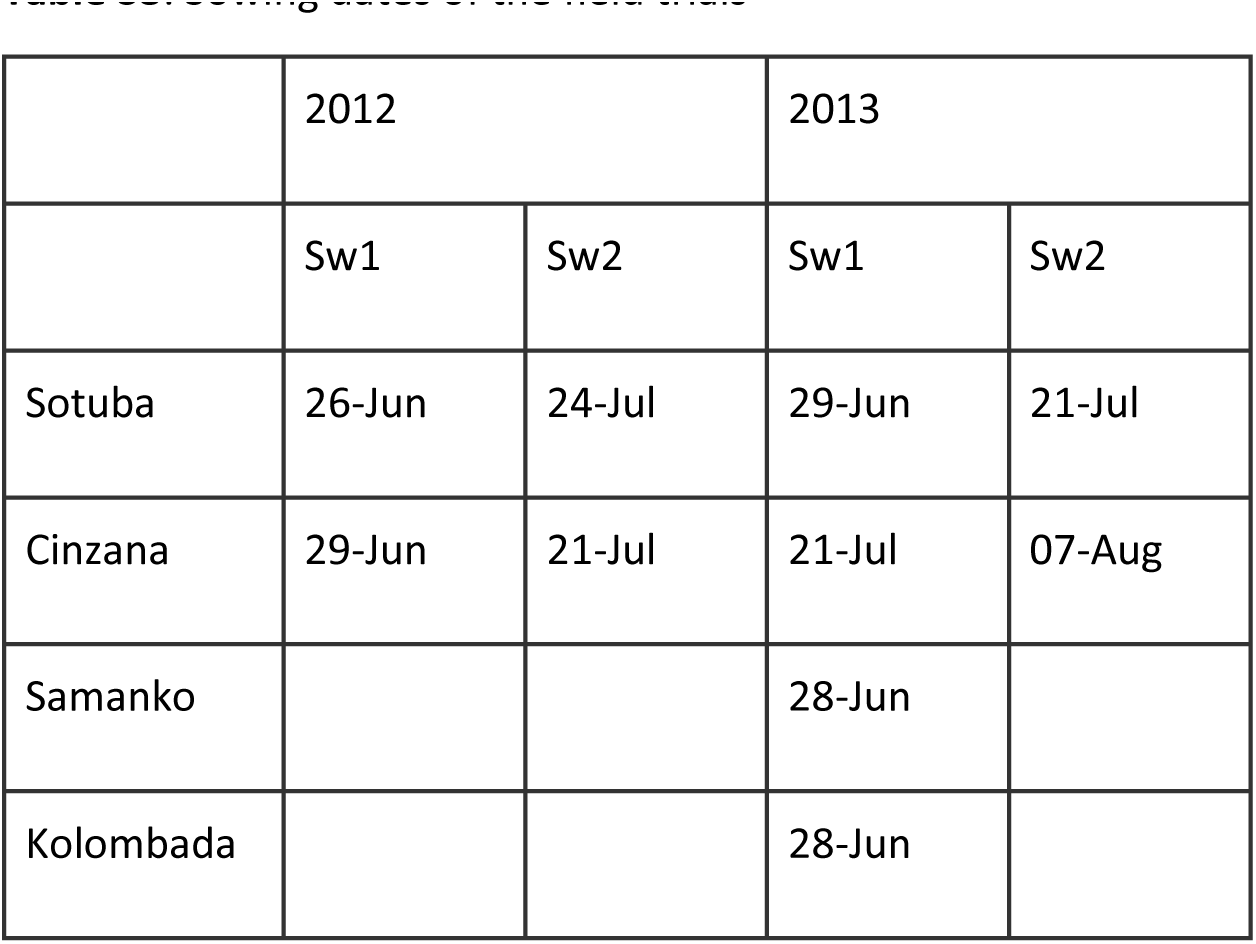
Sowing dates of the field trials

|  | 2012 |  | 2013 |  |
| --- | --- | --- | --- | --- |
|  | Sw1 | Sw2 | Sw1 | Sw2 |
| Sotuba | 26-Jun | 24-Jul | 29-Jun | 21-Jul |
| Cinzana | 29-Jun | 21-Jul | 21-Jul | 07-Aug |
| Samanko |  |  | 28-Jun |  |
| Kolombada |  |  | 28-Jun |  |

**Table S9:**
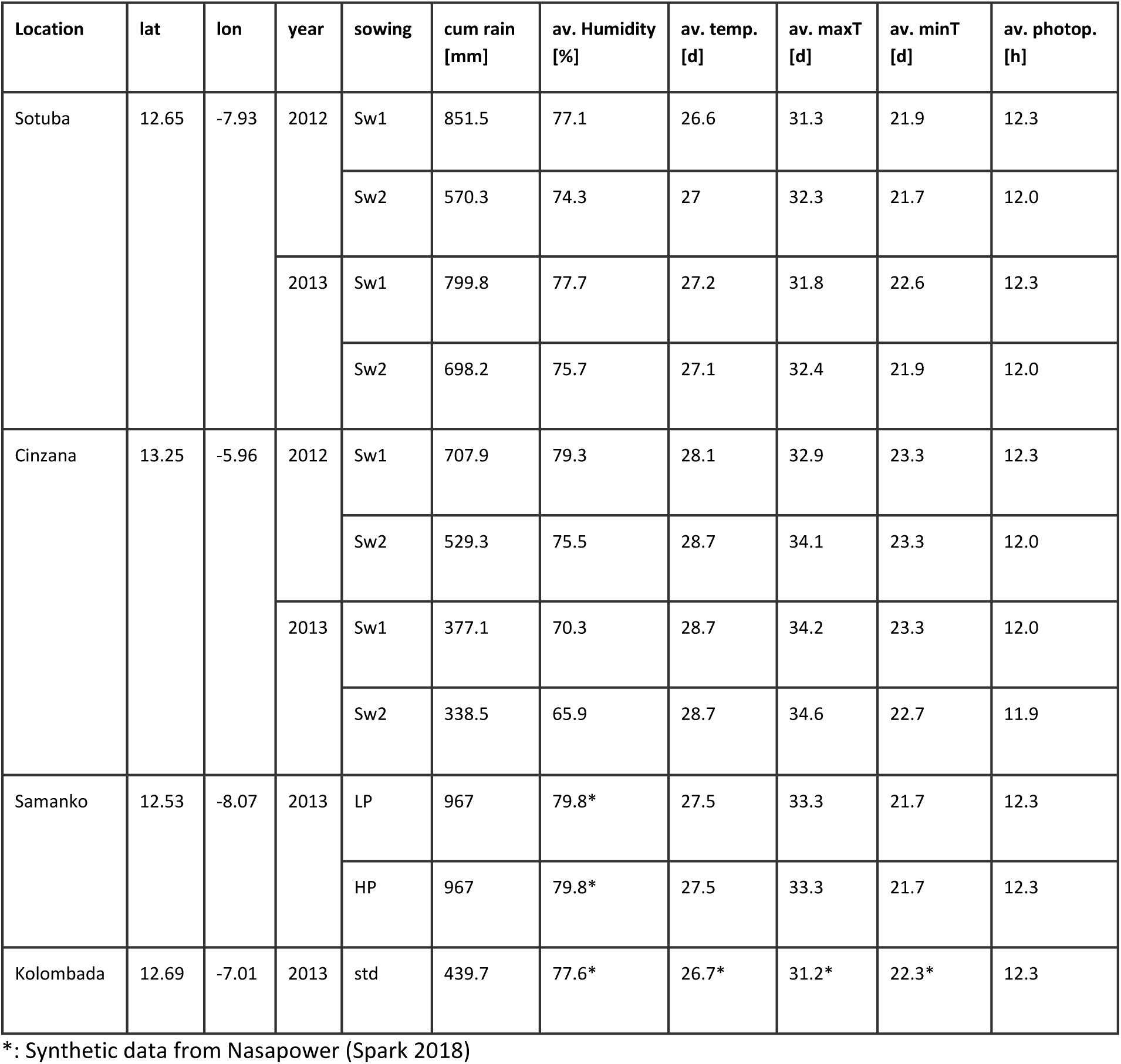
Environmental description of the multi-location trials

**Table S10:** Detail of the phenotyping per population, year environment, and trait

| Population | Year | Env | FLAG | PH | NODE_N | NODE_L | PED | PAN | GWGH | YIELD |
| --- | --- | --- | --- | --- | --- | --- | --- | --- | --- | --- |
| Grinkan | 2012 | Sotuba Sw1 | X | X | X | X | X | X | X | X |
|  |  | Sotuba Sw2 | X | X | X | X | X | X | X | X |
|  |  | Cinzana Sw1 | X | X | X | X | X | X |  | X |
|  |  | Cinzana Sw2 | X | X | X | X | X | X |  | X |
|  | 2013 | Sotuba Sw1 | X | X | X | X | X | X | X | X |
|  |  | Sotuba Sw2 | X | X | X | X | X | X | X | X |
|  |  | Cinzana Sw1 | X | X | X | X | X | X |  | X |
|  |  | Cinzana Sw2 | X | X | X | X | X | X |  | X |
| Kenin-Keni | 2012 | Sotuba Sw1 | X | X | X | X | X | X | X | X |
|  |  | Sotuba Sw2 | X | X | X | X | X | X | X | X |
|  |  | Cinzana Sw1 | X | X | X | X | X | X |  | X |
|  |  | Cinzana Sw2 | X | X | X | X | X | X |  | X |
|  | 2013 | Sotuba Sw1 | X | X | X | X | X | X | X | X |
|  |  | Sotuba Sw2 | X | X | X | X | X | X | X | X |
|  |  | Cinzana Sw1 | X | X | X | X | X | X |  | X |
|  |  | Cinzana Sw2 | X | X | X | X | X | X |  | X |
| Lata3 | 2013 | Samanko LP | X | X | X | X | X | X | X | X |
|  |  | Samanko LP | X | X | X | X | X | X | X | X |
|  |  | Kolombada | X | X | X | X |  |  | X | X |

**Table S11:** List of environmental covariables at the environment trials (Sotuba, Cinzana)

| Category | EC | Abbreviation | Unit | Observed/<br>inferred | Sum/mean |
| --- | --- | --- | --- | --- | --- |
| Atmospheric | cumulated rain | cum rain | mm | obs | sum |
|  | humidity | hum | % | obs | mean |
|  | vapour pressure deficit | VPD | kPa | inf | mean |
|  | slope of saturation VP curve | SVP | kPa/d | inf | mean |
|  | potential evapotranspiration | ETP | mm/day | inf | mean |
|  | water deficit | PETP | mm/day | inf | mean |
| Temperature | minimum temperature | Tmin | d | obs | mean |
|  | maximum temperature | Tmax | d | obs | mean |
|  | temperature range | Trange | d | obs | mean |
|  | cumulated degree day | DD | dd | obs | sum |
|  | T effect on radiation use efficiency | FRUE | 0-1 | inf | mean |
| Radiation | cumulated hour of sun | hsun | h | obs | sum |
|  | photoperiod | photo | h | inf | mean |
|  | solar radiation | SolRad | MJ/m<sup>2</sup>/day | inf | sum |
| Photothermal | Photothermal (photoperiod * DD) | photothermal | h*dd | obs | sum |

## Supporting experimental procedure

### Methods S1

Diversity tree construction methodology

This analysis was based on 137’003 SNPs common to the SAP (Boatwright et al. 2022) and SGT (https://www.globalsorghuminitiative.org/) panels. From a merge of these 2 studies, we excluded all wild accessions and kept SNP with less than 20% of missing data and with a Minor Allele Frequency (MAF) higher than 5%. To lighten the matrix without losing too much precision, we pruned with bcftools (Danecek et al, 2021) with these parameters : windows (-w) 1000 and r² (-m) bigger than 0.8. The software Darwin (https://darwin.cirad.fr/) were used to calculate the dissimilarity matrix and draw the tree (NJ method).

### Methods S2

Phenotype by environmental covariable analysis

To determine the ECs influence on the different traits, we applied the same method as Li et al. (2018), which consists of calculating the correlation between the trait mean across the environments and the EC values (Figure A) inside time windows of different size (20, 40, 60, 80, or 100) starting at different days of the plant cycle (Figure B). For each configuration of population (GR12, GR13, KK12, KK13) x traits we selected the five ECs with the highest average correlation and determined the most influential window and corresponding EC values as the one with the highest EC-trait correlation. Those values were later used in the QTLxEC models.

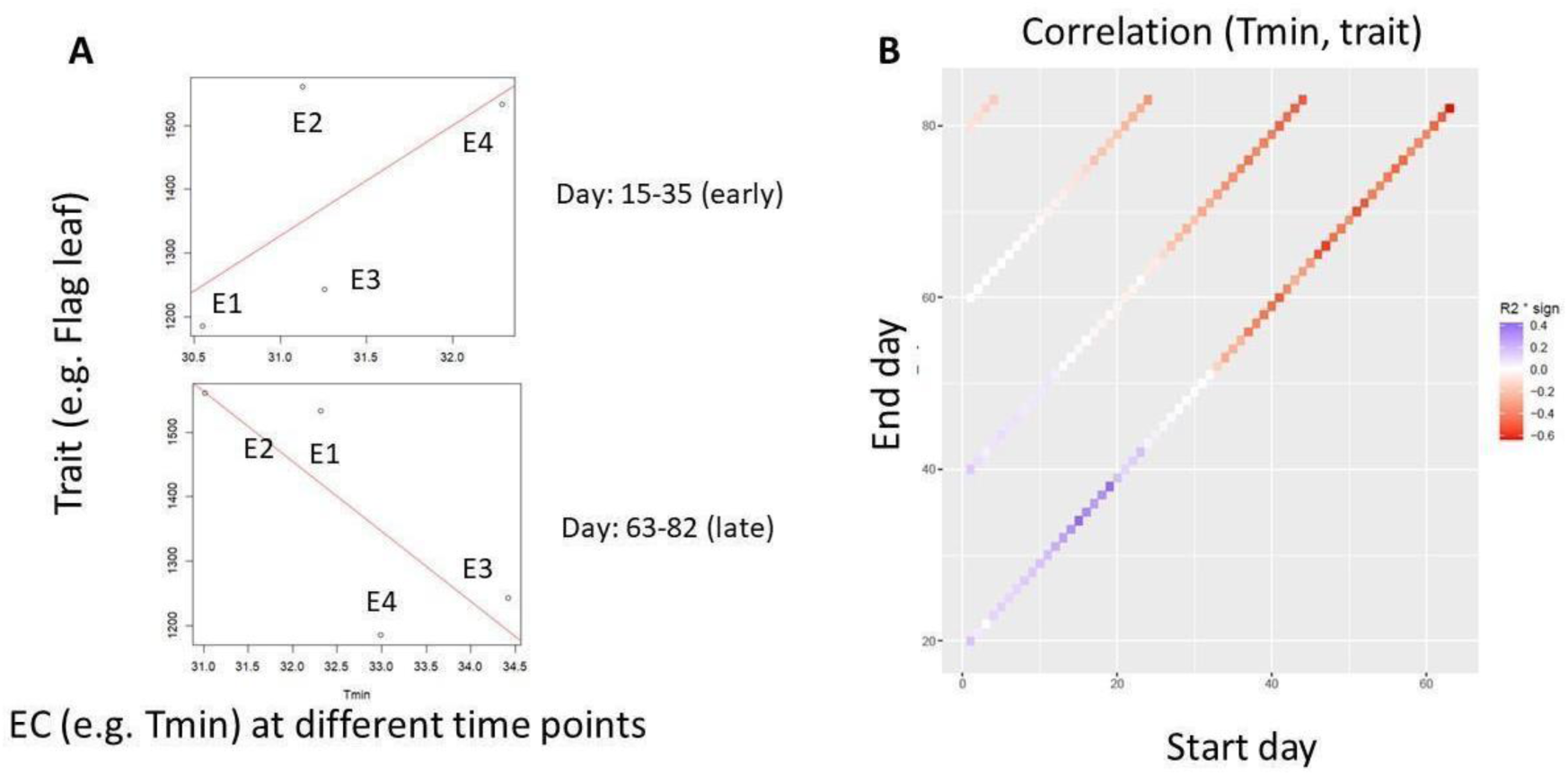

### Method S3

QTL detection procedure and QTL detection threshold

For each combination of population and trait, we performed the following QTL detection procedure:

a. Simple interval mapping (SIM) scan.
b. Selection of cofactors based on the SIM scan profile. Positions with -log10(p-val) larger than the threshold were selected. We selected a maximum of one cofactor per chromosome.
c. Cofactor and QTL detection threshold. The false positive rate for individual cofactors/QTL detection (Type I error) was set to alpha = 0.05. To account for multiple tests, we applied a correction accounting for the number of independent tests, so alpha = 0.05/Meff, where Meff is calculated according to the procedure defined by Li and Ji (2005).
d. Composite interval mapping scan using the selected cofactors
e. The final QTLs were recursively selected per chromosome using the -log10(pval) results of the CIM profile. We first selected the most significant position and then applied an exclusion window of 20 cM around the QTL position. We continued to search for the next most significant position until no more significant positions could be selected.
f. Estimation of the QTL effects. We estimated the QTL effect simultaneously by including all detected QTLs in the estimated model. We also estimated the global R squared of the whole QTL set as well as partial R squared for each final selected QTL position using a linear model. The R squared values were adjusted for the number of degrees of freedom.

QTL detection threshold

| Population | $-\log_{10}(p\text{-val})$ |
| --- | --- |
| GR2012 | 4.647689 |
| GR2013 | 4.483628 |
| KK2012 | 4.181491 |
| KK2013 | 4.187031 |
| Lata | 4.428637 |

### Methods S4

Approximate mixed model computation and QTL test statistic

To reduce the computational power needed to perform the QTL scan we implemented an approximate mixed model computation similar to the generalized least square strategy implemented in Kruijer et al. (2015). The procedure consists of estimating a general VCOV (^*Ṽ*^ ) using model 2 without the tested QTL position, which means estimating the VCOV of model 2 without the QTL term for the SIM scan and the same model with selected cofactors for the CIM scan. The statistical significance of the tested QTL positions and the different allelic effect was obtained by using ^*Ṽ*^ to get the following Wald statistic *W_Q_* = *β^T^V(β)^-1^β*, where *β = (X^T^V^-1^X)^-1^X^T^V^-1^y, V(β) = (X^T^V^-1^X)^-1^*, X represents the fixed effect matrix including the QTL position, and *y* the vector of phenotypic values. *W_Q_* follows a chi-square distribution with degree of freedom equal to the number of tested QTL allelic effects ((*N_par_ – 1) * *N*_env_* for the main QTL term).

### Methods S5

Synonyms of the parental lines’ name

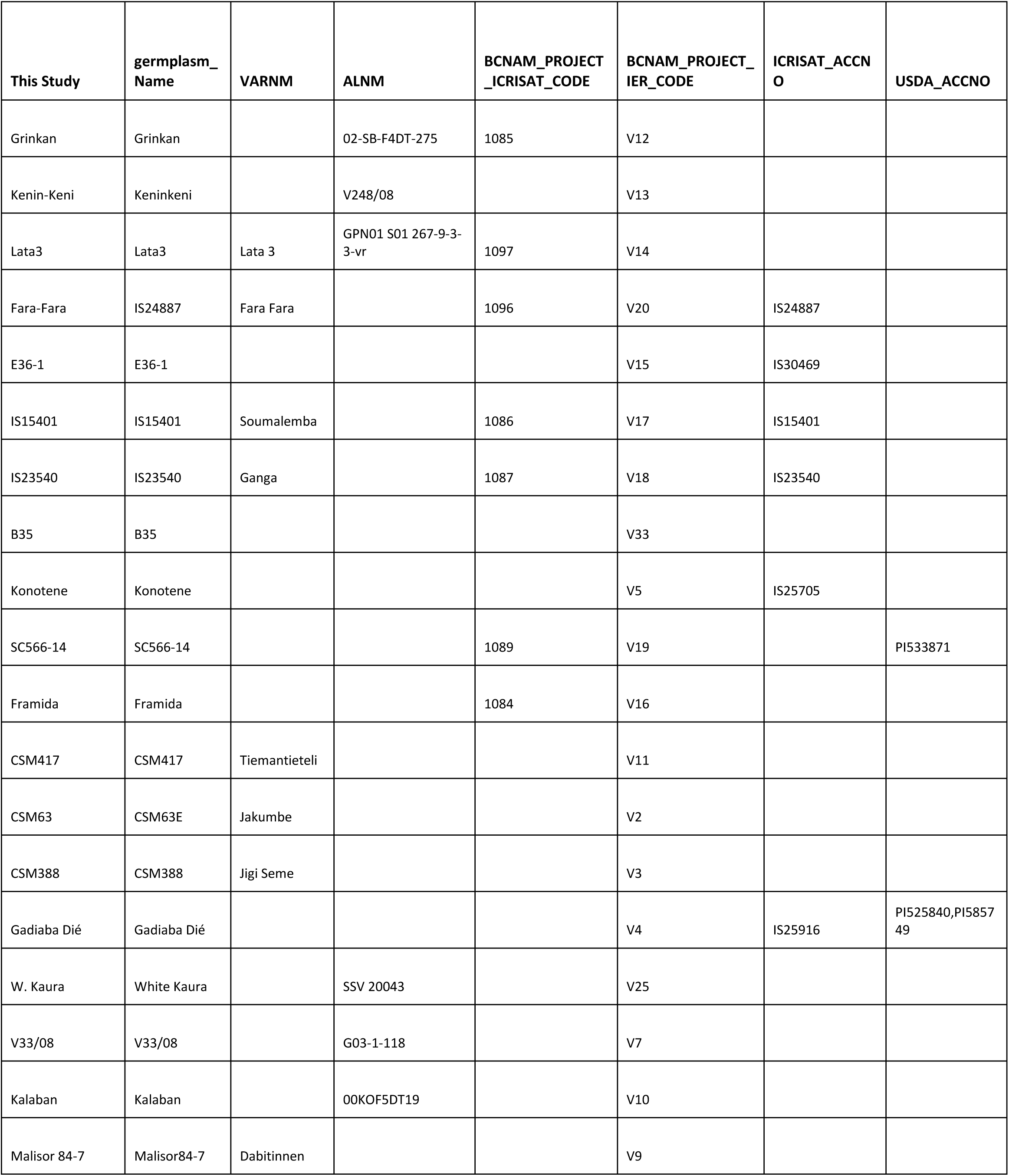
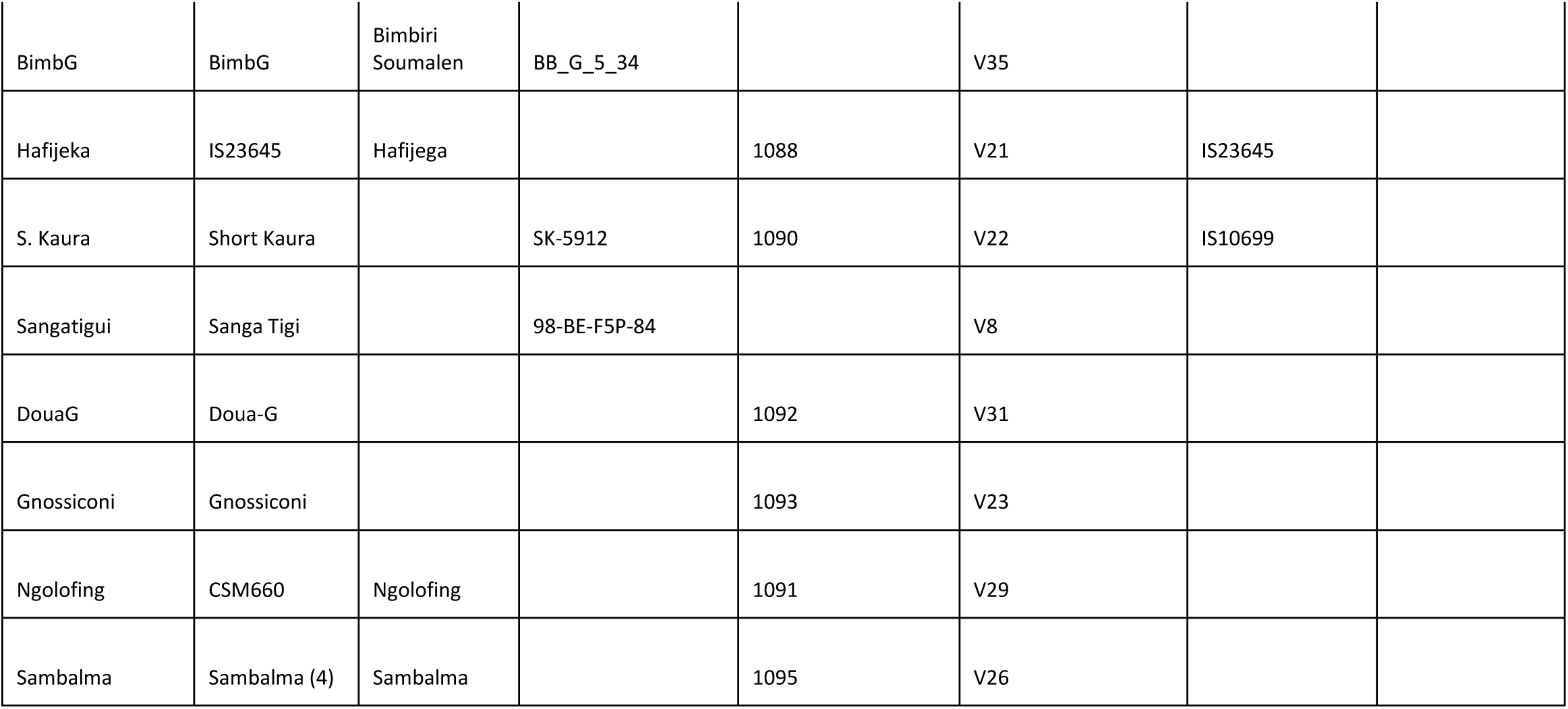

| This Study | germplasm_<br>Name | VARNM | ALNM | BCNAM_PROJECT_<br>ICRISAT_CODE | BCNAM_PROJECT_<br>IER_CODE | ICRISAT_ACCN<br>O | USDA_ACCNO |
| --- | --- | --- | --- | --- | --- | --- | --- |
| Grinkan | Grinkan |  | 02-SB-F4DT-275 | 1085 | V12 |  |  |
| Kenin-Keni | Keninkeni |  | V248/08 |  | V13 |  |  |
| Lata3 | Lata3 | Lata 3 | GPN01 S01 267-9-3-3-vr | 1097 | V14 |  |  |
| Fara-Fara | IS24887 | Fara Fara |  | 1096 | V20 | IS24887 |  |
| E36-1 | E36-1 |  |  |  | V15 | IS30469 |  |
| IS15401 | IS15401 | Soumalembe |  | 1086 | V17 | IS15401 |  |
| IS23540 | IS23540 | Ganga |  | 1087 | V18 | IS23540 |  |
| B35 | B35 |  |  |  | V33 |  |  |
| Konotene | Konotene |  |  |  | V5 | IS25705 |  |
| SC566-14 | SC566-14 |  |  | 1089 | V19 |  | PI533871 |
| Framida | Framida |  |  | 1084 | V16 |  |  |
| CSM417 | CSM417 | Tiemantieteli |  |  | V11 |  |  |
| CSM63 | CSM63E | Jakumbe |  |  | V2 |  |  |
| CSM388 | CSM388 | Jigi Seme |  |  | V3 |  |  |
| Gadiaba Dié | Gadiaba Dié |  |  |  | V4 | IS25916 | PI525840,PI5857<br>49 |
| W. Kaura | White Kaura |  | SSV 20043 |  | V25 |  |  |
| V33/08 | V33/08 |  | G03-1-118 |  | V7 |  |  |
| Kalaban | Kalaban |  | 00KOF5DT19 |  | V10 |  |  |
| Malisor 84-7 | Malisor84-7 | Dabitinnen |  |  | V9 |  |  |
| BimbG | BimbG | Bimbiri<br>Soumalen | BB_G_5_34 |  | V35 |  |  |
| Hafijeka | IS23645 | Hafijega |  | 1088 | V21 | IS23645 |  |
| S. Kaura | Short Kaura |  | SK-5912 | 1090 | V22 | IS10699 |  |
| Sangatigui | Sanga Tigi |  | 98-BE-F5P-84 |  | V8 |  |  |
| DouaG | Doua-G |  |  | 1092 | V31 |  |  |
| Gnossiconi | Gnossiconi |  |  | 1093 | V23 |  |  |
| Ngolofing | CSM660 | Ngolofing |  | 1091 | V29 |  |  |
| Sambalma | Sambalma (4) | Sambalma |  | 1095 | V26 |  |  |

